# “Double-dipping” in testing for phenotypic differences with scRNA-Seq

**DOI:** 10.64898/2026.10.02.756269

**Authors:** Nicolas Sanchez, Lucas Etourneau, Elizabeth Purdom

## Abstract

For the purpose of differential expression (DE) analysis in single-cell RNA-sequencing (scRNA-Seq), phenotype differences between samples are often tested within specific cell types. Cell types are regularly imputed by clustering the same gene expression data which is later used for phenotype testing. This creates the potential for a “double-dipping” or post-selection inference problem resulting in inflated rates of false discoveries. While this selection bias is known to inflate significance in cell-type marker identification, its effect on sample-level phenotype testing, e.g. in patient cohorts, has never been explored despite the growing preponderance of this type of analysis. To address this, we perform an extensive simulation study and demonstrate that naive clustering on uncorrected embeddings can severely inflate the False Discovery Rate (FDR) in the presence of strong phenotypic differences. However, we further show that applying batch-correction methods to remove phenotypic effects prior to clustering resolves the FDR inflation with no obvious loss of power. Finally, we provide measures of phenotypic imbalance that can be applied to real datasets which closely track the false discovery proportion and thus can be used to as part of data exploration to gauge the risk of post-selection inflation of p-values in a particular dataset.

## 1 Introduction

In gene expression studies, it is common to perform hypothesis testing on each gene to evaluate whether there are differences in the amount of gene expression between different conditions, often called differential expression (DE) analysis. In single-cell mRNA-sequencing (scRNA-Seq), the hypothesis tests performed are also frequently limited to cells that are in specific cell-types. The assignment of cells to cell-types is commonly done by clustering of the cells based on their joint gene expression data. This creates a setting of “double-dipping” of the data, also called post-selection inference, where the same data is used both to select the set of cells to test as well as to perform the hypothesis test. Post-selection inference can result in invalid p-values whose significance is inflated. This inflation has been demonstrated in cell-type markers discovery, for example, to substantially over-estimated the significance cell-type markers ([32],[24]).

However, DE analysis with scRNA-Seq data is not confined to identifying cell-type markers. With larger scRNA-Seq studies, it is common to design studies across many samples, often patient cohorts, which collect single cells from each sample. In this type of study, a common DE analysis is the testing for gene expression differences between different *sample-level* phenotypes, such as the disease outcome in a patient cohort or comparisons of perturbations between samples. In this case, the hypothesis that is tested is whether there is a differential mean gene expression level between the cells from different phenotypes *within* a particular cell-type [6]. Such tests still depend on selecting the cell-type from the data and thus result in hypothesis testing following selection. But the relationship of the selection to the hypothesis test is quite different from that of identifying cell-type markers. The gene expression data is still used to determine the assignment of cells to cell-types, but the sample phenotype information being tested is *not* directly used to assign cells to cell-type. As a result, it is unclear the effect of such selection on the DE tests, despite the ubiquity of such DE procedures. Developing a more precise understanding of the inferential consequences of post-selection inference as well as strong guidelines for mitigating its effect are crucial since multi-cohort, cluster-based differential expression results are frequently used to identify potential disease mechanisms and drug targets. We perform the first extensive evaluation of the performance of such phenotype tests via simulation, benchmarking the effect of the clustering of cells on control of the false discovery rate (FDR). We demonstrate that the clustering of the cells can adversely affect the control of FDR when large enough levels of differential expression are present to impact the clustering of cells. We further show that common methods of batch-correction prior to clustering used to remove sample variation can create phenotype-independent cell-type clusters that allow for effective FDR control without loss of power. We provide measures of phenotypic imbalance that can be applied to the found clusters to predict whether the clustering is likely to lead to inflated false discovery proportions. We also show that while high resolution clustering strategies (i.e. those which result in many small clusters) can help in accurately detecting regions of phenotypic differences, that such strategies are sensitive to sample fluctuation, even after batch correction methods, and as a result demonstrate highly inflated levels of false discovery. In conclusion, our results show that so long as the clusters are reasonably sized, cluster-based hypothesis testing for phenotypic differences can be safely performed despite the double-use of the data when paired with batch correction methods.

## 2 Results

In this work, we evaluate (via simulation) the effect of using the same set of gene-expression data to both find putative cell-types by clustering of the data and to test for differences between sample-level phenotypes within these found clusters. We refer to this as a “cluster-based” DE strategy, meaning that the results of the clustering define the hypotheses being tested. This results is a form of post-selection inference, since the set of hypotheses being tested is selected based on the data.

How are the results affected by the fact that assignment of cells to clusters is determined from the data? It has been shown when testing for differences in the mean expression *between* different clusters (“cell-type markers”) that there is severe increase in type I error [32, 24]. In this case, there is a clear correlation between the mean gene expression of cells in a cluster and the cell’s cluster assignment. But in this work we consider a different setting, namely that of comparing the difference of gene expression between phenotypes *within* a cluster, which is a common analysis of patient disease cohorts. In this case, since the phenotype assignment of a cell is known *a priori*, the effect of using the data to assign cells to cluster-based hypothesis is less obvious. In particular, if there are no phenotype-based differential expression in the gene expression, then the phenotype is independent of the observed data and thus also of the assignment of cells to clusters. In this case there should be no inflation in the p-values due to the selection step.

However, if some genes have different expression levels between the two phenotypes in some cells, then these DE cells may influence the clustering via their impact on the low-dimensional embeddings in which the clustering is performed. For example, genes with large amounts of DE may affect the low-dimensional embeddings; PCA, based on decomposing the variance, is known to not be robust to differences in small numbers of observations [5] This could result in cell embeddings that are correlated with the cell’s phenotypic value, which can result in assignments of cells to clusters which are correlated with the phenotype. Figure 1 provides a cartoon illustration of this possibility.

**Figure 1:**
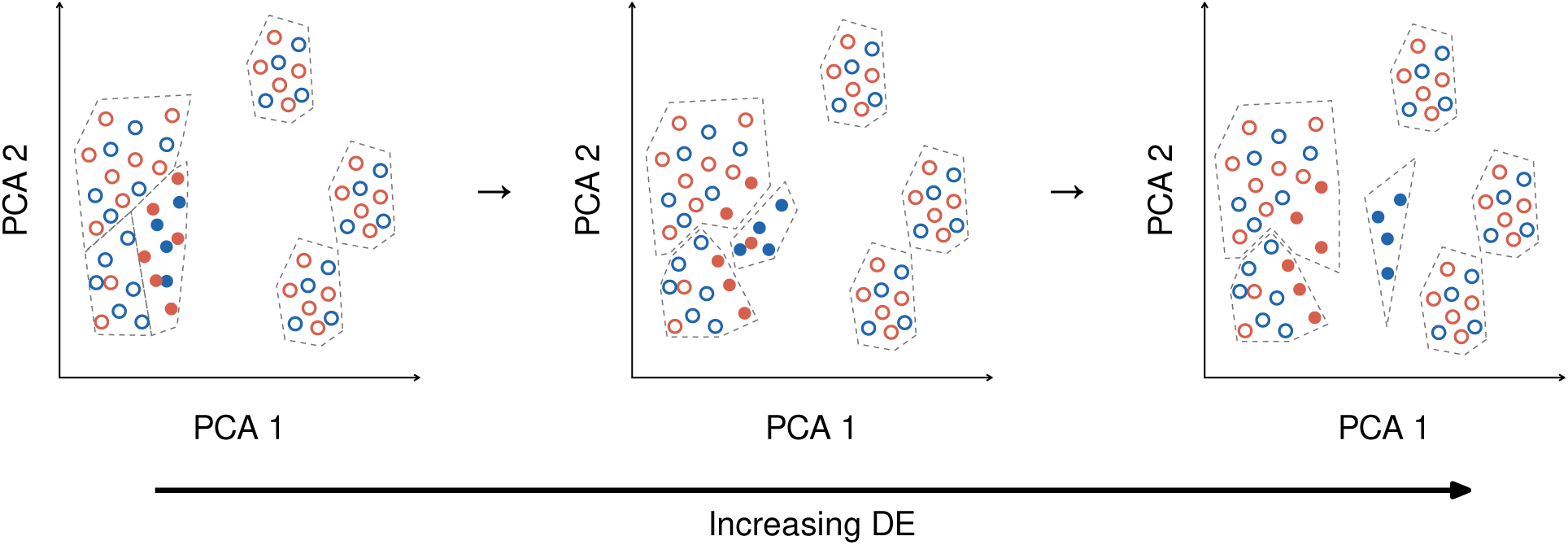
Potential Effect on Embedding and Clustering with Increasing DE. Shown is an illustration of potential clustering failures when increased differential expression results in latent space embeddings that isolate cells from their natural comparison cells. Points with an filled circle are in the true DE neighborhood *N_g_* for a particular gene (for simplicity, the sample of origin of the cells are not indicated, unlike in Figure 2a)). In the first panel, there is no DE effect, and cells from the DE neighborhood are clustered together according to their true shared celltype; with increasing difference between phenotypes in *N_g_*, the latent embeddings show separation in these cells and cells from the different phenotypes of *N_g_* are increasingly clustered into separate clusters. This results in a cluster of phenotype A cells without the “correct” set of phenotype B cells for comparison. Moreover, cells from phenotype B from the DE neighborhood *N_g_* are incorrectly assigned with cells from other celltypes, potentially causing a false DE signal for this gene in those clusters.

To investigate the effect of post-selection inference in this setting, we perform an extensive simulation study where we start with data from a scRNA-Seq study of colorectal tumor [22] and select twenty normal samples from the study. We then randomly assign *samples* to one of two phenotypes, as well as inserting variable numbers of known DE genes. Known DE genes are given simulated gene expression which is the same between conditions, except for the subset of cells that we select to show differential expression. We call the set of cells that show differential expression *N_g_*, and they consist of cells from both phenotypes. We use two different simulation strategies to determine *N_g_*: one where *N_g_* are the same as the underlying cell-types detected in the data (the “aligned” simulation scenario), and one where they vary from the known cell-types (the “non-aligned” simulation scenario). We note that in the absence of DE genes, the true DE regions, *N_g_*, and the clusters found on the data will be essentially the same, offering a baseline as to the performance absent the effect of clustering variation, where the found clusters and the true DE regions are equivalent. More details of the simulation can be found in the Methods, Section 3.2). After randomly assigning samples phenotypes and simulating the DE genes, we perform the standard cluster-based DE pipeline on the simulated data (Figure 2): 1) we perform PCA on the simulated data; 2) we cluster the PCA components to identify clusters of cells, which we will refer to as *Ĉ_k_*; 3) within each sample, we pseudobulk the cells for each cluster and 4) perform DE testing between the two phenotypes. This pipeline is described more fully in the Methods, Section 3.1)

**Figure 2:**
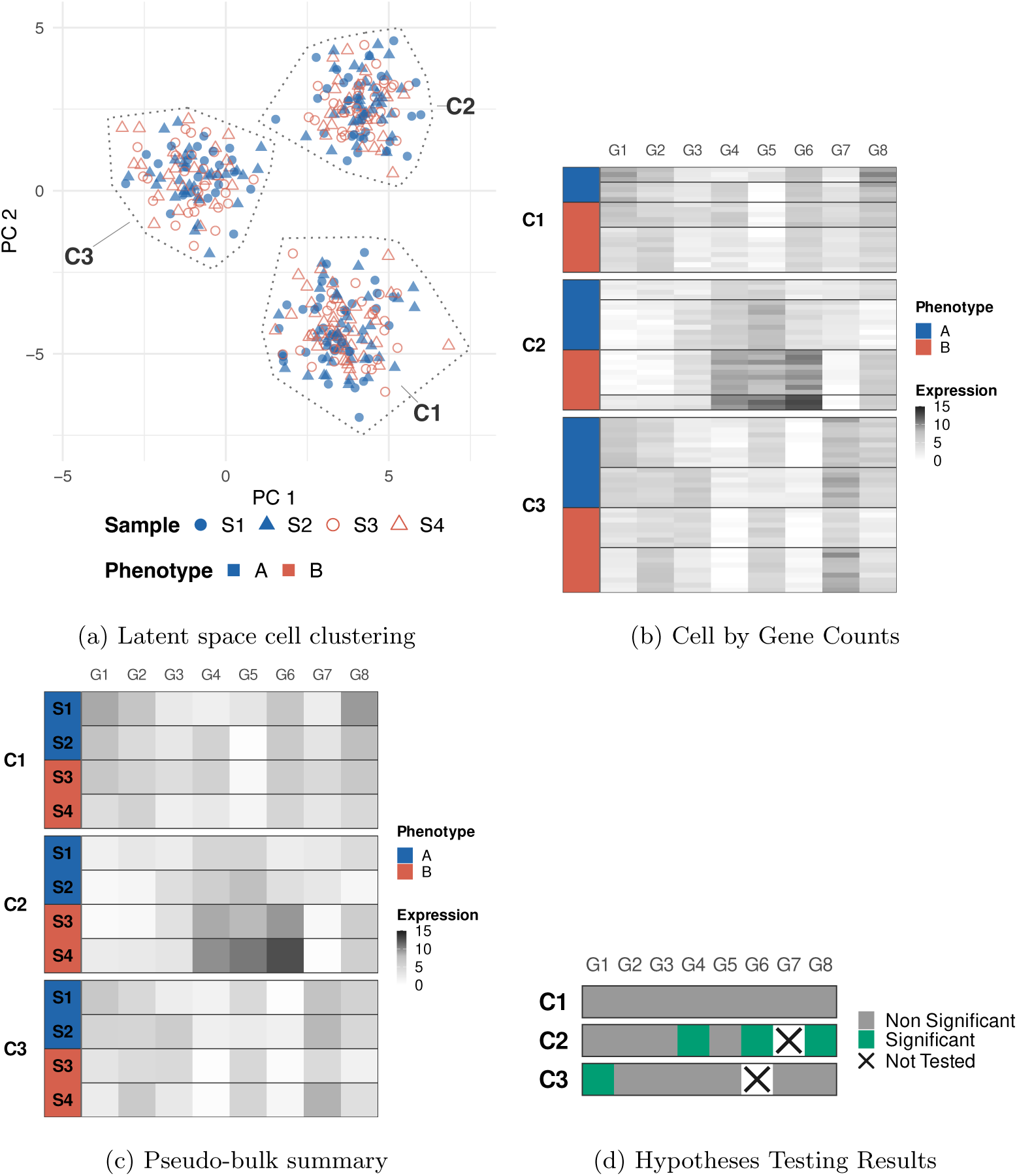
Illustration of Cluster-based DE Analysis Pipeline. (a) Plots of individual cells represented in a lower-dimensional latent space, colored by phenotype and with point shape based on their sample of origin. Latent dimensions are usually estimated irrespective of sample. Individual cells are clustered in the latent space, as indicated by the dotted polygons for each cluster, C1-C3; these clusters form the putative cell-types for downstream analysis. (b) Illustration of the gene count data for individual cells and the relevant known (or estimated) characteristics. Each cell is assigned one of the putative cell-types (represented as distinct blocks of rows and labeled C1-C3) based on the clustering in the latent space of (a). Cells are also known to come from different samples (delineated with thicker horizontal lines), each with their own phenotype (shown by color of leftmost column). (c) Illustration of pseudobulking of individual cell data. Within each cluster, individual cell measurements from (b) are summarized into a single measurement per sample for each gene. (d) Illustration of the results of hypothesis testing. Within each cluster, a hypothesis test is run on the pseudobulk data from (c) for each gene evaluating differences between samples from different phenotypes, unless the hypothesis dropped due to gene-level or cluster level filtering (see Supplementary Material, Text S2.2). All images shown here are for illustration purposes and do not reflect the results of any actual data processing.

### 2.1 False Discovery Rate

We first note that even in the absence of DE genes, this dataset shows slight elevation of the FDR under random assignment of phenotypes; this behavior has been reported in other studies for a variety of testing procedures, and is likely due to deviations of the p-value distribution from the null expectation [14, 25]. We note that this elevation is in the absence of any potential correlation between cluster assignment and phenotype, and so should not reflect post-selection bias. When we synthetically add DE genes, we then see that increasing levels of differential expression results in inflated rates of false discoveries, demonstrating that there is potential for post-selection inference problem, as hypothesized. We see that with large log-fold change (on average 3.0) and large number number of DE genes, there is a clear lack of control (Figures 3a): starting around 250 DE genes (roughly 1.5-2% of all hypotheses tested) the FDR starts increasing, ultimately reaching a FDR as high as 0.64 with 750 genes (roughly 4-6% of all hypotheses tested, Figure S-3). For lower overall log-fold changes (on average 1.5) or low numbers of DE genes, however, we see reasonable control of the FDR relative to baseline (Figure S-4). For comparison, in four real datasets we considered, the proportion of individual cluster hypotheses found significant were almost all in a similar range where FDR problems are seen, between 4-16% (Figure S-5), but the mean observed absolute log-fold changes in genes found significant ranged from 0.8-1.4 (Figure S-6). These summaries initially suggest that real datasets fall on the lower levels of signal and may be safe from these concerns about FDR control. However, we note that when we consider the simulation setting where DE regions are not aligned with found clusters, the effective log-fold change of the *found* clusters in the simulation (rather than the original *N_g_* neighborhoods) drops as a result and is in practice more varied, from 1.2-1.7 (Figure S-6); yet, this simulation demonstrates similar problems with FDR control setting described above (Figure S-4), suggesting that the levels of DE signal needed to demonstrate loss of FDR control are potentially encountered in real data settings.

**Figure 3:**
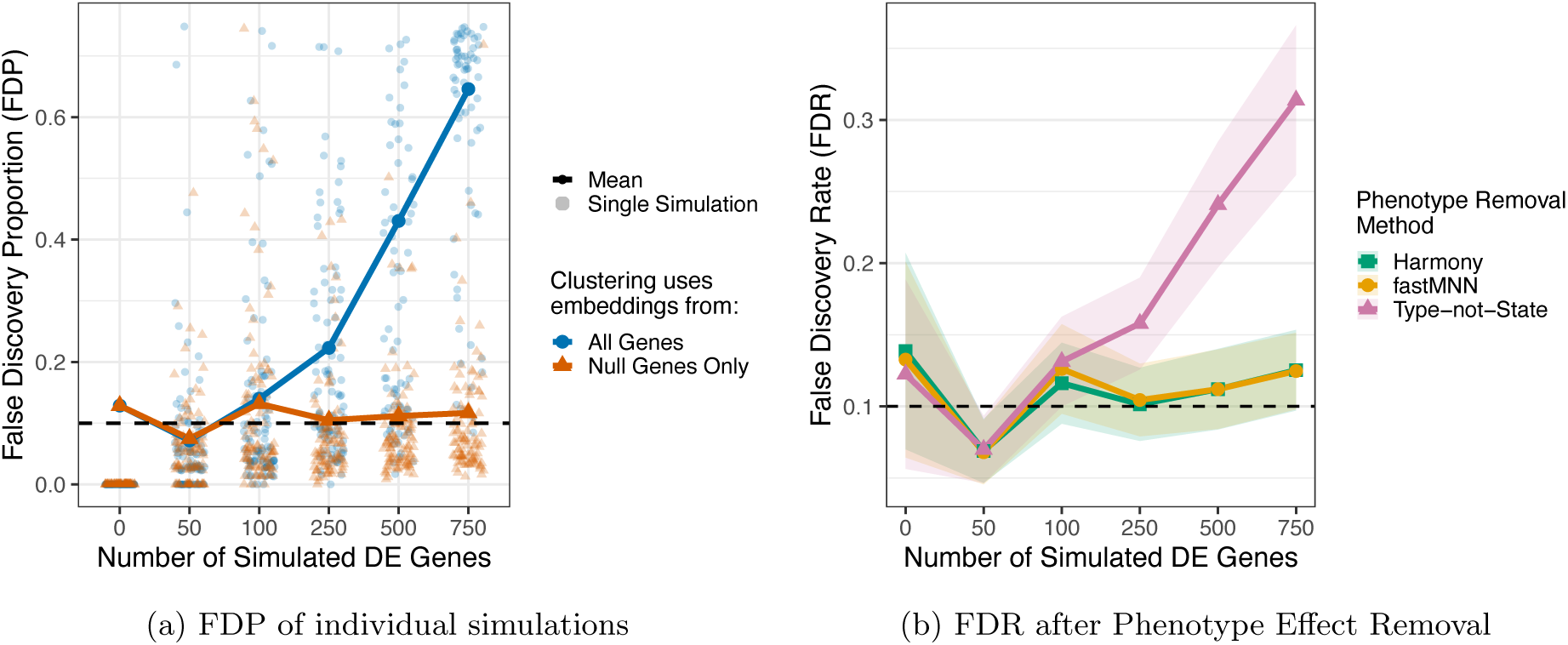
FDR performance of clustering followed by DE. (a) Plot of the the false discovery proportion (FDP) as a function of the number of DE genes when clusters were defined based on only null genes (red) versus on all genes (blue). Each transparent point represents the FDP for a single simulated experiment with 5000 null genes based on BH corrected p-values at nominal FDR cutoff of 0.1, and the number of differentially expressed genes are indicated by the x-axis, jittered for overplotting. The average (FDR) across simulations is shown as solid line. (b) Plot of FDR as a function of the number of DE genes when the clustering of cells is performed after one of three phenotype removal methods (see Section 3.3). For both plots, the simulations are with DE regions defined to be aligned with celltypes, from Harmony-corrected embeddings of the null genes, and with an average log-fold change of 3 (see Section 3.2). The dashed black line indicates the chosen FDR threshold of 0.1. The pattern remains similar for the simulations with non-aligned DE neighborhoods (Figure S-4). See also Figure S-4 for lower levels of log-fold change.

Typical single-cell pipelines for datasets with multiple samples generally include a batch effect removal step to account for possible differences between different samples. The result of such integration procedures is generally new sample-corrected embeddings on which the clustering can be performed. In the context of cluster-based DE, these batch removal methods have also been proposed to avoid the potential of phenotype differences in defining cell-types ([31], [12], [4]). Removing sample effects has the advantage of removing both potential sample or batch artifacts as well as systematic phenotypic differences. We note that the actual hypothesis testing of the resulting found clusters acts on pseudobulking the original data and makes no use of these sample-integrated embeddings; the integrated embeddings created by sample-integration affects solely the cluster assignments of the cells.

We consider the effect of two popular strategies of batch correction: Harmony [12] and fastMNN [7]. We also consider clusterings based on embeddings which use as input only the top 5000 “Type not State” genes (inspired by [31], see Supplementary Text, Section 3.3). In our simulations, both Harmony and fastMNN corrections before clustering result in control of the FDR across all different numbers of genes and log-fold changes (Figures 3b and S-10). This FDR control remains true regardless of the choice in the simulation of which embedding to use in defining the true *N_g_* (Figure S-11), meaning there is not a bias in the simulation based on whether the same embedding method were used in defining *N_g_* and in finding the empirical clusters. Clustering based on “Type not state” genes as input does not result in control of the FDR when there is large DE signal (FDR ranges from 0.27-0.35), though it is still reduced compared no selection or correction applied.

Naively, we could assume that the batch-correction strategies are effective at controlling the FDR because they lead to clusters that are, in effect, the same as clustering the null genes. However, this is not the case. Different choices of clustering strategy and embedding corrections lead to quite different definitions of *N_g_*and of found clusters. In fact, significant difference in clustering occur even for embeddings on the null genes only (Figure S-1). Yet, we see similar levels of FDR control so long as we correct for phenotype using Harmony or fastMNN, even when their similarity to the true DE neighborhoods varies. More specifically, we can see that the correspondence of a simulation’s found clusters to the oracle clusters, as measured by the Adjusted Rand Index (ARI), is not strongly predictive of its FDP (Figure 4a). Rather, we find that the important characteristic for achieving low FDP is having empirical clusters that are statistically less dependent on phenotype. If we consider the adjusted mutual information (AMI) statistic between the assignment of cluster and the phenotype of cells, we see strong correlation with the AMI and the proportion of false discoveries in an individual simulation (FDP) (Figure 4b, S-9). Examining the raw p-values of the null hypotheses, we directly see that p-values from hypotheses with large cluster imbalances are negatively skewed, resulting in inflated significance (Figures S-12). Here cluster imbalance is measured using the residual size-weighted Pearson sample residual (see Section 3.5.1) which measures imbalance for individual clusters within a dataset.

**Figure 4:**
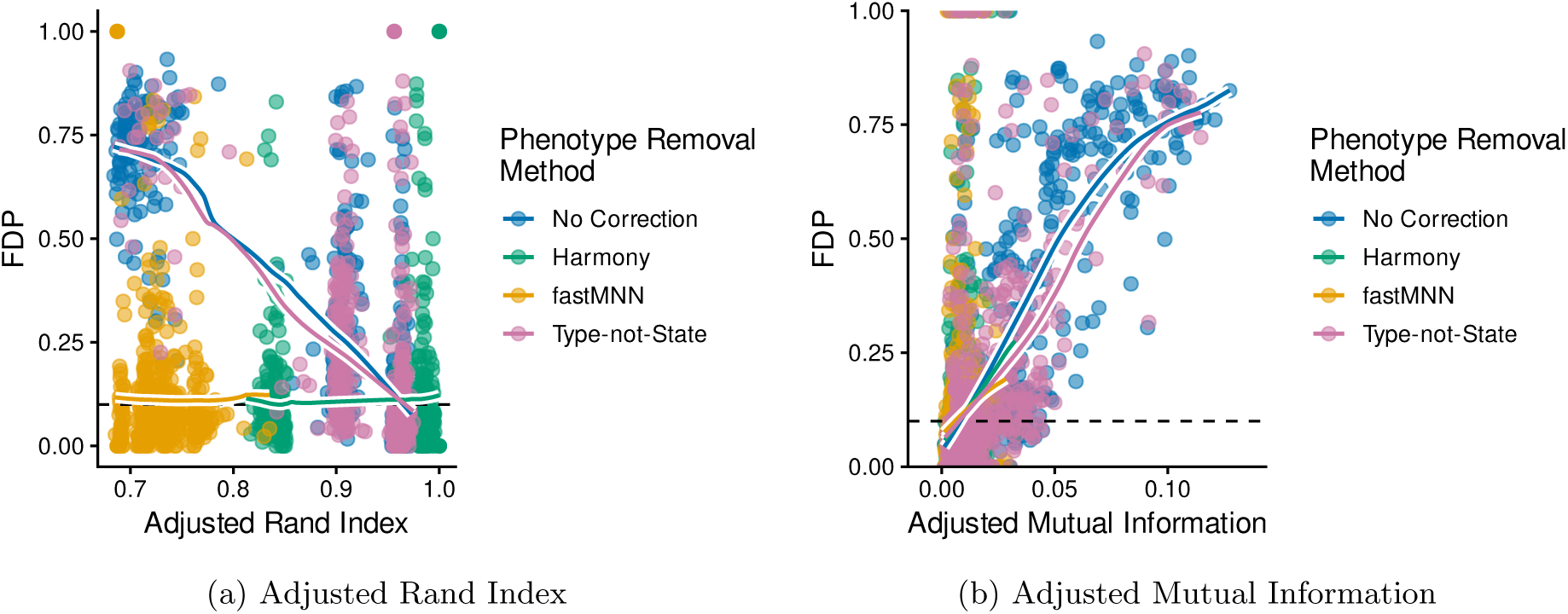
Metrics for potential phenotype bias detection. In both plots, each point shows the false discovery proportion (FDP) for a single simulation (y-axis) plotted against: (a) the Adjusted Rand Index (ARI) between the simulation’s found clusters and the oracle cell-types used to define *N_g_* and (b) the Adjusted Mutual Information (AMI) between found cluster and phenotype labels for all clusters tested for DE. ARI measures how well the empirical clusters correspond to the ground-truth differential neighborhoods *N_g_*, while the AMI measures how much empirical clusters show imbalance between phenotypic differences. Simulations shown here were run with celltype-aligned DE regions and Harmony corrected embeddings of null genes with an average true log-fold change of 3. Plotted are the results across all simulations, regardless of the number of DE genes. Colored trend lines, intended as visual summaries only, are local linear smooths computed for the subset of hypotheses with the indicated phenotype removal method. See Figure S-9 for the same metrics when the simulations have DE neighborhoods unaligned with celltype.

The AMI imbalance metric does not require knowledge of the ground truth and can be calculated directly on observed data values. This suggests that AMI could be effective in real data applications for measuring whether there has been sufficient removal of the phenotype effect in the embedding. We apply the AMI to four datasets, shown in Table 1. Certain datasets exhibit concerning levels of phenotype imbalance when compared to our simulation findings in 4. Both the Pelka [22] dataset, containing MMR-deficient tumor versus adjacent normal colorectal tissue, and the Kang [10] dataset, comparing untreated control and 6-hour IFN-*ω* stimulation, exhibit strong adjusted mutual information values (0.142 and 0.103 respectively). Without phenotype correction a large portion of clusters go completely untested in these two datasets (7 out of 19 for Pelka and 6 out of 11 for Kang), further illustrating their imbalance. Meanwhile, the Liu dataset [15], containing psoriasis vulgaris versus healthy skin samples, and the Melms dataset [20], comparing pre-pandemic controls to COVID-19 decedents, display more muted AMI values. After correcting for phenotype via Harmony or fastMNN, AMI drops for the Pelka, Kang and Liu datasets, leading to all datasets having AMI in ranges with better observed FDR in our simulations (*<*0.05).

**Table 1:** Phenotype Imbalance Measures for Real Datasets. Measures of Adjusted Mutual Information (AMI) for four actual datasets. The datasets and pre-processing steps are described in Supplementary Text, Section S4; note that a subset of normal samples from [22] formed the basis of our simulations.

| Dataset | Phenotype Removal / Embedding Method |  |  |  |
| --- | --- | --- | --- | --- |
|  | None | Harmony | fastMNN | TvsS |
| **Pelka et al. (2021)** [22] |  |  |  |  |
| MMRd vs. Normal | 0.142 | 0.046 | 0.066 | 0.109 |
| **Kang et al. (2018)** [10] |  |  |  |  |
| Stimulated vs. Control | 0.109 | < 0.001 | < 0.001 | < 0.001 |
| **Liu et al. (2022)** [15] |  |  |  |  |
| Normal vs Psoriasis | 0.054 | 0.023 | 0.027 | 0.08 |
| **Melms et al. (2021)** [20] |  |  |  |  |
| Control vs. COVID-19 | 0.013 | 0.029 | 0.030 | 0.027 |

### 2.2 Power

Methods which explicitly remove the dependence between cluster assignments and phenotype can greatly improve the control of FDR. However, as mentioned, by integrating the samples regardless of phenotype, they have the potential to remove biological signal. We now consider whether this control comes at a loss of power in finding DE signal? Two aspects of the simulation are important to consider in considering our ability to detect DE regions: the standard pre-filtering of hypotheses before testing and the definition of which found clusters are considered DE.

Filtering of hypotheses before testing is ubiquitous in genomic studies, and, in the case of independent filtering, has been shown to boost overall power by removing tests unlikely to have power to reject the null and avoiding unnecessary multiple testing burden [3]. However, unlike most genomic studies, the standard filtering implemented in the cluster-based DE pipeline directly uses the phenotypic information of samples, removing clusters with insufficient representation of either phenotype in order to be able to run the tests; this is in addition to filtering of low expression genes (see Supplementary Material, Text S2.2). Furthermore, the different phenotype removal correction methods result in different clusters, *Ĉ_k_*, which in turn have differences in both gene expression levels and sample imbalance. Thus, the proportion of filtered hypotheses will change across the different methods we are comparing, also unusual in method comparisons in genomic studies. In our simulation, we see that the proportion of non-null hypotheses that are filtered grows with stronger phenotypic differences, while staying comparatively stable if they are corrected via Harmony or fastMNN (Figure 5a). Specifically, more clusters are removed due to poor sample representation when clusters are defined based on the uncorrected gene embeddings (Figure S-14). Since removing a cluster results in all gene hypotheses for the cluster being removed, this results in a large increase in filtering of hypotheses (Figures S-15 and S-16).

**Figure 5:**
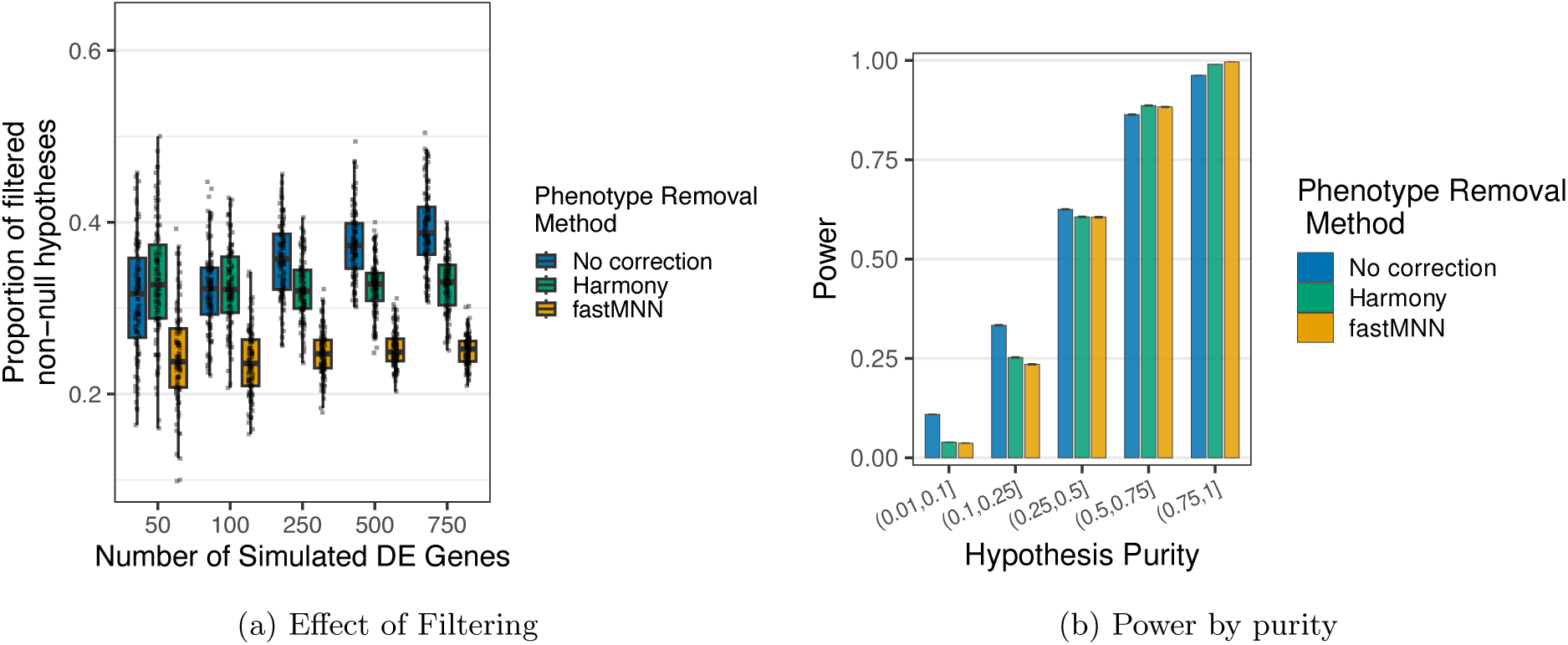
Effects of Phenotype Correction Methods on Filtering and Power. (a) shows the effect of filtering of hypotheses during the DE testing on the power. The y-axis represents the the proportion of non-null hypotheses in an experiment that are dropped due to filtering and are never tested. The distributions of such proportions for each phenotype removal method (indicated by the color) are displayed as box-plots, stratified by the number of DE simulated genes. This plot shows results for the celltype-aligned strategy for choosing DE neighborhoods, on Harmony embeddings. Trends are similar for other *N_g_* choices (see Figure S-15). (b) each bar shows the average power (y-axis), pooled across both the DE-aligned and unaligned strategies for choosing *N_g_*, for hypotheses with specific purity levels (x-axis) for different phenotype removal methods (different colors, see legend).

The second consideration in our simulation is how to determine which *H*_0*,g,k*_ are true, i.e. which *Ĉ_k_* are considered DE for a particular gene. The found *Ĉ_k_* are random will not exactly coincide with a *N_g_* neighborhood; the proportion of the cells of *Ĉ_k_* that overlap *N_g_* we will refer to as the purity of the cluster. Even in simulations where the true DE neighborhoods *N_g_* are designed to align with one of the detected clusters (see Section 3.2), there are often small amounts of overlap of the true *N_g_* with other clusters, leading to a large portion of hypotheses with low purity (see Supplementary Figure S-17). Such low purity is unlikely to allow for detection. Indeed, if we define clusters with *any* DE cells as DE for our power calculations, the average power among tested hypotheses drops to lower than 0.25 (Figure S-18). A more reasonable requirement is to only consider in our power calculations those *H*_0*,g,k*_ with purity greater than a certain minimal amount (see Supplementary Text, Section S2 for details). If we require at least 0.01 purity, the power based on found clusters still drops compared to the oracle that uses only the true DE cells, but less precipitously, to 0.55-0.95, depending on the phenotype removal method (Figure S-19a). In the power discussion that follows, we will require a minimal purity of 0.01 for a *Ĉ_k_* to be considered DE.

More generally, the purity of the cluster is the main driver for differences in power throughout our simulations, and choices in our simulation settings create quite different purity distributions. For example, the DE-aligned simulation setup, as expected, has high purity for most hypotheses, compared to setup that is not DE-aligned and demonstrates more variablity (Figure S-20). This results in clearly higher power overall for the DE-aligned simulation, regardless of any other choices in the simulation or analysis (Figure S-21). Similarly, the relationship between between power and the number of DE genes differs between the DE-aligned and non-aligned simulation settings, with the aligned simulations showing little effect, while the non-aligned show increasing power with increasing number of DE genes. This disparity can be tied to the fact that the high overlap seen in the DE-aligned simulations remains quite stable regardless of the number of DE genes (Figure S-22), while in the non-aligned simulations, the found clusters show an increase of overlap with increasing numbers of DE genes (from Figure S-23).

The predominate role of purity in predicting power makes comparison of the power of different phenotype removal methods difficult. The choice of embedding space for determining *N_g_* in our simulation setup interacts with the phenotype removal method to create different distributions of purity in different settings, particularly in the DE-aligned simulation setting. There, we see a strong relationship between the power of a phenotype removal method and whether that same method was used as the basis for the embedding space to determine *N_g_*; in the non-aligned simulation setting, where the *N_g_* differ more strongly from the found clusters regardless of embedding, this difference is not pronounced (Figure S-21). Once we condition on hypotheses with similar overlap, we see that there is little difference in power between the phenotype removal methods (Figure 5b), meaning that the only conclusion is that the phenotype method that creates clusters that best align with the true DE regions will have the greatest power – something that is impossible to evaluate on real data. However, our simulations do not indicate that phenotype methods are significantly removing the biological signal and are detrimental to power, though obviously this will depend on the dataset and the design of the dataset.

### 2.3 Cluster Resolution

All of the previous analyses use a default resolution of clustering that provides roughly 11-15 clusters (a 0.1 resolution parameter in the Leiden clustering algorithm). This is a standard choice that roughly matches the number of clusters determined by the authors. However, based on the previous power discussion, it is natural to ask whether smaller clusters might be more sensitive to finding the right set of cells for obtaining high overlap with true DE regions. On the other hand, smaller clusters might also be more susceptible to isolating particular samples or DE cells, creating more phenotype imbalance and risk of false discoveries.

To explore this tradeoff, we vary the Leiden resolution parameter (from 0.1 to 4), resulting in a range of around 10 clusters to roughly 75 found clusters per simulation (Figure S-25). Our simulations show diverging behavior between the extremes of these values. Higher resolution leads to larger levels of false discoveries, even *without* the presence of DE genes (Figures 6a, b, and S-27). However, unlike the low resolution we have considered so far, the poor performance of higher resolution clusters does not grow with the increasing presence of DE. Phenotype correction methods consistently improve false-discovery levels, but have a far larger beneficial impact on low-resolution cluster analysis than that of high-resolutions: in the low resolution setting, the phenotype correction largely resolves the false discovery problems, while high resolution simulations retain noticeably inflated levels of false discoveries.

**Figure 6:**
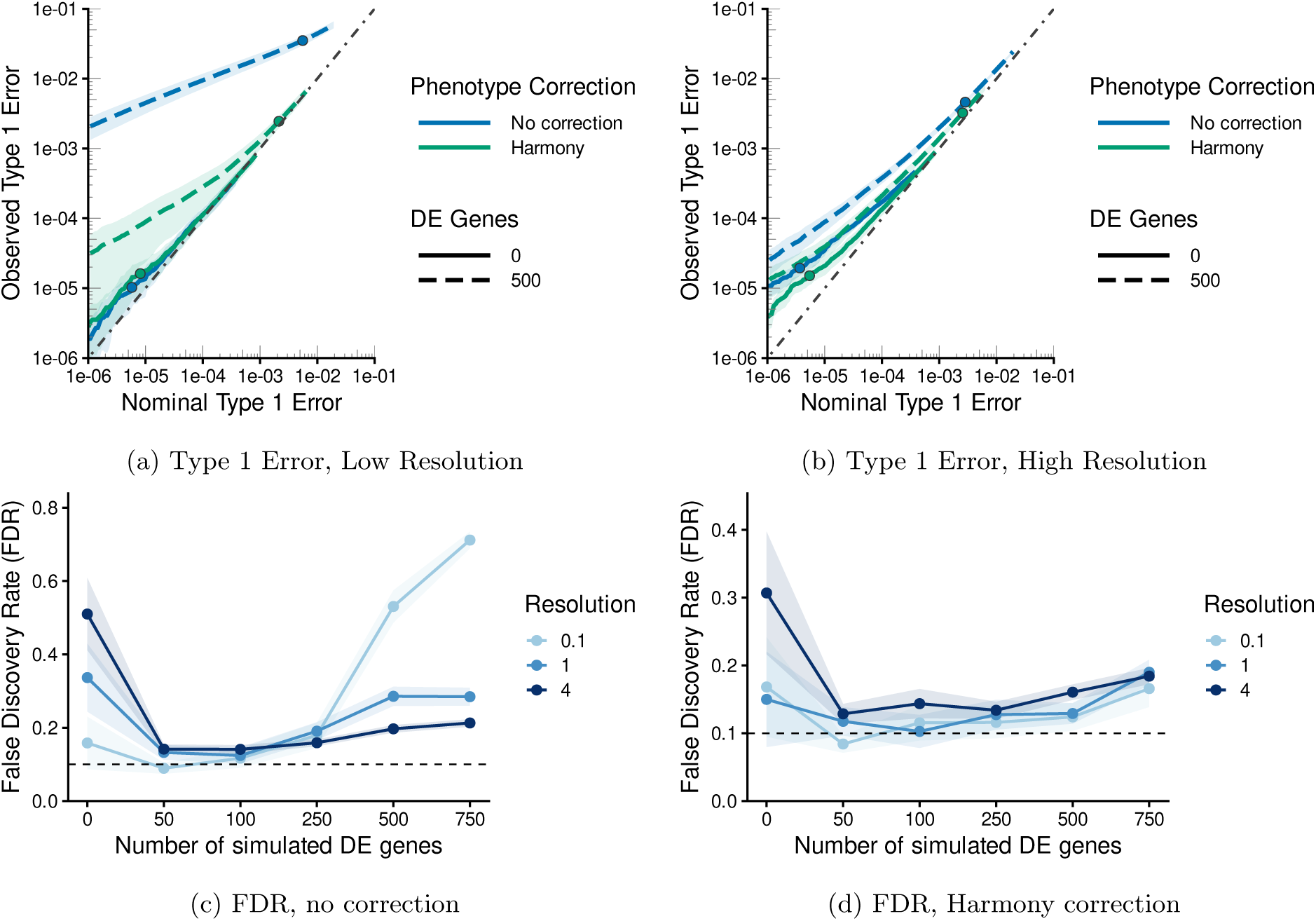
Loss of FDR and Type 1 Error Control with varying resolutions and number of DE genes. The top row compares with observed type 1 error rates (y-axis) to nominal type 1 error rates (x-axis) when using raw, uncorrected p-values for either no DE genes or 500 DE genes (shown by line type) and with a given phenotype correction method (shown by color). (a) shows results for the low resolution clustering of the simulated data (Leiden resolution parameter of 0.1) while (b) shows results for high resolution clustering of the simulated data (Leiden resolution parameter of 4). The bottom row shows the observed FDR (y-axis) using a nominal FDR cutoff of 0.1 (horizontal dotted line) applied to BH corrected p-values, plotted against increasing number of DE genes (x-axis). Different resolutions used for clustering of simulated data are shown as differently colored lines. (c) shows the results when there are no corrections to phenotype effect applied to the simulated data before clustering while (d) shows results when Harmony correction at the sample level is applied before clustering. All plots were run on the simulation setting with DE neighborhoods are not aligned to celltype and use the Harmony sample-corrected null genes for the embedding.

We would note that more detailed examination of trends shows differences between FDR and Type I error as measures of false discovery. This is due to the fact that FDR correction methods, like Benjamini-Hochberg, involve the joint distribution of both true positives and false positives. Thus the insertion of simulated DE genes changes the p-value distribution which has complicated effects on the FDR. In particular, we see a drop in FDR once DE genes are introduced at all resolutions, but particularly in high resolutions (Figure 6c,and S-30c); this is due to having introduced true positives with large signal creating a p-value distribution which is highly separated from the distribution of that of the null hypotheses (Figure S-31), see discussion in Supplementary Section S3. Real data would not demonstrate this kind of stark differentiation. Indeed with lower log-fold change in our simulated DE genes, we see a slower decline in the FDR for low resolution (Figure S-32), with the lack of FDR control continuing across the range of DE genes.

While high resolution clustering has greater issues with false discoveries, it also shows higher power than lower resolution even when considering ROC curves which calibrate the p-values to account for varying levels of false discovery rates (Figures 7a, b, S-33). This is in spite of the fact that simply splitting true DE regions into smaller subsets for testing generally leads to lower power (Figure S-34). The clusters of higher resolution show higher overlap with DE regions (Figure 7c), suggesting that their smaller clusters are more likely to overlap the right set of cells, regardless of the size of the true *N_g_* regions (Figure S-35); the ability to find clusters with high levels of purity of DE cells outweighs the reduction in power due to smaller number of cells in settings where the DE signal diverges from the underlying celltype. Finally, we note that higher resolutions have significantly higher rates of filtering, with many more hypotheses untested (Figure S-36). Yet, in spite of the improved ability of extremely high resolution to detect such DE clusters, their corresponding high levels of false discovery suggest that more moderate choices in resolution provide a more reliable approach.

**Figure 7:**
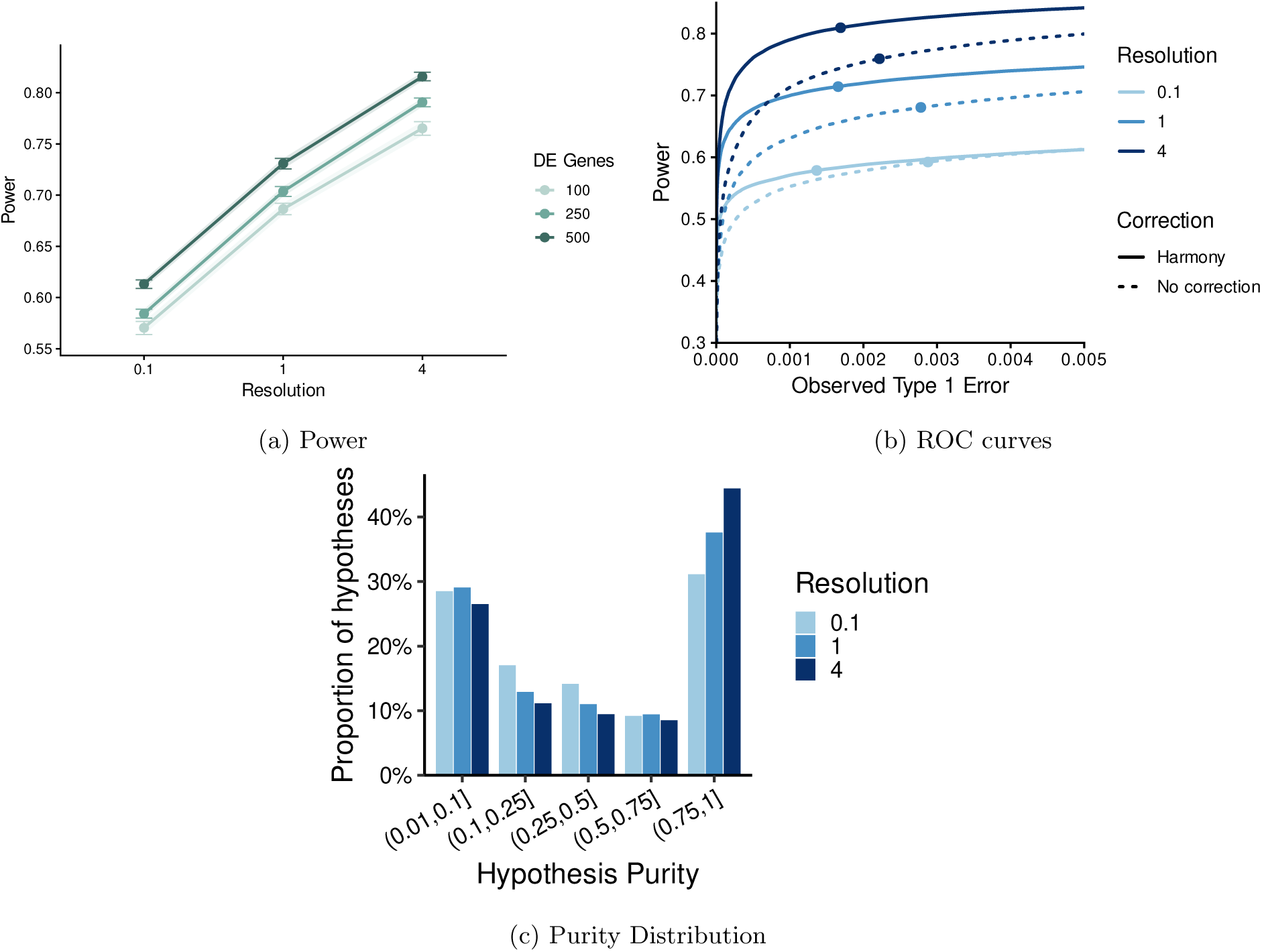
Power, overlap distribution, and ROC curves for varying resolutions. (a) shows power (x-axis) with increasingly higher resolutions (y-axis). Different number of DE genes are shown as differently colored lines. The result is averaged across all simulations regardless of phenotype removal method (no correction, Harmony and fastMNN). We observe similar trends when stratifying separately per correction methods (See Figure S-33). (b) shows the power (y-axis) vs observed type 1 error (x-axis) for varying raw p-value cutoffs. Different resolutions are shown with different colored lines; all have 250 DE genes. The plot displays results with and without Harmony phenotype correction (solid vs dotted respectively). Solid points on each curve are located by determining the raw p-value that would match a BH threshold of 0.1, averaging this across simulations, and then plotting the value of Type 1 error and power that corresponds to this raw p-value cutoff.(c) shows the distribution of cluster overlaps for increasing resolution. In this plot, we are showing results only for without phenotype correction. For all plots, simulations use DE neighborhoods that are not aligned with celltype and Harmony sample-corrected null gene embeddings.

## 3 Methods

### 3.1 Setup: Per cluster differential expression testing

In a single cell experiment, we collect cells *C* = *{c_i,j_}*, where *c_i,j_*is the *i*-th cell from the *j*-th sample. We can consider that each cell *i* from a sample *j* is sampled from the full set of cells of that sample; the number of cells per sample is often in the neighborhood of 2,000-5,000 cells. Cellular features are measured for each cell; for scRNA-Seq, which is our focus here, the measured features correspond to the mRNA levels of genes. The measured quantity per gene *g*, *x_i,j,g_* is a count of the number of times the mRNA of gene *g* was observed in the cell. We will indicate all data across all cells, samples, and features as *X* = *{x_i,j,g_}* with *E*(*x_i,j,g_*) = *ε_i,j,g_*. The number of possible genes is often around 20,000, but frequently only a few thousand of those genes are detected in any particular cell.

Typically, cells are thought to come from distinct cell-types, and for many genes, cells from different cell-types will have *ε_i,j,g_* that differs between cells of different cell-types. However, the cell-type of an observed cell is unknown (i.e. a latent variable) and must be estimated from the observed data *X*. The cell-type is often the largest source of variation between the cells, and the overwhelming purpose of single-cell experiments is to account for this large source of heteroskedasticity. Furthermore, though there may be variations between the cell-types in different sample phenotypes, it is frequently the case that the set of possible cell-types are assumed to be shared across samples and sample phenotypes.

For the purpose of this paper, we will be interested in measuring differential phenotype expression. This corresponds, for a particular gene, to a subset of cells for which their expression level depends on the covariates (or phenotype) of the *sample* from which the cell comes. For simplicity, we will assume the samples are from only two phenotypic groups. We call the set of cells that demonstrate differential expression between phenotypes for a gene a *DE neighborhood*. We assume that a DE neighborhood consists of cells of both phenotypes, so it includes the “right” set of cells to be compared to measure the phenotype effect. In practice, a gene could have multiple non-overlapping DE neighborhoods, meaning that they demonstrate different phenotypic-expression levels in the different DE neighborhoods, but in our simulations we will assume only a single DE neighborhood, *N_g_*, per gene. Genes that do not exhibit any differential expression between phenotypes have *N_g_*= *→*. DE neighborhoods could potentially coincide with cell-types, but they could also span multiple cell-types, be a subset of a cell-type, or other complicated hybrids. Both DE neighborhoods and cell-types are based on unknown, latent variables defined per cell, but it should be emphasized that there is a latent variable per gene for DE neighborhoods, while cell-types are defined based on a single latent variable and not dependent on gene.

#### 3.1.1 Cluster-based DE Analysis

To be able to estimate the differential expression of genes between phenotype, we are missing the latent variable *N_g_* which defines the true set of cells per gene that are differentially expressed. A common strategy is to not estimate a latent variable *N_g_* per gene, but rather to assume that *N_g_* will (roughly) correspond to a cell-type, and instead estimate the unknown cell-type. We will call this strategy a cluster-based DE pipeline.

Unlike estimating per-gene latent DE neighborhoods, estimating the cell-type of a cell is a ubiquitous problem in single-cell data with many well-studied strategies [27]. The most common manner for cell-type estimation is to estimate a set of lower-dimensional latent variables or embeddings across all the cells, for example using PCA or scVI [16], and then cluster the cells in the estimated latent variable space [28] (Figure 2a). The estimated clusters are then associated with the unknown cell-type of the cells.

Once clusters of cells are estimated, the cluster-based DE strategy uses the estimated clusters as a proxy for *N_g_*. Specifically, the *K* clusters, *C*^^^_1_*,…, Ĉ_K_* are considered candidates for being DE neighborhoods, and for each gene *g*, each cluster *k* is evaluated for whether the cells show phenotypic differences in the expression of gene *g*.

To evaluate whether the cluster *k* is DE for a particular gene, the most common strategy [26, 2, 6] divides cells *c_i,j_ ɛ r Ĉ_k_* into their separate samples, and the data *x_i,j,g_* that come from sample *j* is combined into a single summary value *X_g,k,j_* – a process called “pseudo-bulking” (Figure 2a and 2c). Usually the summary value is simply the sum of the counts *x_i,j,g_* (see [6] for overview),

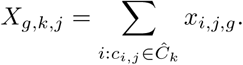

This results in a vector of *n* sample-level values for each gene and cluster combination. The pseudobulk data is then the input for performing for each gene within each cluster a hypothesis test, *H*_0*,g,k*_, as to whether the suitably normalized *X_g,k,j_* has the same expected value in the two phenotypic groups (see Figures 2c, 2d). This is now a standard DE analysis for “bulk” mRNA-Seq, i.e. data not collected on individual cells, and there are many methods that have been optimized for this setting [23, 13, 17] (see [6] for other strategies for sample-level DE other than pseudobulking).

We would note that there is commonly a filtering of both genes and samples in this DE analysis pipeline. In particular, some samples are removed when testing a particular cluster *k* – for example those with low numbers of cells in the cluster – and some *H*_0*,g,k*_ are filtered out and not tested when there are low levels of expression. or low numbers of samples of a particular phenotype (see Supplementary Text, Section S2.2. for more details). Thus while we start with the same number of clusters per gene, we end up testing different numbers of hypotheses per gene and the tests may be based on different numbers of samples; we define the number of hypotheses tested for a gene as *K_g_*, with *K_g_ ≤ K*.

This results, for each gene *g*, in a set of *K_g_* (raw) p-values *p_g,k_*, corresponding to each of the clusters tested for that gene. Those clusters identified as DE will be those with an multiple-testing adjusted p-value *p̃_g,k_ ≤ ϑ* for some value *ϑ*. While adjusting for the FDR over tested genes with the BH procedure is a common method for identifying DE genes in bulk experiments, two different strategies are possible here: either apply the BH method to the entire set of p-values across all *p_g,k_*’s (global control); or apply BH to each cluster independently, aiming at FDR control for all the *p_g,k_*’s within a given cluster *k* (cluster control). Note that having global control cannot guarantee cluster control and *vice versa*. The choice between the two fundamentally depends on the questions the biologist seek to answer. In this work, we choose the global correction as we are not focusing on any particular cell-type or cluster.

We note that other methods have been recently introduced that estimate the DE neighborhoods more adaptively. For example, MiloDE [21] will estimate a large set of small, overlapping clusters *Ĉ_k_*, and then determine which *Ĉ_k_* are DE for a particular gene using similar pseudobulk evaluations. Another approach, LEMUR [1], estimates a large number of gene-specific sets of potential neighborhoods *Ĉ_g,k_*, which are evaluated using pseudobulk methods described above; these methods adaptively define new candidates *Ĉ_g,k_* and test them, and the full set of possible *Ĉ_g,k_* are all possible subsets and thus never explicitly enumerated.

### 3.2 Simulation Study

Here we give a brief overview of the simulation strategy. For more details, see Supplementary Text, Section S1.

#### Creation of Simulated Data

Using data from a study of colorectal tumor [22], we isolate 20 samples that are all from a normal condition. We filter the data to the 5,000 highest variance genes, which constitute the set of null genes for all simulations. For each simulation, we randomly assign samples (and thus their cells) to one of two phenotypes; this means that the expression value of the null genes will remain as a block across simulations for a given sample and their cells, but shuffled between phenotypes. To these null genes, we add specified quantities of DE genes. Constructing the gene expression levels for a DE gene *g* consists of two steps: 1) determining which cells will be part of the DE neighborhood *N_g_* and 2) creating expression values for all cells in all patients for the gene. This strategy modifies that used by [1] for evaluating their method Lemur.

To choose *N_g_* for a given DE gene, we first define the possible set of *N_g_* by performing a clustering of cells based on the fixed set of null genes. This clustering is based on a lower dimensional embedding of the gene expression data of the null genes, which have no phenotypic differences. However, there may be differences or separation of cells due to sample variation present in the data; while biological variability is expected between patients, we want the definition of “true” DE neighborhoods in our simulation to not be impacted by these sample differences. Therefore we use embeddings of the null genes that have removed potential sample differences using Harmony [12]; to avoid potential bias in favor of Harmony, we also run simulations based on embeddings using an alternative batch-correction method, fastMNN [7] to remove sample effects, and also PCA embeddings with no sample correction. These different embeddings lead to noticeable differences in clusters (see Figure S-1), with clusters from Harmony embeddings more similar to clusters found from PCA embeddings with no sample correction, compared to those of fastMNN.

For a particular choice of embeddings we then cluster the cells in this embedding space to form a set of possible *N_g_* regions; two different strategies of clustering are implemented 1) Leiden clustering [28] or 2) *K*-means clustering [19] for three different choices of *k* (*k* = 3, 10, or, 20). The two different choices represent different choices about how closely the DE regions *N_g_* will be expected to follow the data-derived clusters, *Ĉ_k_* (which will be based on Leiden clustering as well). Leiden clustering of the null genes will result in DE neighborhoods *N_g_* that will be quite similar to the clusters we will get when we cluster the simulated data, with any differences in the embeddings primarily introduced by the DE genes. The K-means simulation strategy, however, results in DE neighborhoods that are quite different from the clusters we would get from Leiden clustering of the simulated data even when there is no phenotypic differences – *N_g_* may span multiple clusters, be a subset of some cluster, or combinations in between (see Figure S-2). For this reason, we when the *N_g_* regions are chosen by Leiden clustering of the null genes, we will refer to this as the cell-type *aligned* strategy, and when *N_g_*regions are chosen by the K-means clustering of the null genes as the *non-aligned* strategy.

Once the possible set of DE regions is determined, then for each DE gene, we randomly select one of these regions to be the set of DE cells *N_g_*. Once *N_g_* is chosen for a gene, we create expression levels for this gene for *all* cells by simulating from a negative binomial (Gamma–Poisson) model, with a log-linear mean that includes a baseline expression, a treatment effect which is restricted to DE cells, and a sample-level random effect (see Supplementary Text S1.4). Notably for a given gene, the baseline expression is shared across cells outside of the DE neighborhood and cells inside the DE neighborhood assigned to control samples. Any natural clustering of cells present in null genes (potentially suggesting biological cell-type structure) is not preserved in simulated genes, where the simulated distribution of the gene is differentiated only due to phenotypes.

#### Clustering and DE analysis of the Simulated Data

For each simulated dataset, we create low-dimensional embeddings of the data and cluster the cells in this embedding space using Leiden clustering. The baseline choice of embedding is the top 50 Principal Components based on performing PCA on the 2,000 most highly variable genes. We also consider an “oracle” embedding based on the PCA embedding of only the 2,000 most highly variable *null* genes, which have no phenotype difference, similar to the step performed above in choosing the set of possible DE regions. Note that because our null genes stay the same for all simulations and only the phenotypic labels of samples are permuted, this means that for the simulation setting where the set of choices of *N_g_* is based on the Leiden method, the set of possible true *N_g_* regions and the clusters identified based on our oracle clustering of the simulated data are practically equivalent, except for differences due to different initializations of the clustering algorithm. We also consider embeddings based on phenotype-removal methods intended to remove batch effects or other signal (see Supplemental Text, Section 3.3 for details).

After the clustering is run for a particular embedding, we then pseudo-bulk the cells per cluster for each sample and run edgeR [23] to test for differences between the two phenotypes per gene on the pseudo-bulk data for each cluster. This results in DE results (i.e. p-values) for each gene in each cluster, except for gene-cluster pairs which have been filtered out (see Supplementary Text, Section S2.3). We then adjust the raw p-values for multiple testing by performing a Benjamini-Hochberg correction across the p-values from all non-filtered genes and clusters.

#### Evaluation of Results

Because the clusters are determined based on the data, we have to determine whether a null hypothesis *H*_0*,g,k*_ corresponding to a cluster *Ĉ_k_* should be considered as truly null for the purposes of determining which discoveries are true or false. To do this, we consider purity of a hypothesis, defined to be the percentage overlap of the cluster *Ĉ_k_* with the true DE neighborhood *N_g_* (see Supplementary Text Section S2 for details). Often, we will consider that *any* overlap of *Ĉ_k_* with *N_g_* means that *H*_0*,g,k*_ is non-null, but for power calculations, we will require a higher level of minimum purity.

### 3.3 Correction Methods Overview

This section provides a description of each of the methods that were used to correct sample effects both in the embeddings used for the generation of *N_g_* (Section S1) and for clustering of the simulated data (See section 3.4). All of these corrections methods have as initial input embedding the PCA of all *G_null_* + *G_DE_* genes.

#### Harmon

Harmony [12] employs an iterative soft-clustering framework to estimate and subtract linear batch effects while preserving biological variation. In each iteration, cells are soft-assigned to clusters based on their current embedding coordinates (input PCA in the first iteration), and cluster-specific correction factors for the batch covariate are estimated via ridge-regularized regression for each PC coordinate. A correction factor is then computed for each cell as a weighted sum of the above cluster-specific factors based on the cells soft membership to each cluster. Once these factors are subtracted from all cells, the resulting embedding, of same dimensionality as the input PCA embeddings serves as the initial embedding for a new iteration of correction.

#### fastMNN

fastMNN (`batchelor::fastMNN`, [7]) constructs a batch-corrected embedding by sequential pairwise alignment of batches in a low-rank representation of the expression matrix. Within each pair of batches it identifies mutual nearest neighbour (MNN) cells—pairs that are each other’s closest match in the opposite batch in PCA space—treats their average expression displacement as an estimate of the batch effect, and corrects that effect. Corrected coordinates from successive batches are merged into a single reference embedding.

#### Type versus state (TvsS) feature selection

Following the “type-not-state” motivation of Wang, Crowell, and Robinson (2025) [31], we distinguished transcriptional programs associated with discrete substructure (cell type) from those driven by the treatment contrast (cell state or condition). Using `muscat::aggregateData` [6], Pseudobulk profiles are extracted from an initial clustering of the full gene matrix and are fed to a linear mixed model with random intercepts for cluster, sample, and condition. Denote by *t*PVE*_g_* and *s*PVE*_g_*the estimated variance fractions for cluster and condition respectively. Genes were ranked by rank(*t*PVE*_g_*)*—*rank(*s*PVE*_g_*), and the chosen number of top genes (default 5000) are identified as most likely reflective of cell-type differences (rather than condition differences).The count data restricted to these 5000 genes is used as the starting cell by gene count data for the downstream differential analysis pipeline.

### 3.4 Downstream Differential Analysis Pipeline

Each pipeline begins from a cell by gene count matrix (for simulated counts, genes would be *G*_DE_ *↗ G*_null_). Using Seurat, gene counts are normalized per cell and log transformed to a common sequence (using `Seurat::NormalizeData`, [8]). The highest variable genes (HVGs) were then found and standardized (using `Seurat::FindVariableFeatures` and `Seurat::ScaleData`, [8] with default 2000 top variance genes) as inputs to a Principle Component Analysis (PCA). 50 Principal Components were then retained (using

Seurat::RunPCA, [8]) to serve as the input embeddings (where appropriate) into one the following strategies to get corrected embeddings where the clustering will be performed (see Section for more detail):

- **No correction** - input PCA Embeddings are used directly, reflecting no phenotype correction.
- **Harmony correction.** Harmony corrected embeddings are used after setting the nuisance/batch parameter to the sample identifier.
- **fastMNN.** fastMNN corrected embeddings are used after setting the nuisance/batch parameter to the sample identifier.
- **Type versus state (TvsS) feature selection.** After finding the genes with least condition specific variance, ranked by rank(*t*PVE*_g_*) *↔* rank(*s*PVE*_g_*) (see 3.3), PCA is rerun on this gene subset only.

#### Null-gene versions of embeddings

To build a representation deliberately blind to the DE signal, we subset the object to genes that are in *G*_null_, calculate their PCA embeddings, and recreate “null gene” versions of each of the above (No correction, Harmony and fastMNN phenotype removal methods). Pseudobulk DE was still performed on the *full* gene set (*G*_null_ *↗ G*_DFE_), but with cluster assignments based on these null gene versions of the embeddings.

#### Clustering, Pseudobulk and Raw P-values

After obtaining the desired embedding depending on the phenotype removal correction, Leiden clustering was performed on those embeddings with resolution of 0.1 to obtain found or candidate clusters *Ĉ_k_*. For every resulting cluster assignment, counts were pseudo-bulked (summed within each cluster–sample combination). Treatment versus control was tested with using muscat package (`muscat::pbDS`) with edgeR (quasi-likelihood) framework. Raw *p*-values from all cluster–gene tests were pooled and adjusted jointly using the Benjamini–Hochberg (BH) procedure, producing the per-gene, per-cluster adjusted *p*-values used in the evaluation which we denote by *p̃_g,k_*. Unless otherwise noted, for all false discovery rate or proportions, we use a cutoff value or FDR threshold, of 0.1.

### 3.5 Adjusted Mutual Information (AMI) between cluster and phenotype

Let *U* denote the random condition label and *V* the random cluster label for a cell. From the contingency table of counts *n_ck_* condition *c*, cluster *k*) with 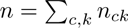, we compute Shannon entropy using the natural logarithm, e.g.

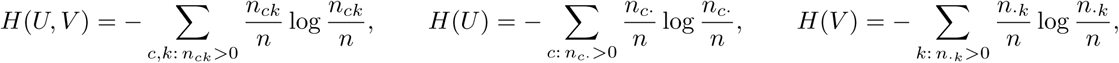

with row sums *n_c·_* and column sums *n_·k_*. The (empirical) mutual information is

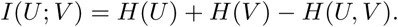

To account for random chance for fixed marginals, we use the definition of Adjusted Mutual Information defined in [29]. Letting EMI denote E[*I*(*U*; *V*)]:

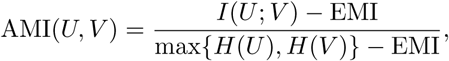

This defines the default adjusted mutual information returned by `aricode::AMI` in R. Values near 1 indicate agreement between cluster and condition beyond what is expected from the margins alone; values near 0 indicate association comparable to chance under the fixed-margins null.

Since we are interested in the effect of this imbalance on DE error rates, we limit the starting contingency table to clusters that were not fully filtered out at the DE stage.

#### 3.5.1 Size-weighted sample Pearson Residual

To score imbalance at the cluster level rather than the entire experiment level, we use a Pearson residual on sample counts, which fundamentally consider the sample the experimental unit. Under a naive null where all control samples are of same size and are assigned independently with probability equal to the global prevalence of control samples 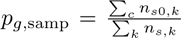 is the number of control samples in cluster *k* and*n_s,k_* is the total number of non-empty samples in cluster *k*). The expected number of control samples in cluster *c* is then *E*_0,samp*,c*_ = *n_s,c_ p_g,_*_samp_. The absolute value of the sample Pearson residual is then:

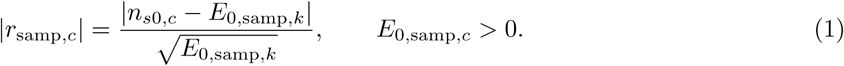

Each sample present in the cluster contributes equally, regardless of how many cells it contributes.

To capture the fact that certain samples have larger number of cells than others, we use a size-weighted sample Pearson residual. The null still assumes that samples are chosen independently but weigh each sample by its cell count in the cluster *w*_samp*,k*_. Writing as before *n*_0*,k*_ = Σ*_s_ w*_samp*,k*_ *I_s_* with *I_s_* ∼ Bernoulli(*p_g,_*_samp_) independent under the null,

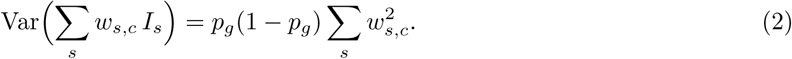

Matching the Pearson denominator convention used for the unweighted residual, we define:

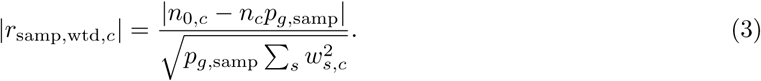

Unequal sizes inflate 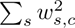 relative to 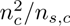, so the size-weighted residual is typically smaller than a naive cell Pearson residual for the same absolute difference in condition prevalence.

## 4 Discussion

This work investigates post-selection inference issues in the standard cluster-based, patient-level differential expression (DE) pipelines for scRNA-Seq data. Using a simulation framework, we investigate how the reuse of the data in both clustering the cells as well as testing their differences impacts both FDR control and statistical power.

Our results demonstrates that a naive clustering pipeline which does not remove phenotype effects can lead to a severe loss of FDR control in the presence of differentially expressed genes, but only with relatively strong DE signal. Encouragingly, we also show that for standard levels of cluster resolution, routine batch and sample correction methods meant to correct for unwanted sample variation also effectively remove phenotypic effects and mitigate both the loss of FDR control and the degradation of power as phenotypic differences increase. At the same time, we do not see a drop in power due to use of batch corrections. Consequently, our work provides inference-related support for employing these sample correction tools, beyond their standard benefits with regards to biological and technical integration.

We also find that FDR deterioration is directly tied to the dependency between the phenotype effect and the found cluster, an association that can be measured directly on data. We suggest the Adjusted Mutual Information (AMI) to measure the association between clusters and phenotypes, and demonstrate that it can serve as a direct measure of phenotype leakage and an indicator of post-selection inference risk. Researchers can leverage this metric as a diagnostic tool when evaluating integration techniques and assessing real-world datasets.

Finally, our power results demonstrate problems underlying this pipeline if the basic assumption is violated, namely that the found clusters are good proxies for the true DE neighborhoods. We show that poor overlap of the found clusters with the true DE neighborhoods has negative effects on the power. This indicates that even if clustering accurately finds true cell types, genes that are DE in a small proportion of a cell-type or in sets of cells that do not align well with a cell-type will be difficult to identify with such strategies. Choosing lower resolution clusters can improve the power to detect such DE behavior, but at the significant risk of higher false discovery rates. This is true even in simulations with no DE genes, indicating that just sample variation may lead to false discoveries in high resolution clusters. Our evaluation of different cluster resolutions also provide context for other DE frameworks—such as MiloDE [21] or treeclimbR [9] that both build upon small cell clusters.

## Acknowledgments

We thank Pierre Neuvial for useful feedback on earlier drafts of this work. This work was supported by NIH grant R01GM144493. Elizabeth Purdom is a Biohub San Francisco, Investigator.

*AI Acknowledgment* AI was used in this work for debugging, documenting and generating visualization scripts - including Cursor and Google’s Gemini for code development - as well as to provide feedback on grammar and phrasing of authors’ manuscript drafts. Any AI-assisted output that was used was at all times reviewed and tested by the authors.

## Author contributions

All authors contributed to the design and conception of the paper. N.S. performed the simulations and data analysis. N.S. and EP wrote the initial manuscript, and all authors contributed to editing the manuscript. All authors reviewed and approved the final version.

## Data Availability

The four scRNA-seq datasets used for simulations and analysis are publicly available. Three datasets are available under the Gene Expression Omnibus. The Pelka [22], Melms [20], and Kang [10] datasets have accession codes GSE178341, GSE171524 and GSE96583 respectively. The Liu dataset [15] is available under the European Genome-Phenome Archive with dataset ID EGAD00001008624.

## Code Availability

Code and scripts to recreate the results presented in this study are available at https://github.com/epurdom/scSampleSimScripts.

## References

[1] Constantin Ahlmann-Eltze and Wolfgang Huber. Analysis of multi-condition single-cell data with latent embedding multivariate regression. Nature Genetics, 57(3):659–667, March 2025.

[2] Robert A. Amezquita, Aaron T. L. Lun, Etienne Becht, Vince J. Carey, Lindsay N. Carpp, Ludwig Geistlinger, Federico Marini, Kevin Rue-Albrecht, Davide Risso, Charlotte Soneson, Levi Waldron, Hervé Pagès, Mike L. Smith, Wolfgang Huber, Martin Morgan, Raphael Gottardo, and Stephanie C. Hicks. Orchestrating single-cell analysis with Bioconductor. Nature Methods, 17(2):137–145, February 2020.

[3] Richard Bourgon, Robert Gentleman, and Wolfgang Huber. Independent filtering increases detection power for high-throughput experiments. Proceedings of the national academy of sciences, 107(21):9546– 9551, 2010.

[4] Andrew Butler, Paul Hoffman, Peter Smibert, Efthymia Papalexi, and Rahul Satija. Integrating single-cell transcriptomic data across different conditions, technologies, and species. Nature biotechnology, 36(5):411–420, 2018.

[5] Emmanuel J Candès, Xiaodong Li, Yi Ma, and John Wright. Robust principal component analysis? Journal of the ACM (JACM), 58(3):1–37, 2011.

[6] Helena L Crowell, Charlotte Soneson, Pierre-Luc Germain, Daniela Calini, Ludovic Collin, Catarina Raposo, Dheeraj Malhotra, and Mark D Robinson. Muscat detects subpopulation-specific state transitions from multi-sample multi-condition single-cell transcriptomics data. Nature communications, 11(1):6077, 2020.

[7] Laleh Haghverdi, Aaron TL Lun, Michael D Morgan, and John C Marioni. Batch effects in single-cell rna-sequencing data are corrected by matching mutual nearest neighbors. Nature biotechnology, 36(5):421–427, 2018.

[8] Yuhan Hao, Stephanie Hao, Erica Andersen-Nissen, William M Mauck, Shiwei Zheng, Andrew Butler, Maddie J Lee, Aaron J Wilk, Charlotte Darby, Michael Zager, et al. Integrated analysis of multimodal single-cell data. Cell, 184(13):3573–3587, 2021.

[9] Ruizhu Huang, Charlotte Soneson, Pierre-Luc Germain, Thomas S.B. Schmidt, Christian Von Mering, and Mark D. Robinson. treeclimbR pinpoints the data-dependent resolution of hierarchical hypotheses. Genome Biology, 22(1):157, May 2021.

[10] Hyun Min Kang, Meena Subramaniam, Sasha Targ, Michelle Nguyen, Lenka Maliskova, Elizabeth McCarthy, Eunice Wan, Simon Wong, Lauren Byrnes, Cristina M Lanata, et al. Multiplexed droplet single-cell rna-sequencing using natural genetic variation. Nature biotechnology, 36(1):89–94, 2018.

[11] Günter Klambauer, Karin Schwarzbauer, Andreas Mayr, Djork-Arne Clevert, Andreas Mitterecker, Ulrich Bodenhofer, and Sepp Hochreiter. cn. mops: mixture of poissons for discovering copy number variations in next-generation sequencing data with a low false discovery rate. Nucleic acids research, 40(9):e69–e69, 2012.

[12] Ilya Korsunsky, Nghia Millard, Jean Fan, Kamil Slowikowski, Fan Zhang, Kevin Wei, Yuriy Baglaenko, Michael Brenner, Po-ru Loh, and Soumya Raychaudhuri. Fast, sensitive and accurate integration of single-cell data with harmony. Nature methods, 16(12):1289–1296, 2019.

[13] Charity W Law, Yunshun Chen, Wei Shi, and Gordon K Smyth. voom: Precision weights unlock linear model analysis tools for RNA-seq read counts. Genome Biology, 15(2):1–17, 2014.

[14] Yumei Li, Xinzhou Ge, Fanglue Peng, Wei Li, and Jingyi Jessica Li. Exaggerated false positives by popular differential expression methods when analyzing human population samples. Genome biology, 23(1):79, 2022.

[15] Yale Liu, Hao Wang, Mark Taylor, Christopher Cook, Alejandra Martínez-Berdeja, Jeffrey P North, Paymann Harirchian, Ashley A Hailer, Zijun Zhao, Ruby Ghadially, et al. Classification of human chronic inflammatory skin disease based on single-cell immune profiling. Science immunology, 7(70):eabl9165, 2022.

[16] Romain Lopez, Jeffrey Regier, Michael B Cole, Michael I Jordan, and Nir Yosef. Deep generative modeling for single-cell transcriptomics. Nature methods, 15(12):1053–1058, 2018.

[17] Michael I Love, Wolfgang Huber, and Simon Anders. Moderated estimation of fold change and dispersion for RNA-seq data with DESeq2. Genome Biology, 15(12):550, 2014.

[18] Leland McInnes, John Healy, and James Melville. Umap: Uniform manifold approximation and projection for dimension reduction. *arXiv preprint arXiv:1802.03426*, 2018.

[19] James B McQueen. Some methods of classification and analysis of multivariate observations. In Proc. of 5th Berkeley Symposium on Math. Stat. and Prob., pages 281–297, 1967.

[20] Johannes C Melms, Jana Biermann, Huachao Huang, Yiping Wang, Ajay Nair, Somnath Tagore, Igor Katsyv, André F Rendeiro, Amit Dipak Amin, Denis Schapiro, et al. A molecular single-cell lung atlas of lethal covid-19. Nature, 595(7865):114–119, 2021.

[21] Alsu Missarova, Emma Dann, Leah Rosen, Rahul Satija, and John Marioni. Leveraging neighborhood representations of single-cell data to achieve sensitive DE testing with miloDE. Genome Biology, 25(1):189, July 2024.

[22] Karin Pelka, Matan Hofree, Jonathan H Chen, Siranush Sarkizova, Joshua D Pirl, Vjola Jorgji, Alborz Bejnood, Danielle Dionne, William H Ge, Katherine H Xu, et al. Spatially organized multicellular immune hubs in human colorectal cancer. Cell, 184(18):4734–4752, 2021.

[23] Mark D Robinson, Davis J McCarthy, and Gordon K Smyth. edger: a bioconductor package for differential expression analysis of digital gene expression data. bioinformatics, 26(1):139–140, 2010.

[24] Dongyuan Song, Siqi Chen, Christy Lee, Kexin Li, Xinzhou Ge, and Jingyi Jessica Li. Synthetic control removes spurious discoveries from double dipping in single-cell and spatial transcriptomics data analyses. In International Conference on Research in Computational Molecular Biology, pages 400–404. Springer, 2025.

[25] Dongyuan Song and Jingyi Jessica Li. Pseudotimede: inference of differential gene expression along cell pseudotime with well-calibrated p-values from single-cell rna sequencing data. Genome biology, 22(1):124, 2021.

[26] Jordan W. Squair, Matthieu Gautier, Claudia Kathe, Mark A. Anderson, Nicholas D. James, Thomas H. Hutson, Rémi Hudelle, Taha Qaiser, Kaya J. E. Matson, Quentin Barraud, Ariel J. Levine, Gioele La Manno, Michael A. Skinnider, and Grégoire Courtine. Confronting false discoveries in single-cell differential expression. Nature Communications, 12(1):5692, September 2021.

[27] Tim Stuart, Andrew Butler, Paul Hoffman, Christoph Hafemeister, Efthymia Papalexi, William M Mauck, Yuhan Hao, Marlon Stoeckius, Peter Smibert, and Rahul Satija. Comprehensive integration of single-cell data. cell, 177(7):1888–1902, 2019.

[28] Vincent A Traag, Ludo Waltman, and Nees Jan Van Eck. From louvain to leiden: guaranteeing well-connected communities. Scientific reports, 9(1):5233, 2019.

[29] Nguyen Xuan Vinh, Julien Epps, and James Bailey. Information theoretic measures for clusterings comparison: is a correction for chance necessary? In Proceedings of the 26th annual international conference on machine learning, pages 1073–1080, 2009.

[30] Hao Wang, William Torous, Boying Gong, and Elizabeth Purdom. Visualizing scrna-seq data at population scale with gloscope. Genome biology, 25(1):259, 2024.

[31] Jiayi Wang, Helena L Crowell, and Mark D Robinson. On feature selection to disentangle cell type and state transcriptional programs. BMC genomics, 26(1):1006, 2025.

[32] Jesse M Zhang, Govinda M Kamath, and N Tse David. Valid post-clustering differential analysis for single-cell rna-seq. Cell systems, 9(4):383–392, 2019.

