## Supplemental Figures for "“Double-dipping” in testing for phenotypic differences with scRNA-Seq"

### Supplementary Figures

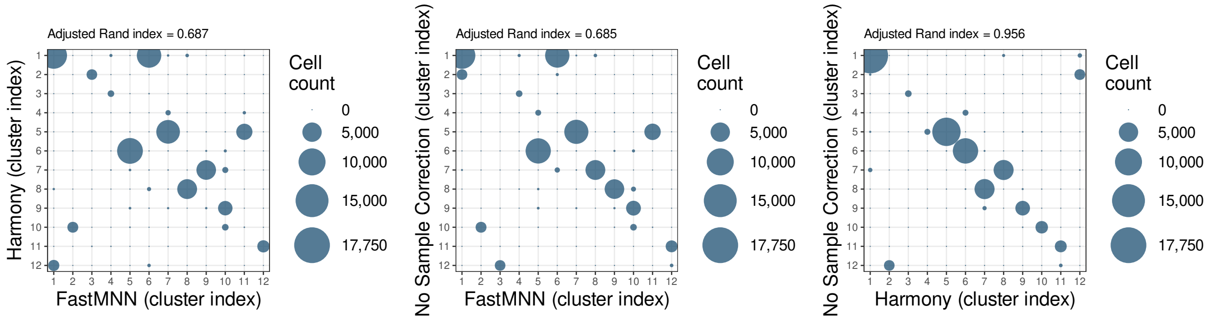


(a) Harmony vs fastMNN (b) No Correction vs fastMNN (c) No Correction vs Harmony

Figure S-1: **Pairwise comparisons of Leiden cluster assignments using different corrected cell embeddings applied only to null genes.** X and Y axes represent cluster indices and bubble sizes represent raw cell-count overlap between the two clusters found on null genes. Adjusted Rand Index is also shown in the figure subtitle.

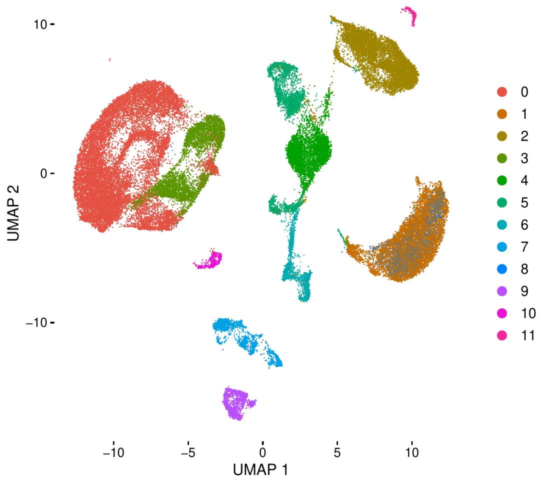


(a) Leiden Clusters with resolution 0.1

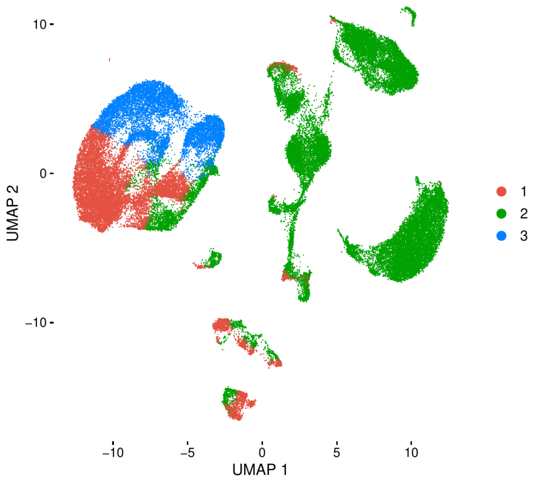


(b) K-Means Clusters with $K = 3$

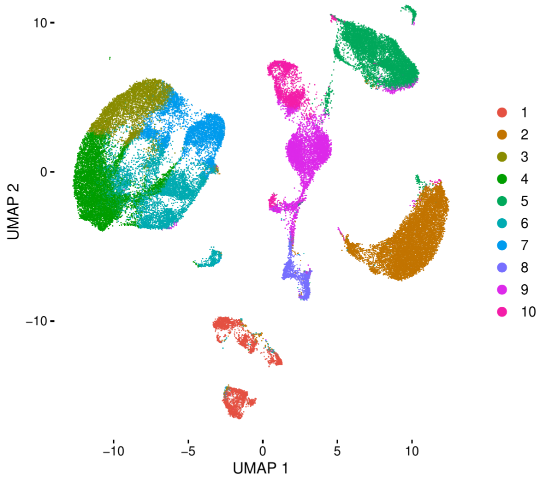


(c) K-Means Clusters with $K = 10$

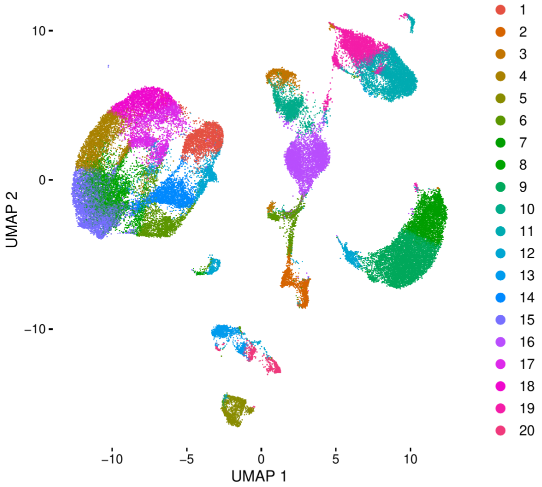


(d) K-Means Clusters with $K = 20$

Figure S-2: **Uniform manifold approximation and projection (UMAP) of the different clusterings of the null genes** All plots show the UMAP [18] visualizations of PCA embeddings found when run on null genes (no simulated DE genes) according to the regular pipeline (see 3.4). (a) shows the clusterings for the celltype aligned setting where the estimated cell-types will be more likely to closely match underlying DE neighborhoods (and in the case where only null genes are clustered will be the same up to randomness in the clustering algorithm). (b)-(d) show the celltype non-aligned setting where the DE neighborhoods will differ significantly from the estimated cell-types, with different choices of $K$ in the $K$-means clustering determining whether the DE neighborhoods are likely to span many estimated cell-types (e.g. K=3) or to be subsets of the estimated cell-types (e.g. K=20).

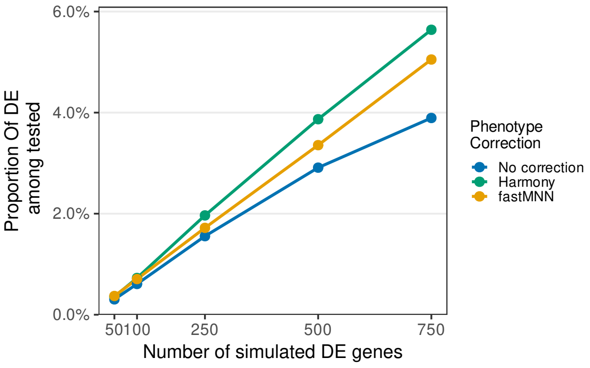


(a) Celltype Aligned DE Neighborhoods

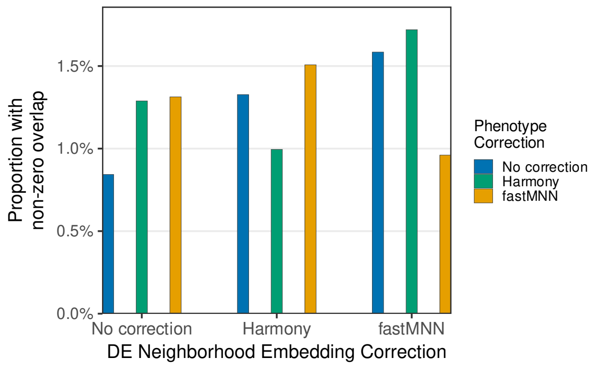


(b) Celltype Aligned DE Neighborhoods - 100 DE Genes

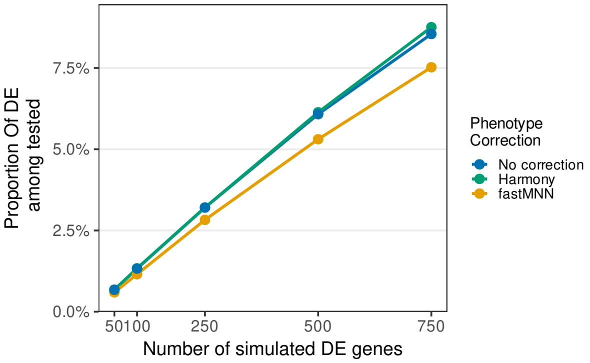


(c) Celltype Non-Aligned DE Neighborhoods

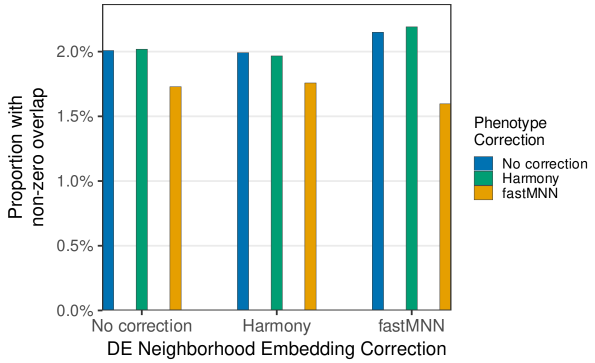


(d) Celltype Non-Aligned DE Neighborhoods - 100 DE Genes

Figure S-3: **Proportion of Hypotheses with non-zero overlap with DE Neighborhoods** Values show the proportion of cluster-gene hypotheses with any overlap averaged across simulations. (a) and (c) show the proportion with increasing number of DE genes. Here the set of averaged simulations are all simulations where the correction method used on the null genes for DE neighborhood creation matches the phenotype correction used on the downstream analysis. (d) and (b) instead stratify by each combination of DE neighborhood embedding correction and phenotype removal method combination, holding the number of DE genes fixed at 100.

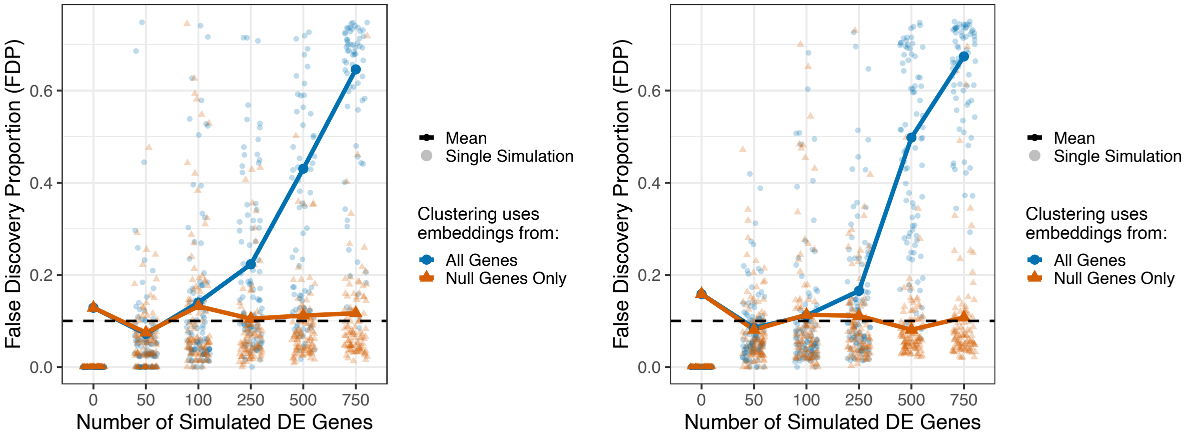


(a) Celltype Aligned DE, lfc = 3 (b) Celltype Non-Aligned DE, lfc = 3

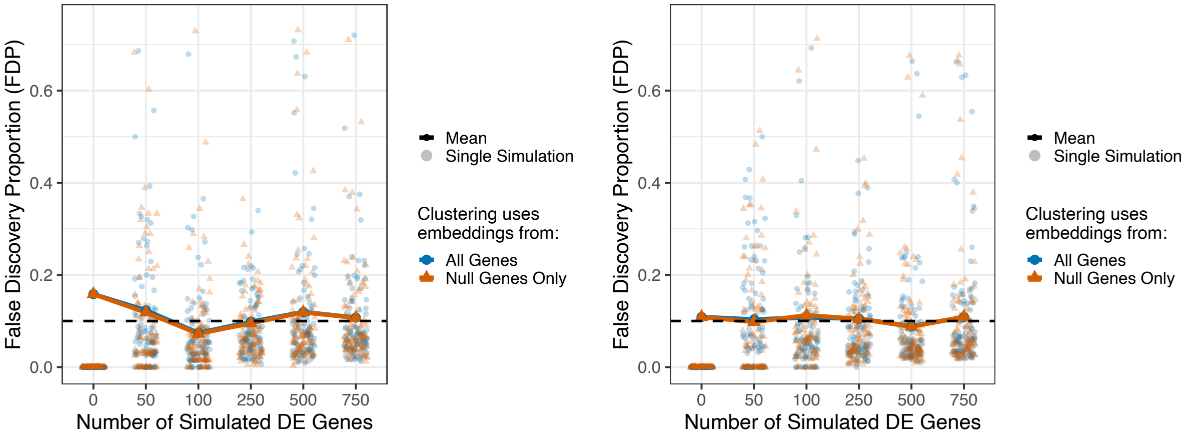


(c) Celltype Aligned DE, lfc = 1.5 (d) Celltype Non-Aligned DE, lfc = 1.5

Figure S-4: **Loss of FDR control using clustering on embeddings without phenotype removal** Each transparent point represents the false discovery proportion for a simulated experiment with 5000 null genes and the number of differentially expressed genes indicated by the x-axis and the average (FDR) across simulations shown as solid line. In panels on the left (a and c) $N_g$ are chosen to be celltype aligned, namely Leiden clusters on harmony corrected embeddings of the null genes. In panels on the right (b and d) the DE cells chosen using the second, celltype non-aligned strategy, namely $K$-means clustering for $k = 3, 10, 20$ on the harmony corrected embedings on the null genes. The top row (a and b) shows simulations with high levels of log fold-change differences in the DE genes (average of 3), while the bottom row (c and d) shows low levels of log fold-change (average of 1.5).

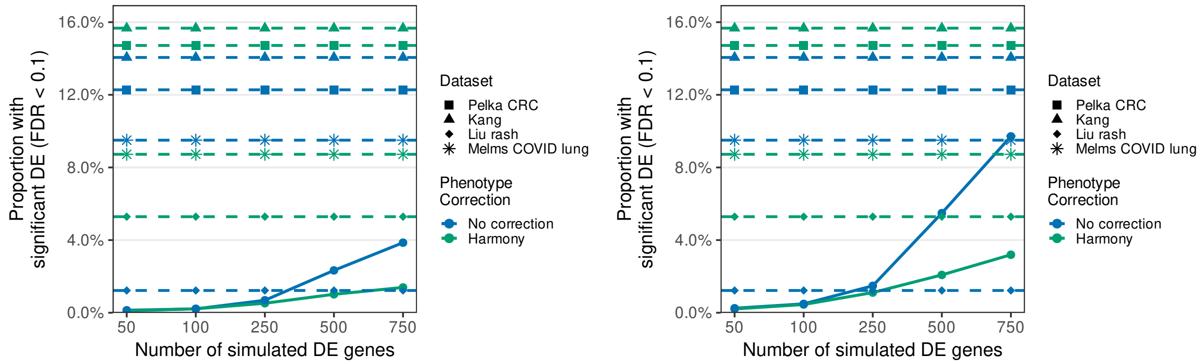


(a) Leiden DE Neighborhoods

(b) K-Means DE Neighborhoods

Figure S-5: **Proportion of gene/cluster pairs found to be significant among all hypotheses** Values show the proportion of cluster-gene hypotheses found to be significant after the DE analysis pipeline. Horizontal lines of different shapes show the result from doing the analysis on real datasets as described in S4 with different phenotype removals. Solid lines represent the proportions for simulations with different phenotype removal methods and number of DE genes. (S-5a) captures the approach where $N_g$ is created with Leiden clusters vs the K-means approach captured by (S-5b) as described in S1. Both plots are for simulations for mean lfc of 3.

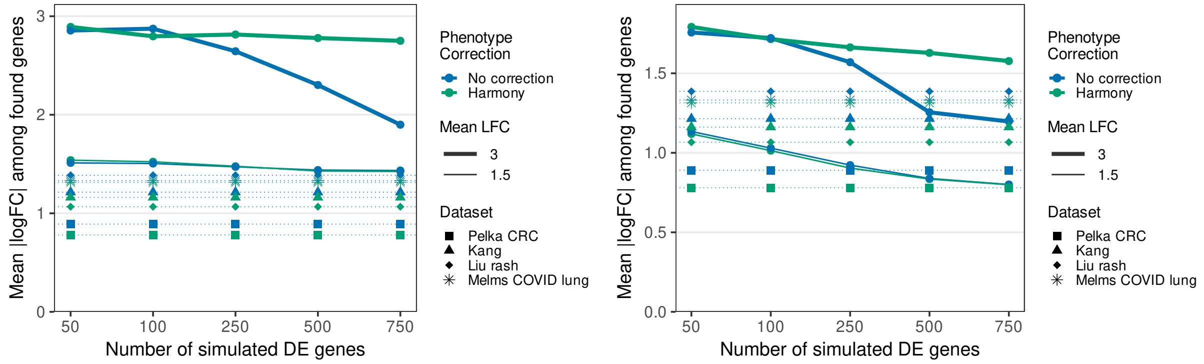


(a) Celltype aligned DE Neighborhoods

(b) Celltype non-aligned DE Neighborhoods

Figure S-6: **Comparison of the mean absolute log-fold change in simulations vs in real data** Y-axis represents the the mean absolute log fold change of cluster-gene hypotheses found to be significant after the DE analysis pipeline. Dotted horizontal lines of different shapes show the results from doing the analysis on real datasets as described in S4 with different phenotype removals. Solid lines represent the proportions for simulations with different phenotype removal methods and number of DE genes. (S-6a) captures the approach where $N_g$ is created with Leiden clusters vs the K-means approach captured by (S-6b) as described in $S1.3$. Simulations with mean log fold change of 3 and 1.5 are plotted for both solid lines (thick and thin lines).

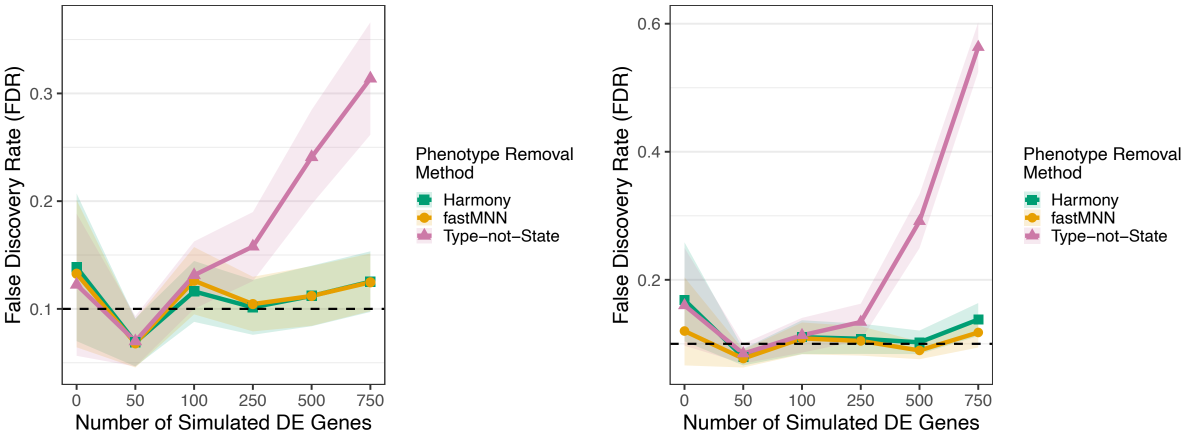


(a) Celltype aligned $N_g$ (b) Celltype non-aligned $N_g$

Figure S-7: **Comparison of FDR for Different Methods for** $N_g$ FDP for simulations with varying number of DE genes and different phenotype removal methods. For all cases the Harmony corrected embeddings of null genes were used for determining $N_g$, with average log fold change of 3.

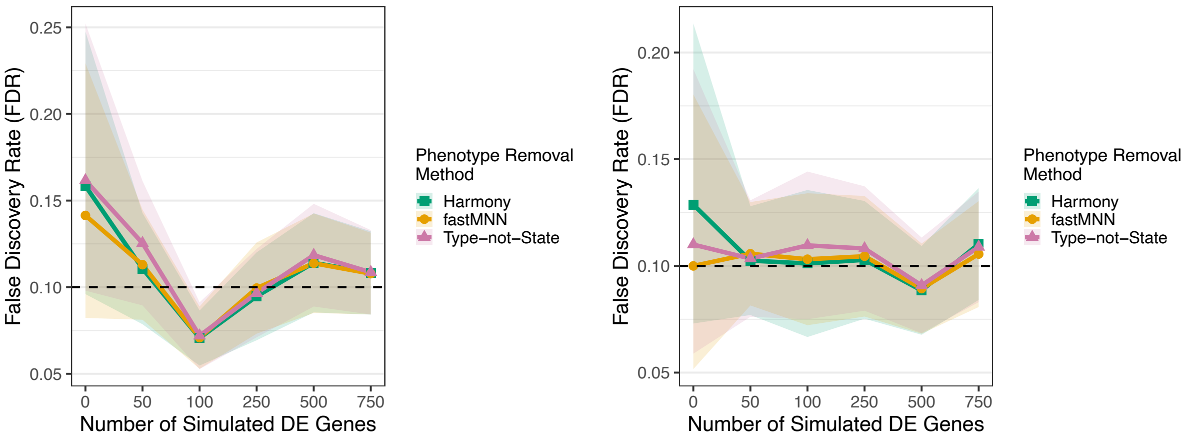


(a) $N_g$ chosen by original Leiden clusters (b) $N_g$ chosen by K-means for $k = 3, 10, 20$.

Figure S-8: **Comparison of FDR for Different Methods for** $N_g$ **with low log fold change** FDP for simulations with varying number of DE genes and different phenotype removal methods. For all cases the Harmony corrected embeddings of null genes were used for determining $N_g$, and average log fold change of 1.5.

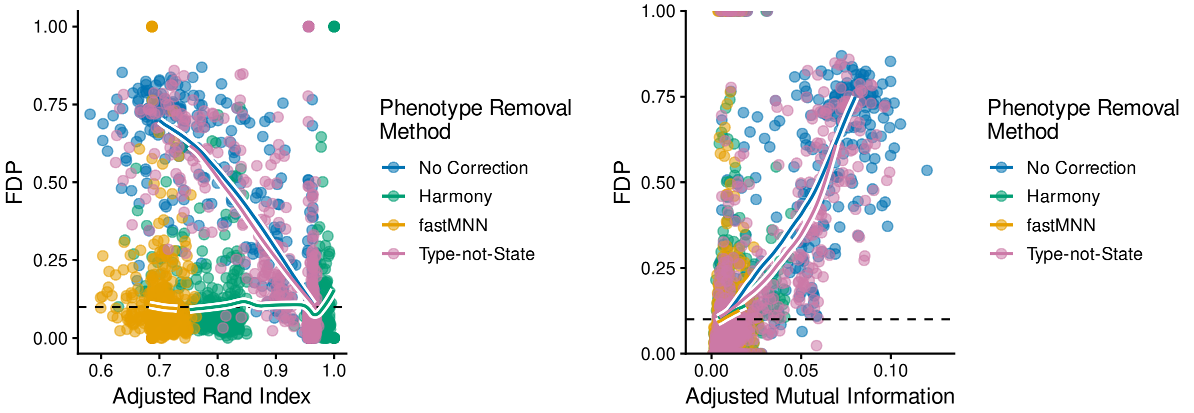


(a) Adjusted Rand Index

(b) Adjusted Mutual Information

Figure S-9: **Metrics for potential phenotype bias detection using celltype non-aligned** $N_g$ (a) Adjusted Rand Index per simulation between the simulation's found clusters and the oracle cell-types used to define $N_g$, a measure of how well the empirical clusters correspond to the ground-truth differential neighborhoods $N_g$. (b) The Adjusted Mutual Information per simulation between cluster and phenotype labels for all cells, measuring how much empirical clusters reflect phenotypic differences. Simulation settings are with $N_g$ chosen from K-means clustering of the Harmony embedding of null genes with an average true log-fold change of 3. Plotted are the results across all simulations, regardless of the number of DE genes. Colored trend lines, intended as visual summaries only, are local linear smooths computed as triangle-weighted averages over a fixed-width, moving window set to half the size of the full $x$-range, excluding the most extreme 10% data points on either tail.

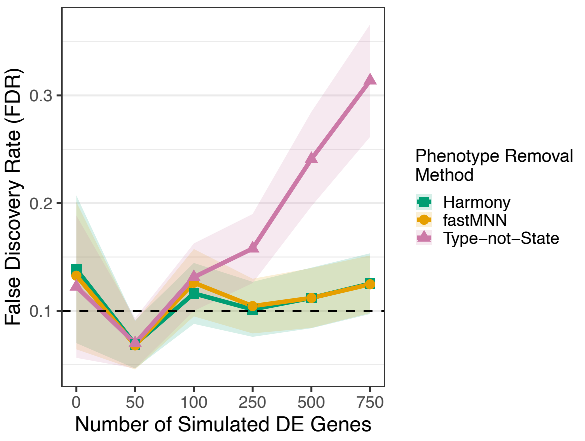


(a) Celltype Aligned DE, lfc = 3.0

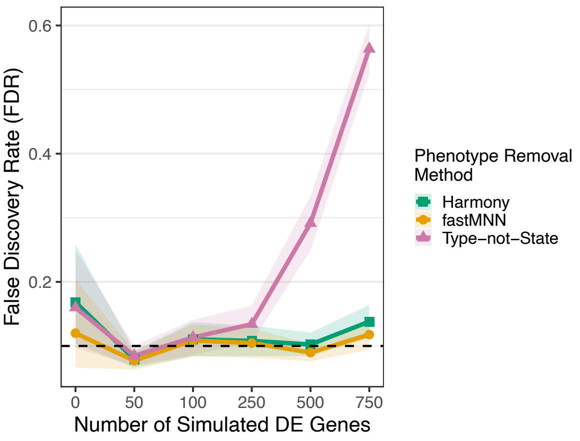


(b) Celltype Non-Aligned DE, lfc = 1.5

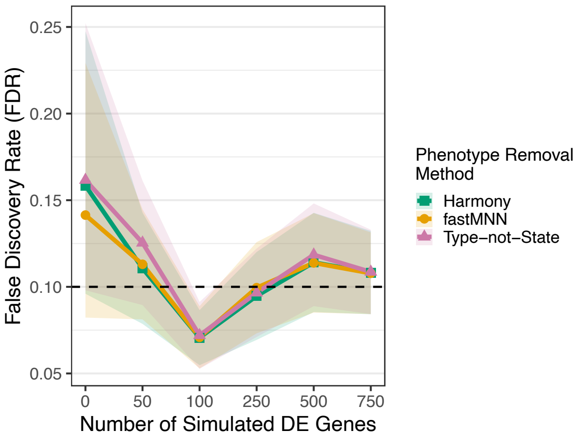


(c) Celltype Aligned DE, lfc = 1.5

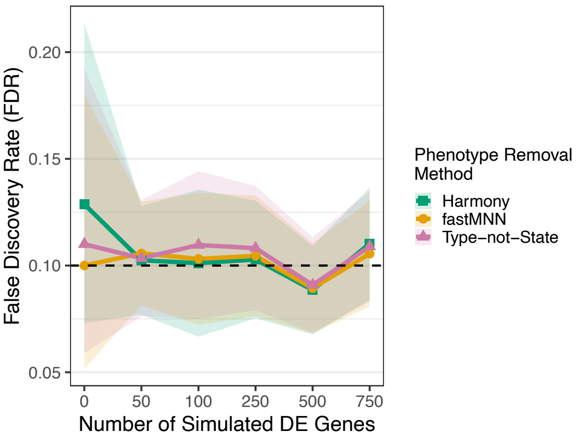


(d) Celltype Non-Aligned DE, lfc = 1.5

Figure S-10: **Comparison of FDR for Different Methods for Phenotype Effect Removal** Plot of FDR as a function of the number of DE genes when clusters are defined using all genes and one of three phenotype removal methods. All simulations were run using harmony corrected null genes for $N_g$.

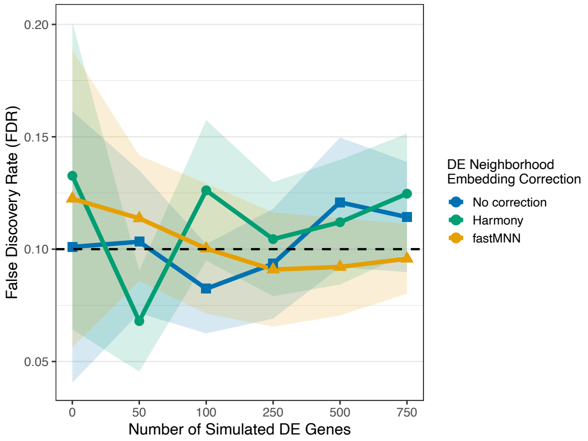


(a) fastMNN correction - Celltype Aligned DE

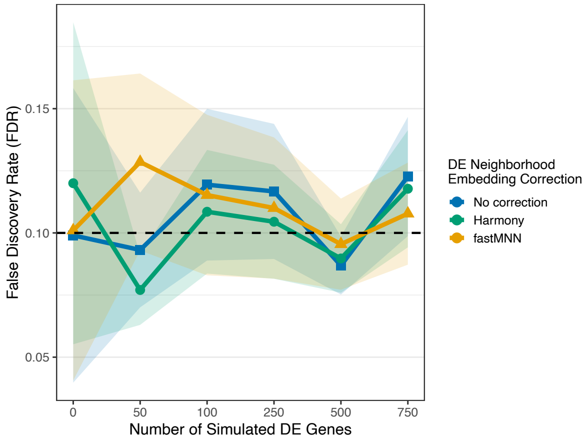


(b) fastMNN correction - Celltype Non-Aligned DE

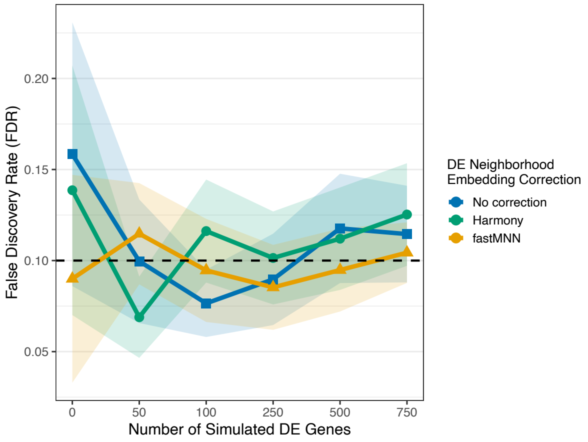


(c) Harmony correction - Celltype Aligned DE

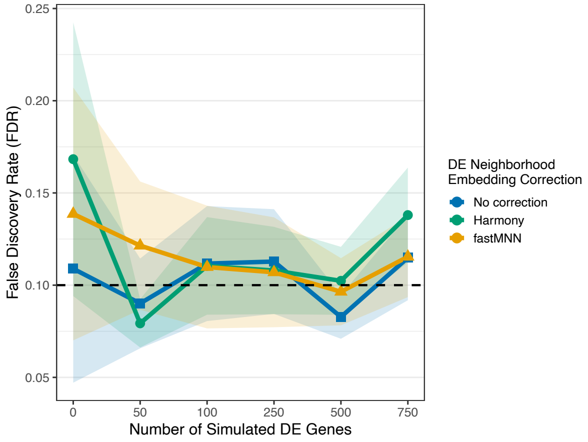


(d) Harmony correction - Celltype Non-Aligned DE

Figure S-11: **FDR control for different choices of gene embeddings for determining** $N_g$ Plotted are the False Discovery Rates plotted against the number of DE genes for different choices of gene embeddings used for defining the true DE neighborhoods $N_g$ (blue = no correction, green=Harmony, and yellow=fastMNN). The top row shows results when the simulated data use fastMNN method for phenotype correction while the bottom row shows the results for applying the Harmony as the phenotype correction. The left column shows when $N_g$ are created to be aligned with celltypes vs non-aligned with celltypes as described in $S1.3$

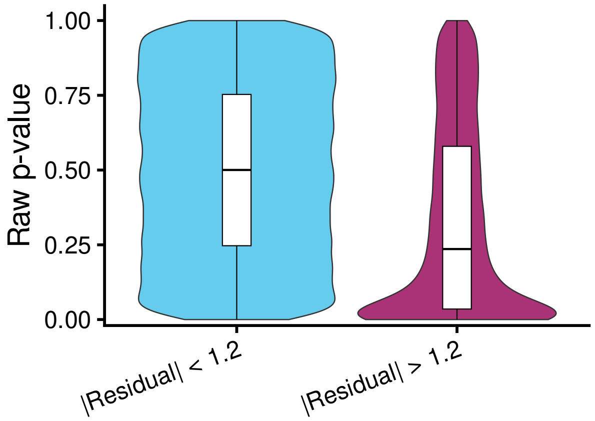


(a) Celltype Aligned $N_g$ - No Correction

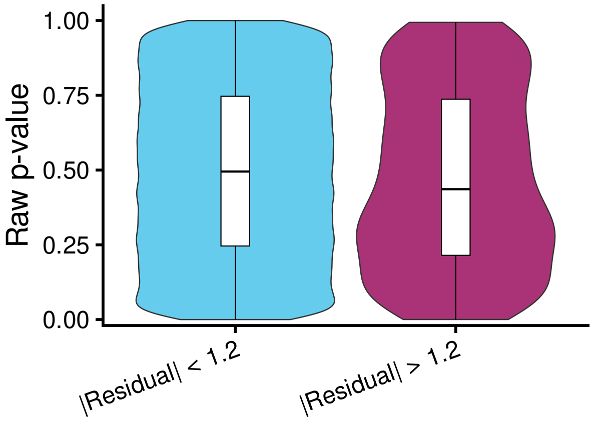


(b) Celltype Aligned $N_g$ - Harmony Correction

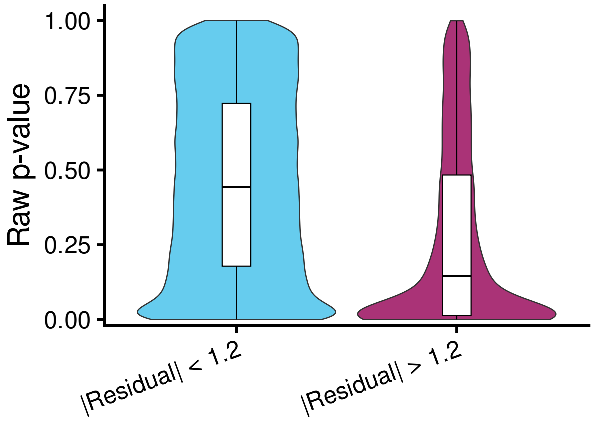


(c) Celltype Non-Aligned $N_g$ - No Correction

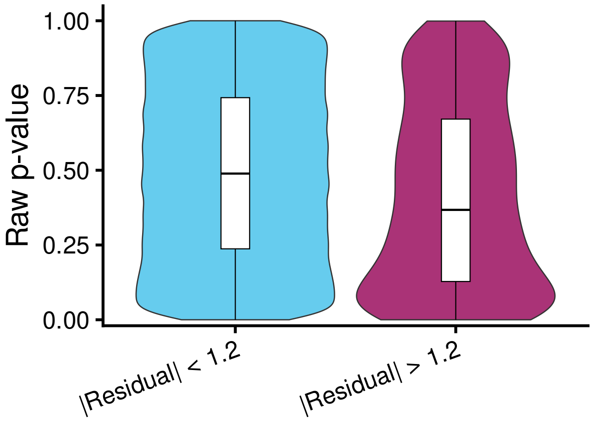


(d) Celltype Non-Aligned $N_g$ - Harmony Correction

Figure S-12: **Negative skew of raw p-value distribution for null hypotheses with cluster imbalance** The violin plots show the distribution of raw p-values for null (non-DE) hypotheses. Distributions of lower imbalance vs higher imbalance hypotheses are next to each other (low sample Pearson means low imbalance). The top row shows the results for simulations with aligned DE regions vs the bottom row shows results for simulations with non-aligned DE regions. Left column shows results with no phenotype corrections vs right column showing (muted) results with Harmony correction. All simulations here were run with 750 DE genes.

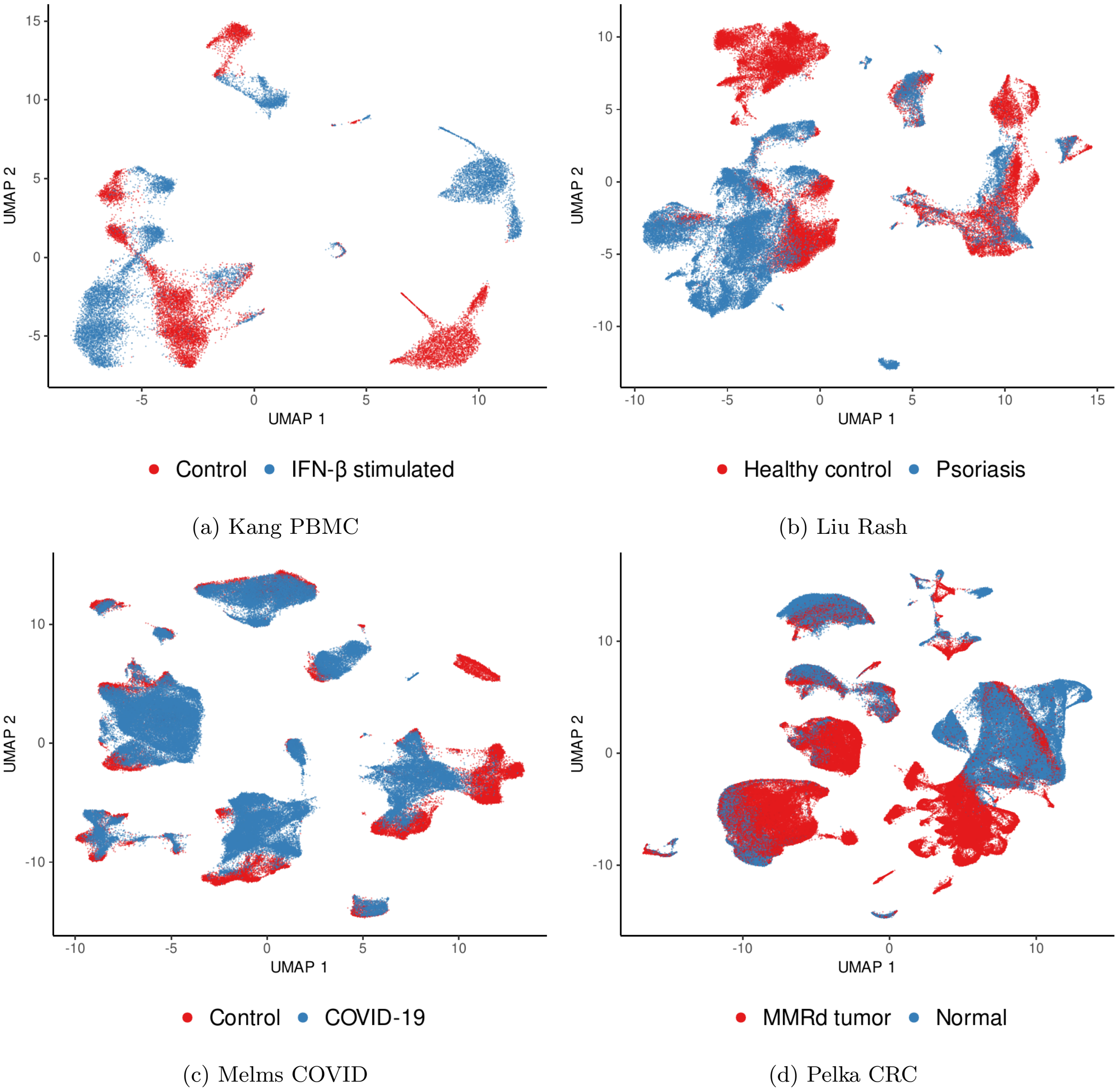


(a) Kang PBMC

(b) Liu Rash

(c) Melms COVID

(d) Pelka CRC

Figure S-13: **UMAP of real datasets with no embedding correction** These UMAPs are shown on real data which are described in *S*4. These follow the standard analysis flow 3.4 by applying UMAP to the top 50 principal components.

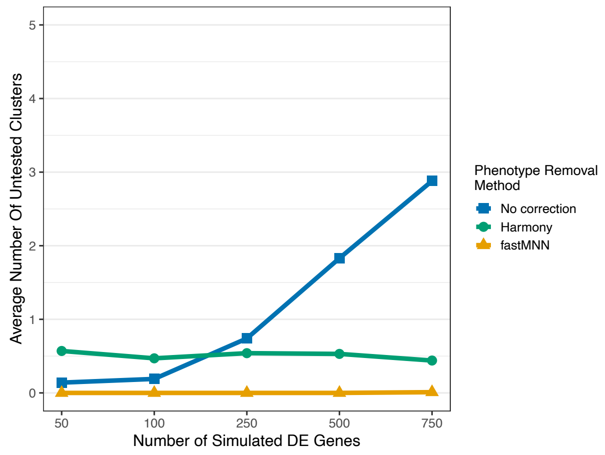


(a) Celltype aligned DE - uncorrected embeddings

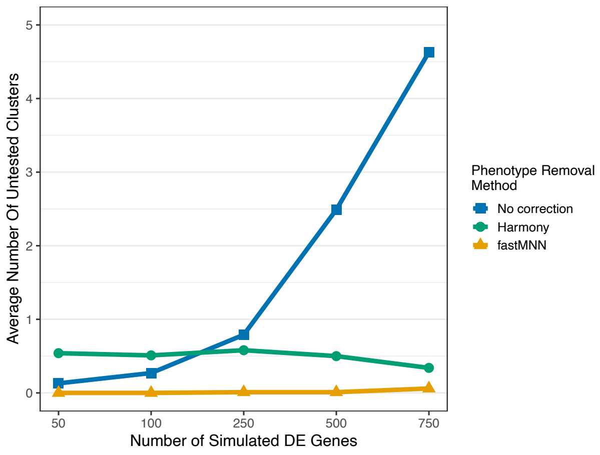


(b) Celltype non-aligned DE - uncorrected embeddings

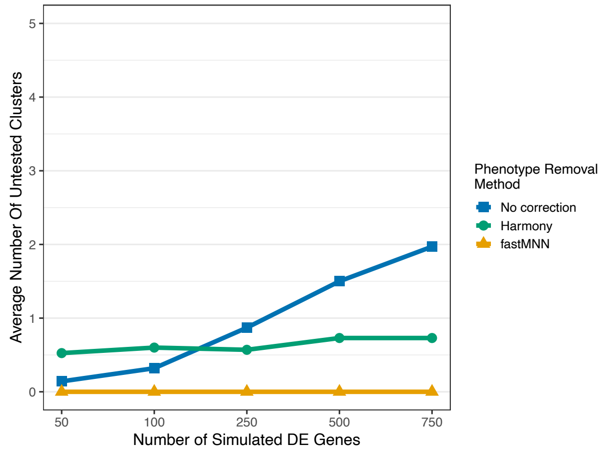


(c) Celltype aligned DE - Harmony embeddings

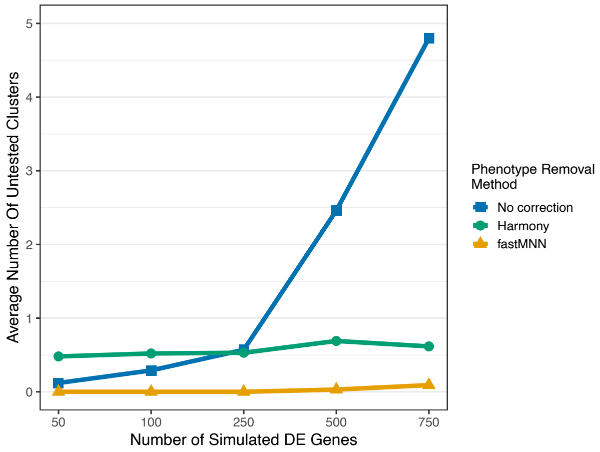


(d) Celltype non-aligned DE - Harmony embeddings

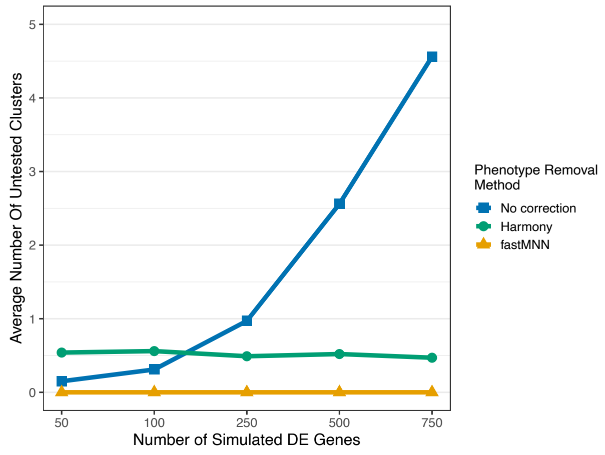


(e) Celltype aligned DE - FastMNN embeddings

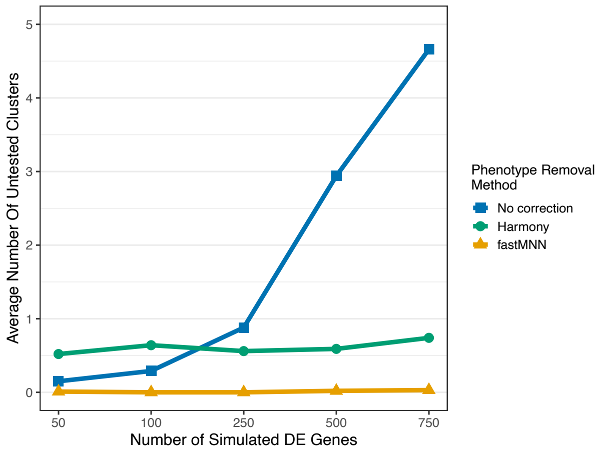


(f) Celltype non-aligned DE - FastMNN embeddings

Figure S-14: **Average number of clusters filtered out for testing per simulation** Each plot shows the per-simulation average of the number of found clusters that are filtered out, stratified by the number of DE genes simulated in the simulation on the x-axis, and by phenotype removal method. Each plot shows the data for a different strategy and embedding correction choice for creating $N_g$.

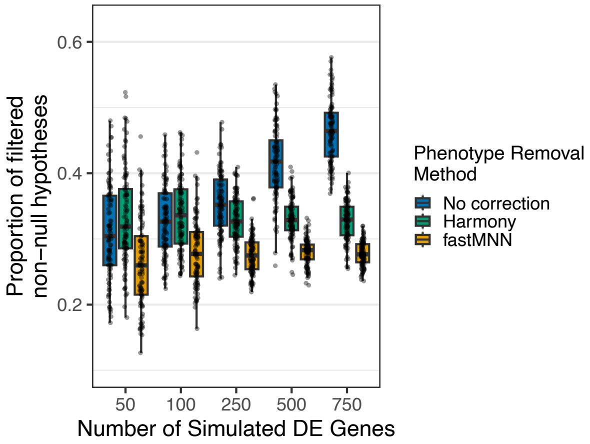


(a) Celltype aligned DE - uncorrected embeddings

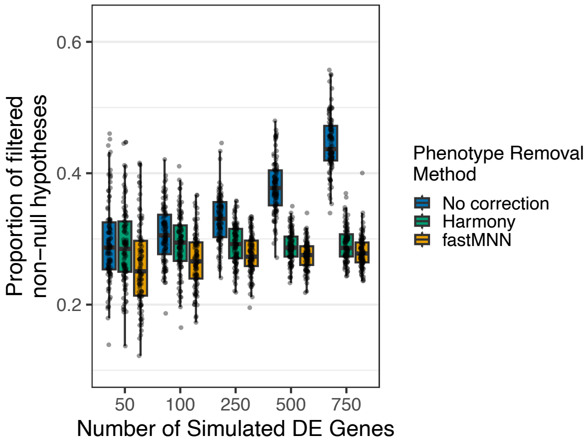


(b) Celltype non-aligned DE - uncorrected embeddings

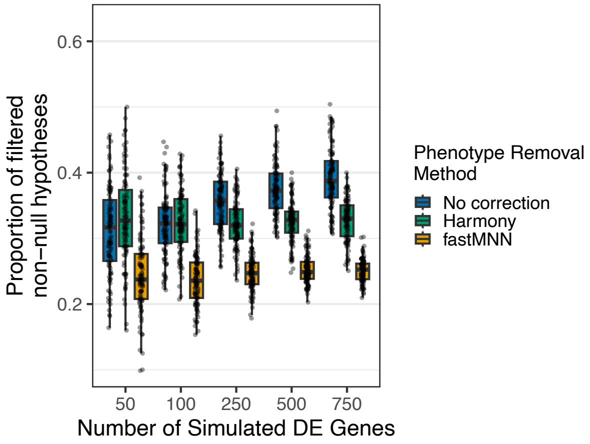


(c) Celltype aligned DE - Harmony embeddings

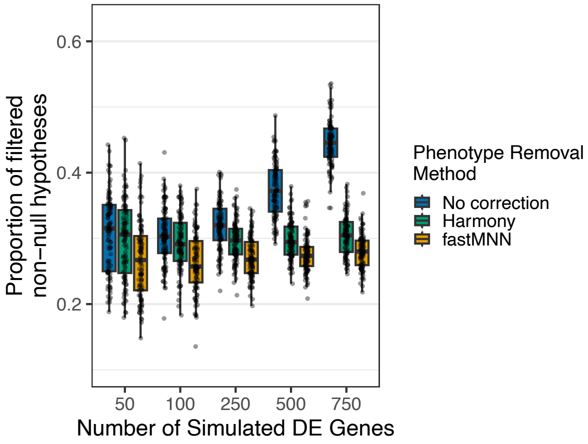


(d) Celltype non-aligned DE - Harmony embeddings

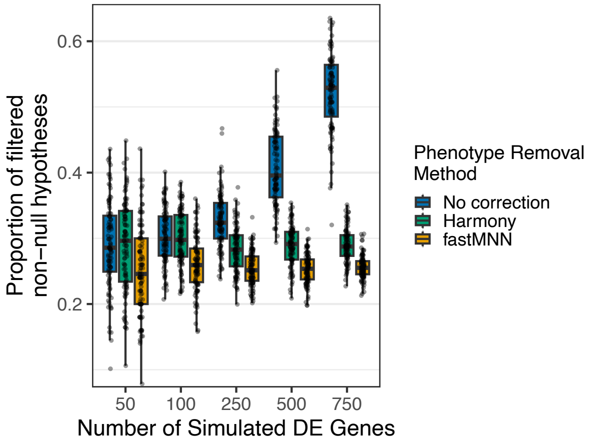


(e) Celltype aligned DE - FastMNN embeddings

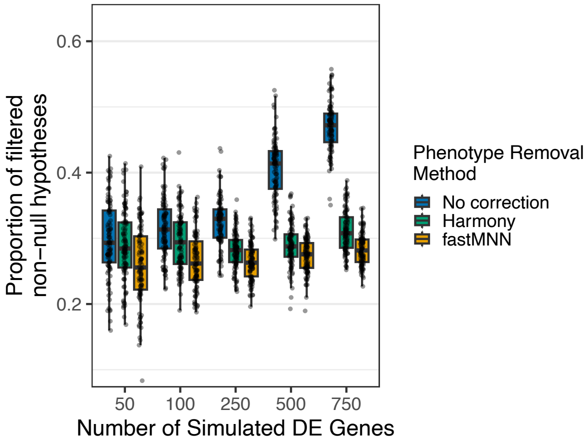


(f) Celltype non-aligned DE - FastMNN embeddings

Figure S-15: **Proportion of non-null hypothesis filtered** Each plot shows the per-simulation distribution of the proportion of non-null hypotheses that are filtered out, stratified by the number of DE genes simulated in the simulation on the x-axis, and by phenotype removal method. Each plot shows the data for a different strategy and embedding correction choice for creating $N_g$.

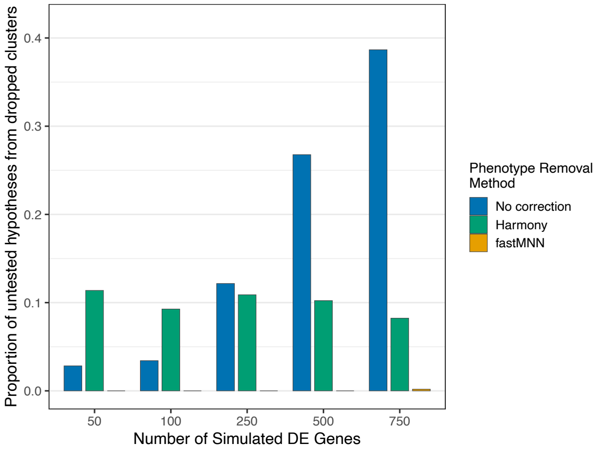


(a) Celltype aligned DE - uncorrected embeddings

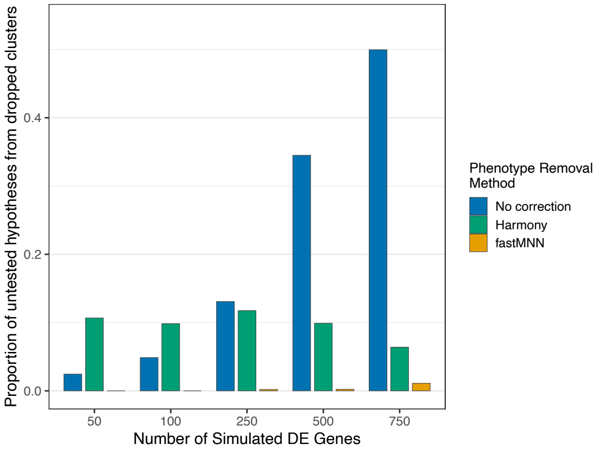


(b) Celltype non-aligned DE - uncorrected embeddings

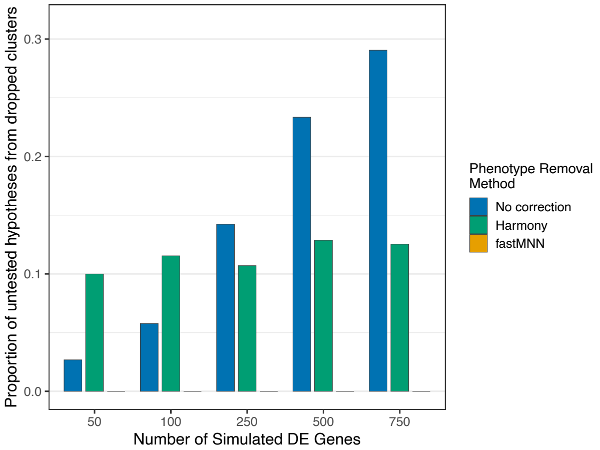


(c) Celltype aligned DE - Harmony embeddings

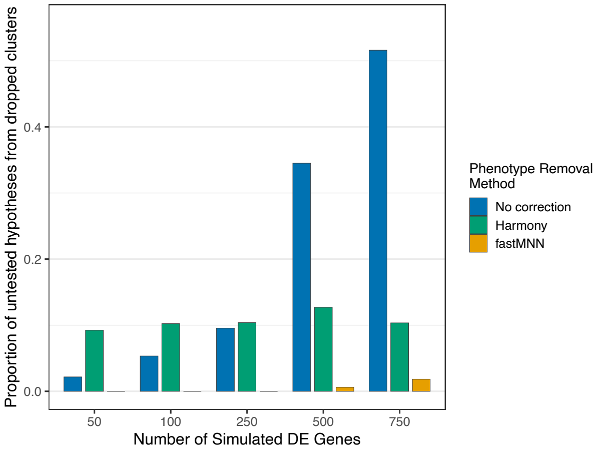


(d) Celltype non-aligned DE - Harmony embeddings

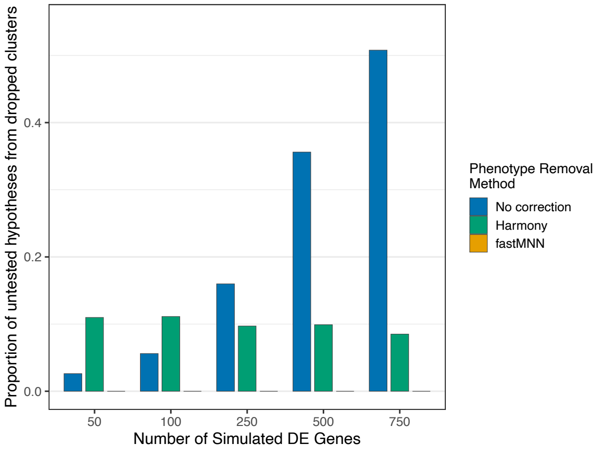


(e) Celltype aligned DE - FastMNN embeddings

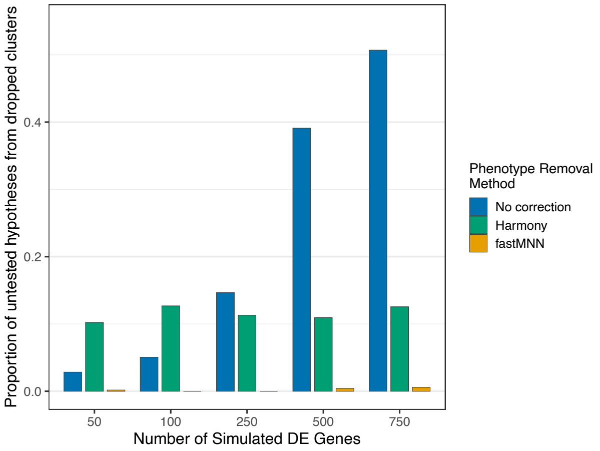


(f) Celltype non-aligned DE - FastMNN embeddings

Figure S-16: **Proportion of filtered non-null hypotheses in fully untested clusters** Each plot shows the proportion of untested non-null hypotheses where all hypotheses in the cluster were untested, stratified by the number of DE genes simulated in the simulation on the x-axis, and by phenotype removal method. Each plot shows the data for a different strategy and embedding correction choice for creating $N_g$.

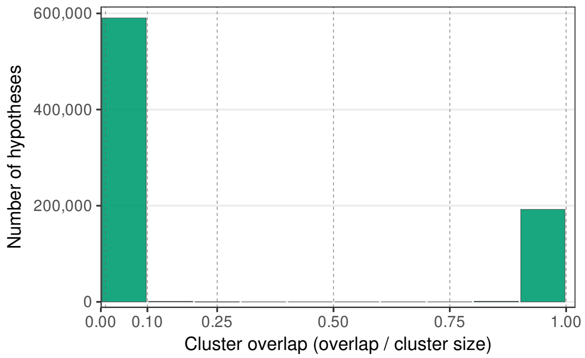


(a) Celltype aligned DE - all overlaps

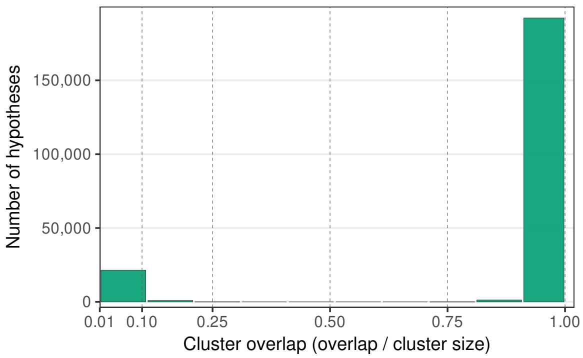


(b) Celltype aligned DE - overlaps > 0.01

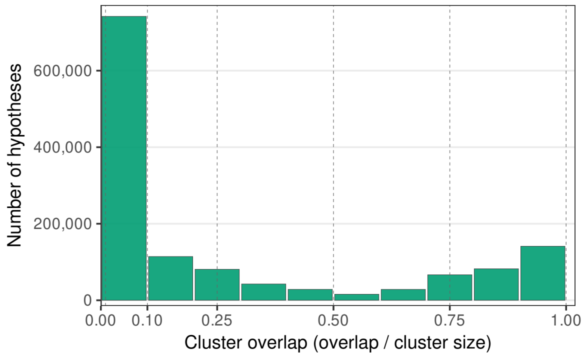


(c) Celltype non-aligned DE - all overlaps

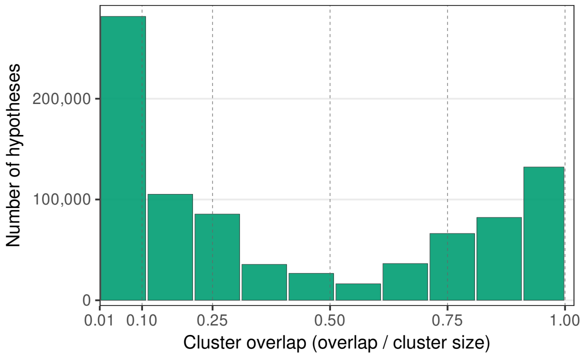


(d) Celltype non-aligned DE - overlaps > 0.01

Figure S-17: **Comparison of overlap distribution when counting very small overlaps** Each graph shows the distribution of overlap positive hypotheses across all simulations for a given combination of $N_g$ creation strategy. The histogram aggregates all simulations that employed the chosen $N_g$ creation strategy and used the phenotype removal method that matched the sample correction for the embeddings used for $N_g$. The left column shows the histogram across all hypotheses while the right column excludes hypotheses below 0.01 using the cluster overlap definition.

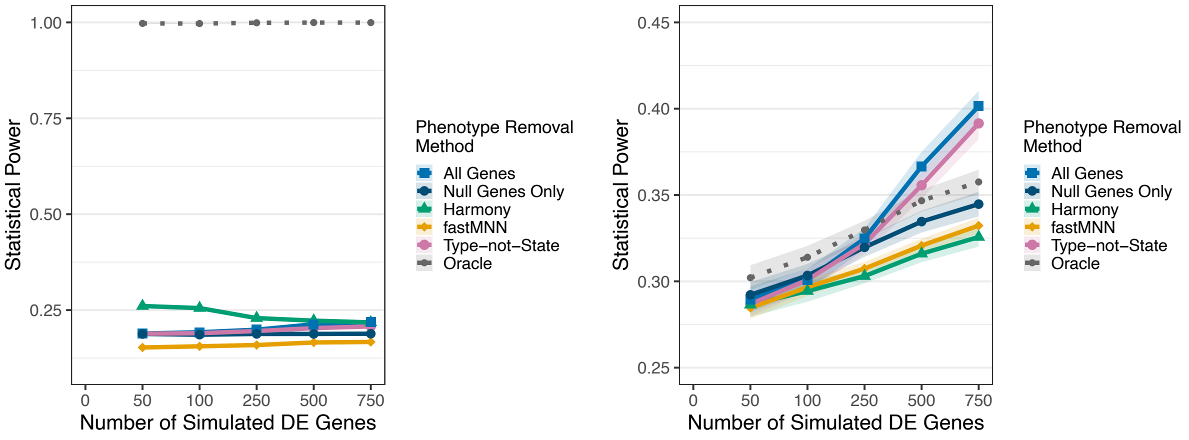


(a) Celltype Aligned DE

(b) Celltype Non-Aligned DE

Figure S-18: **Comparison of Power to find *any* overlap with DE cells,** $b = 0.0$**.** The the power to find DE neighborhoods (y-axis) as a function of the number of DE genes (x-axis), with different phentoype-correction strategies plotted in different colors indicated in the legend. The dark grey dotted line corresponds to the “oracle”, where the true clusters are used for testing. True DE neighborhoods for all simulations are chosen from embeddings based on null genes with Harmony correction with the average log fold change of 3.

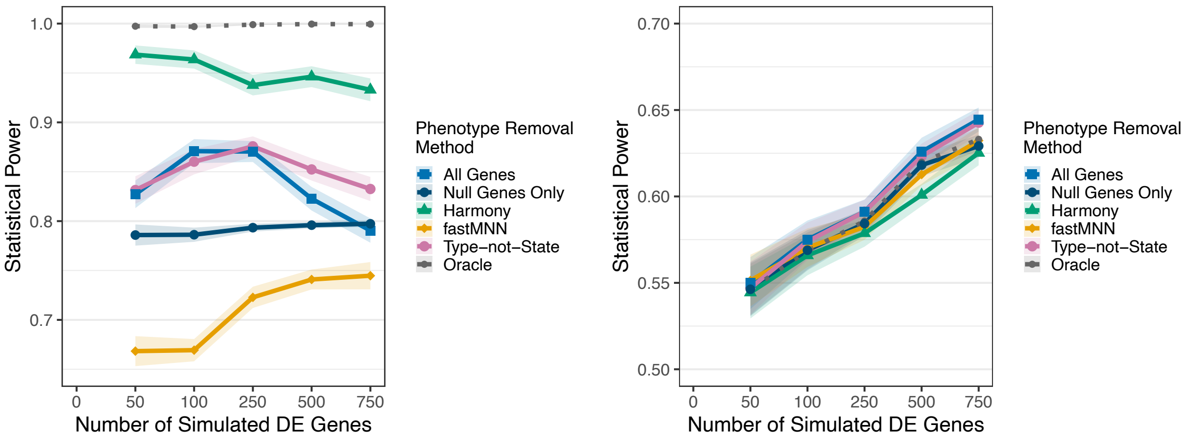


(a) Celltype aligned DE

(b) Celltype non-aligned DE

Figure S-19: **Comparison of Power to find DE if restricted to clusters with overlap with** $N_g$ **of at least** $b = 0.01$**.** The the power to find DE neighborhoods (y-axis) as a function of the number of DE genes (x-axis), with different phentoype-correction strategies plotted in different colors indicated in the legend. The dark grey dotted line corresponds to the “oracle”, where the true clusters are used for testing. True DE neighborhoods for all simulations are chosen from embeddings based on null genes with Harmony correction with the average log fold change of 3.

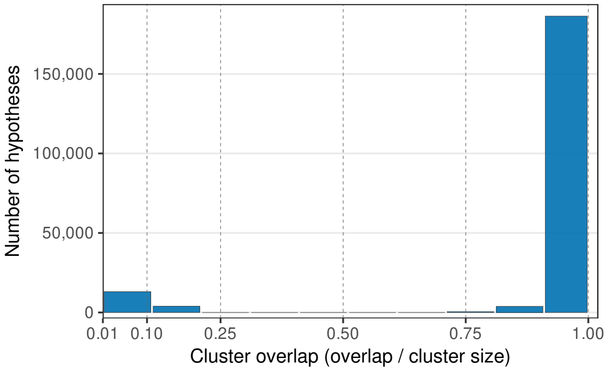


(a) Celltype aligned - uncorrected

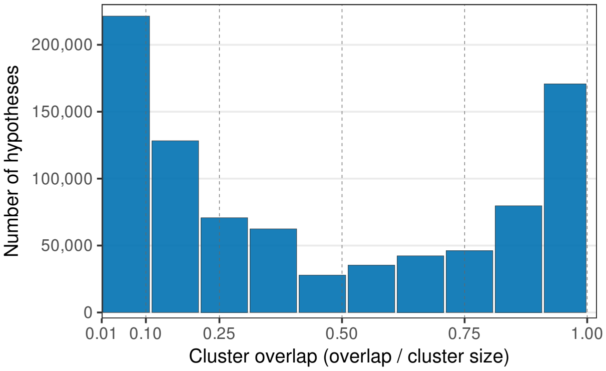


(b) Celltype non-aligned - No Correction

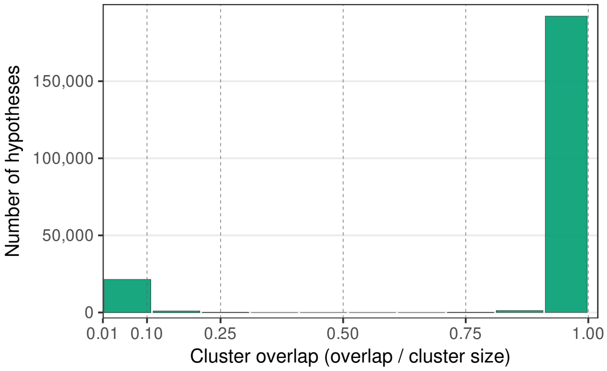


(c) Celltype aligned - Harmony corrected

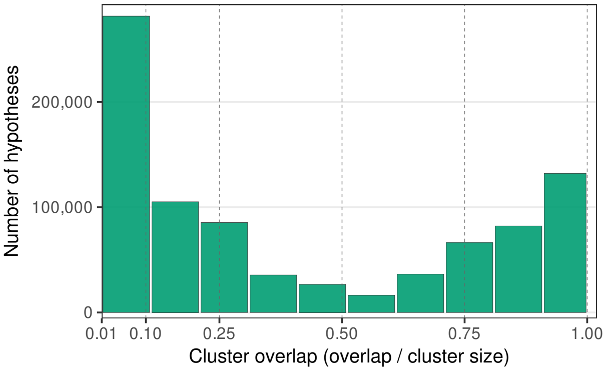


(d) Celltype non-aligned - Harmony corrected

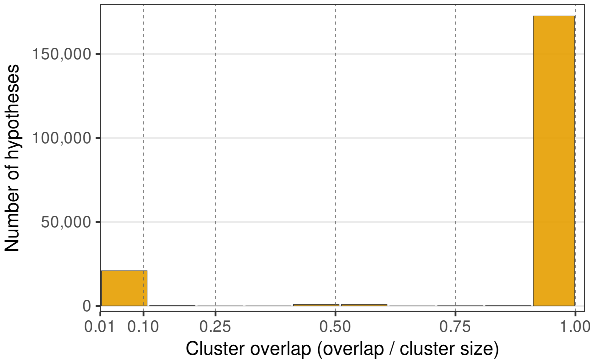


(e) Celltype aligned - FastMNN corrected

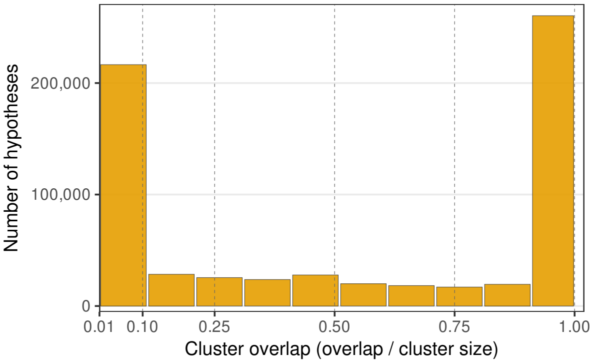


(f) Celltype non-aligned - FastMNN corrected

Figure S-20: **Distributions of non-null hypotheses across all $N_g$ creation strategies** Each graph shows the distribution of hypotheses (limited to hypotheses with overlaps above 0.01) across all simulations for a given combination of $N_g$ creation strategy. The histogram aggregates all simulations that employed the chosen $N_g$ creation strategy and used the phenotype removal method that matched the sample correction for the embeddings used for $N_g$.

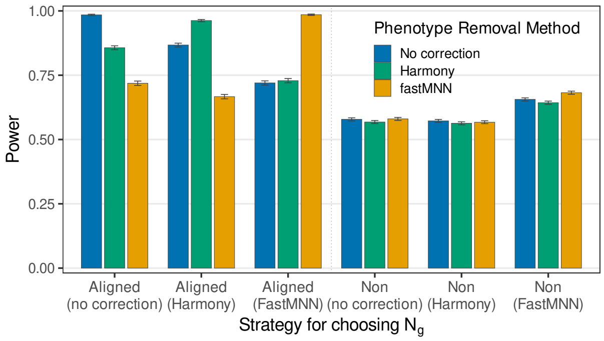


(a) 100 DE genes

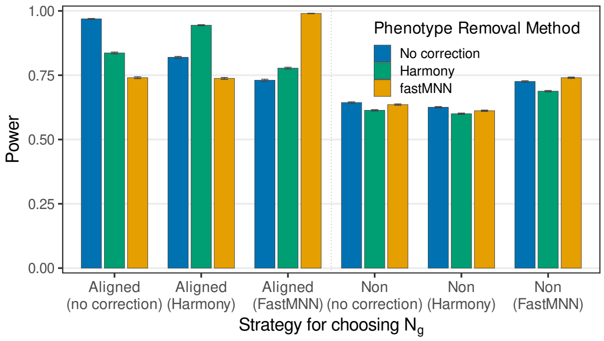


(b) 500 DE genes

Figure S-21: **Power for different pairings of embeddings for generation of $N_g$ and clustering of data** Plots the average power of each phenotype remval method (each colored bar),method for generating $N_g$ (”Aligned”=celltype-aligned vs ”Non”=celltype-nonaligned) and method for sample correction of null genes (in parentheses of $x$-label).

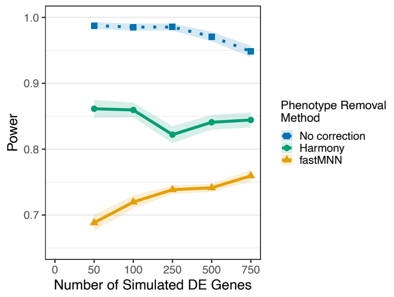


(a) No Sample Correction

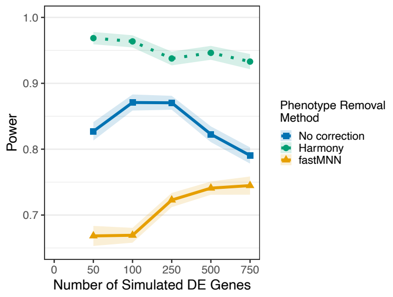


(b) Harmony Correction

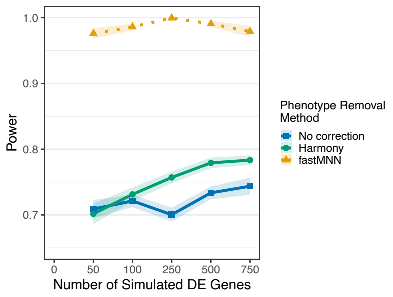


(c) fastMNN Correction

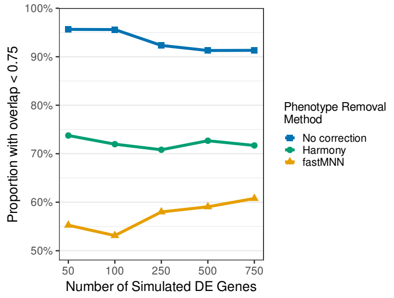


(d) No Sample Correction

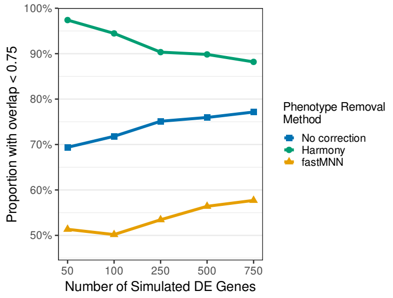


(e) Harmony Correction

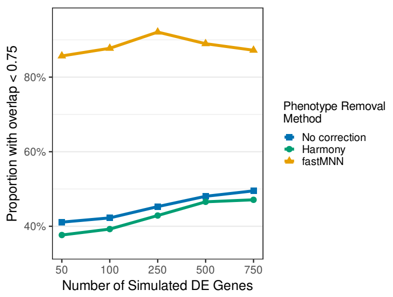


(f) fastMNN Correction

Figure S-22: **Power and high-overlap proportions for celltype aligned generation of $N_g$ with increasing DE genes** On the top row, the plot shows the average power for simulations aggregated by number of DE genes (on the x-axis), phenotype removal method (color scale) for a given $N_g$ embedding correction method. The bottom row shows the proportion of hypotheses that are above 0.75 cluster overlap, aggregating over the same set of simulations as for the top row. Each column represents a different $N_g$ embedding correction method. All plots celltype aligned DE.

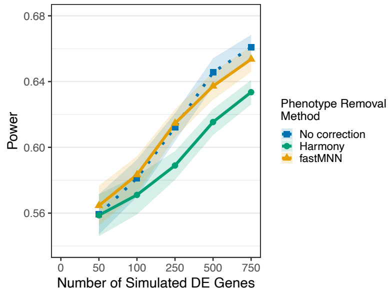


(a) No Sample Correction

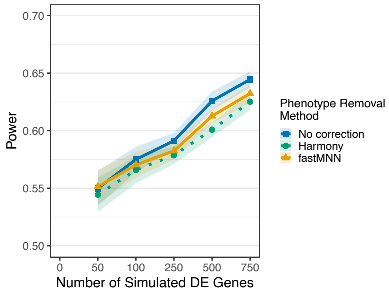


(b) Harmony Correction

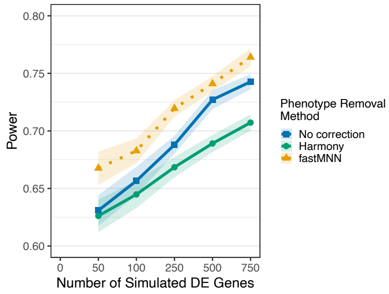


(c) fastMNN Correction

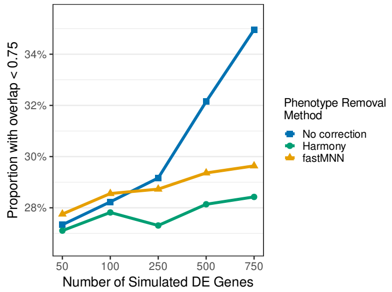


(d) No Sample Correction

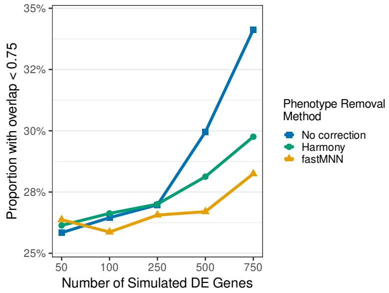


(e) Harmony Correction

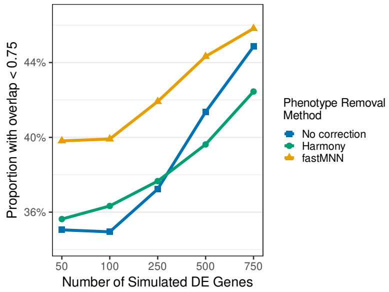


(f) FastMNN Correction

Figure S-23: **Power and high-overlap proportions for celltype non-aligned generation of $N_g$ with increasing DE genes** On the top row, the plot shows the average power for simulations aggregated by number of DE genes (on the x-axis), phenotype removal method (color scale) for a given $N_g$ embedding correction method. The bottom row shows the proportion of hypotheses that are above 0.75 cluster overlap, aggregating over the same set of simulations as for the top row. Each column represents a different $N_g$ embedding correction method.

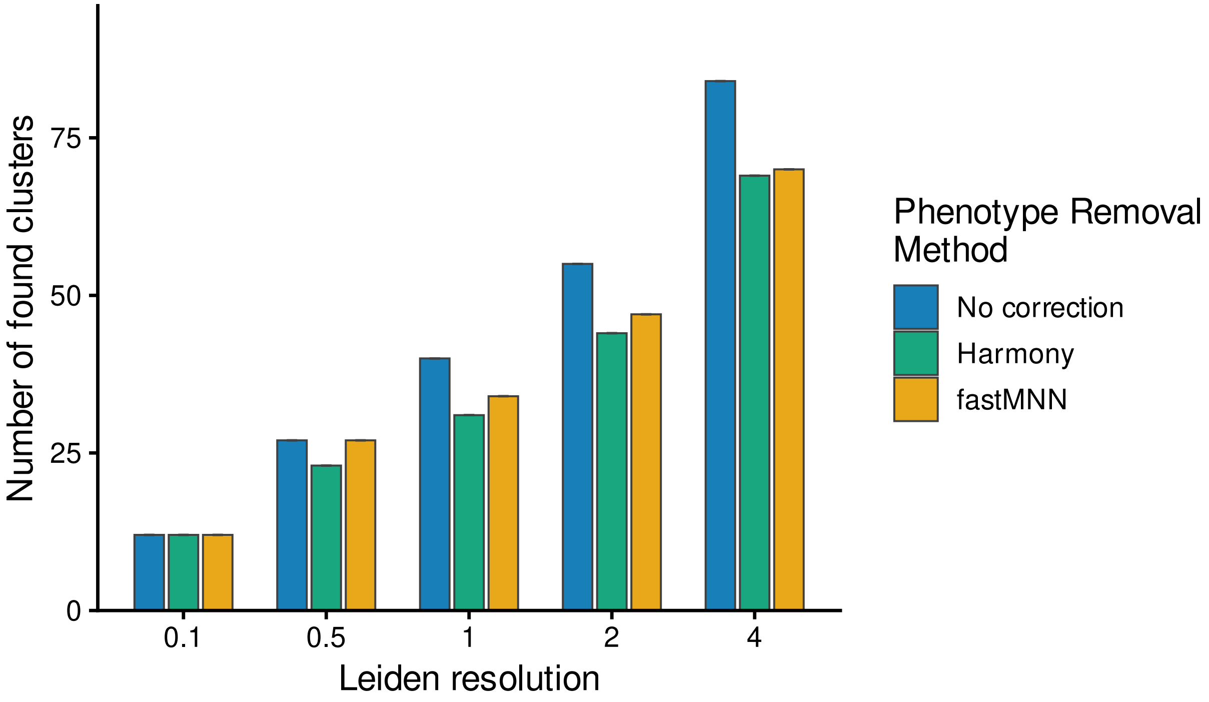


Figure S-24: Total Number Of Clusters

Figure S-25: **Number of found clusters on null genes with varying cluster resolutions** Number of found clusters associated with each choice of Leiden resolution parameter, demonstrating progressively finer partitions. These are run on null and real data, with no simulated DE genes.

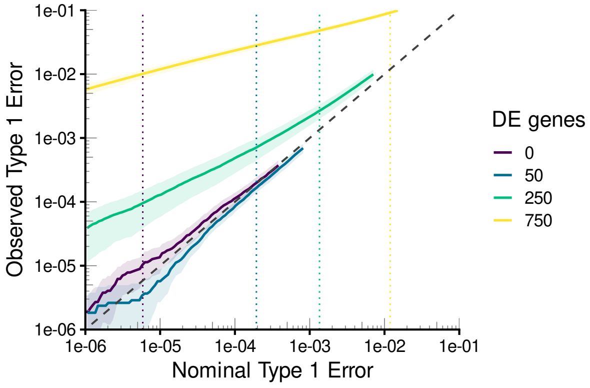


(a) No Correction, Resolution 0.1

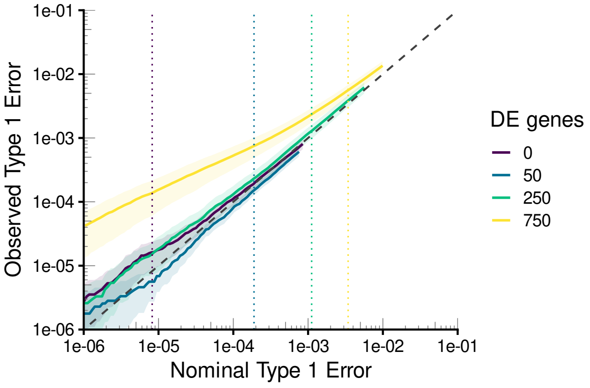


(b) Harmony, Res 0.1

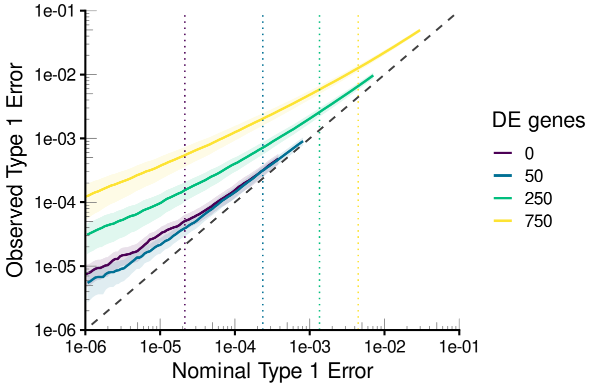


(c) No Correction, Res 1

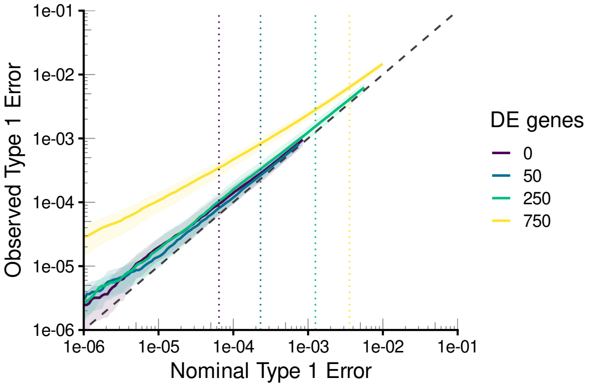


(d) Harmony, Res 1

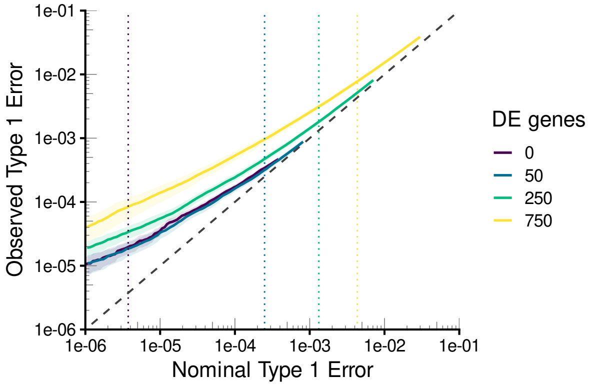


(e) No Correction, Res 4

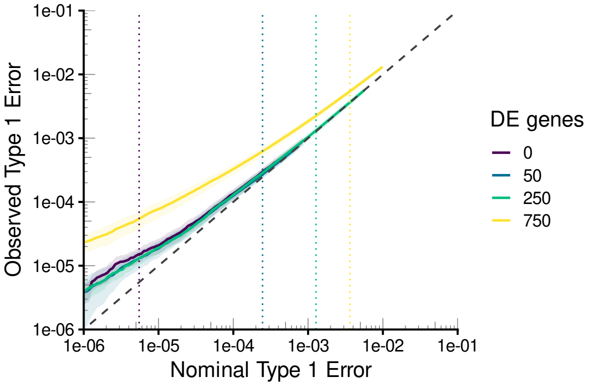


(f) Harmony, Res 4

Figure S-26: **Nominal vs Observed Type 1 Error for different resolutions and phenotype corrections using celltype non-aligned DE** Each graph shows the average observed type 1 error across simulations against the nominal type 1 error implied by the raw p-values. Each line and plot shows a different set of simulations across which the average is taken. The left column shows simulations with no phenotype corrections, while the right column show harmony corrected simulations. The data is further divided by resolution, with resolutions 0.1, 1, 4 being used for the top, middle and bottom row respectively. Finally, simulations are further stratified by number of DE genes within each plot, with one line for each level. All simulations come from a celltype non-aligned $N_g$ with harmony corrected embeddings. Average cutoffs across simulations for nominal type 1 error corresponding to FDR of 0.1 are shown as vertical dashed lines of the corresponding color.

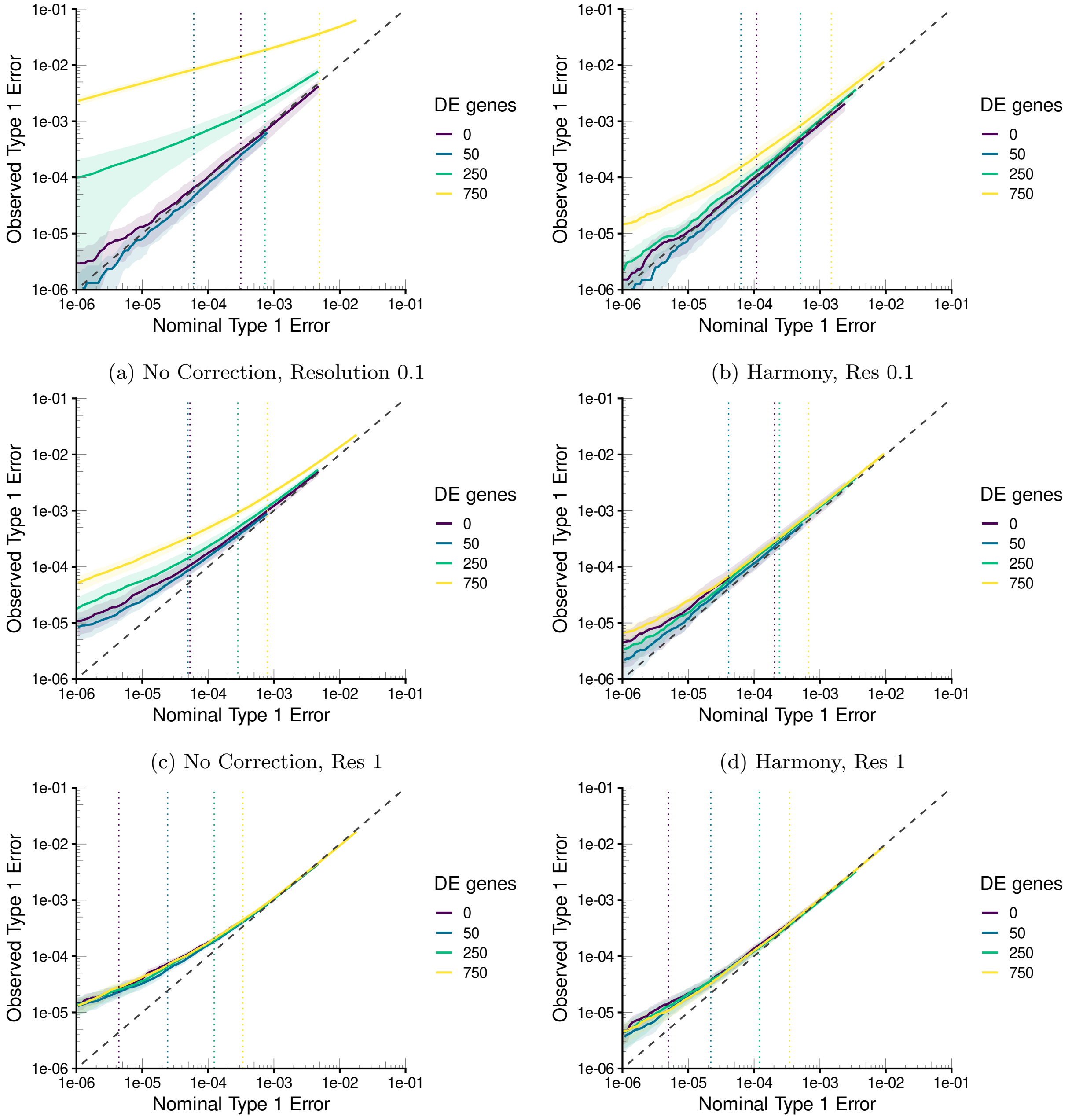


(a) No Correction, Resolution 0.1

(b) Harmony, Res 0.1

(c) No Correction, Res 1

(d) Harmony, Res 1

(e) No Correction, Res 4

(f) Harmony, Res 4

Figure S-27: **Nominal vs Observed Type 1 Error for different resolutions and phenotype corrections using celltype aligned DE** Each graph shows the average observed type 1 error across simulations against the nominal type 1 error implied by the raw p-values. Each line and plot shows a different set of simulations across which the average is taken. The left column shows simulations with no phenotype corrections, while the right column show harmony corrected simulations. The data is further divided by resolution, with resolutions 0.1, 1, 4 being used for the top, middle and bottom row respectively. Finally, simulations are further stratified by number of DE genes within each plot, with one line for each level. All simulations come from a celltype aligned $N_g$ with harmony corrected embeddings. Average cutoffs across simulations for nominal type 1 error corresponding to FDR of 0.1 are shown as vertical dashed lines of the corresponding color.

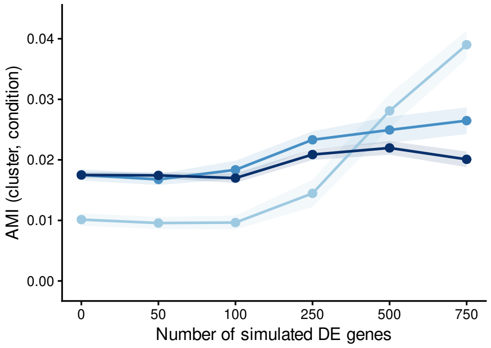


(a) AMI, No Correction

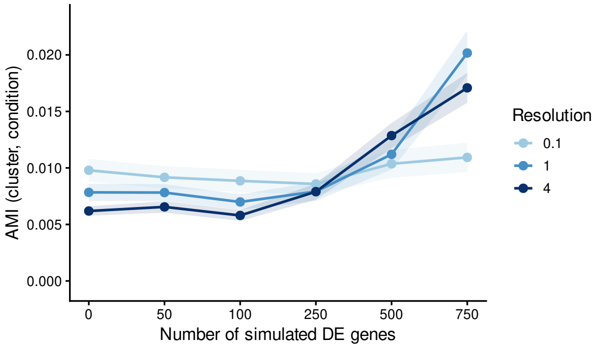


(b) AMI, Harmony Correction

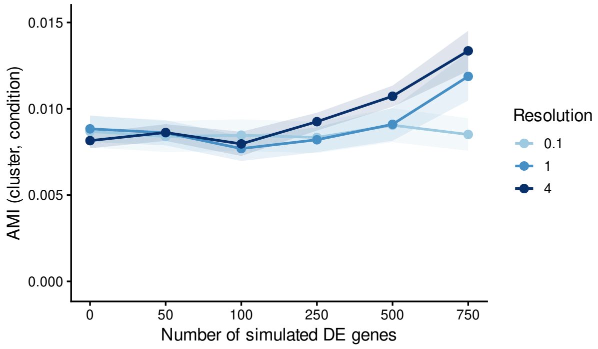


(c) AMI, fastMNN Correction

Figure S-28: **Imbalance measures across resolutions with celltype non-aligned DE** Each plot shows the average value of Adjusted Mutual Information (AMI) across simulations stratified by number of DE genes (x-axis), resolution (color) and phenotype correction method (row). The AMI is computed by the contingency table of cluster by phenotype of cells, including only clusters that are not completely dropped

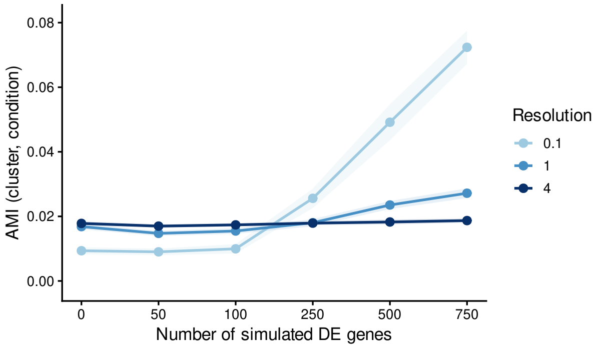


(a) AMI, No Correction

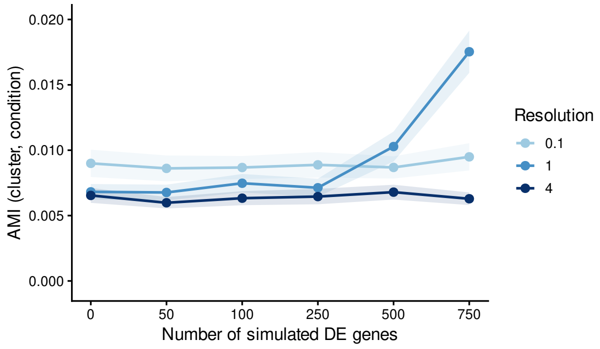


(b) AMI, Harmony Correction

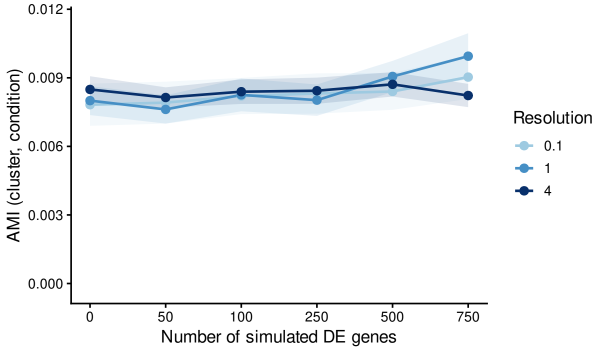


(c) AMI, fastMNN Correction

Figure S-29: **Imbalance measures across resolutions with celltype aligned DE** Each plot shows the average value of Adjusted Mutual Information (AMI) across simulations stratified by number of DE genes (x-axis), resolution (color) and phenotype correction method (row).The AMI is computed by the contingency table of cluster by phenotype of cells, including only clusters that are not completely dropped

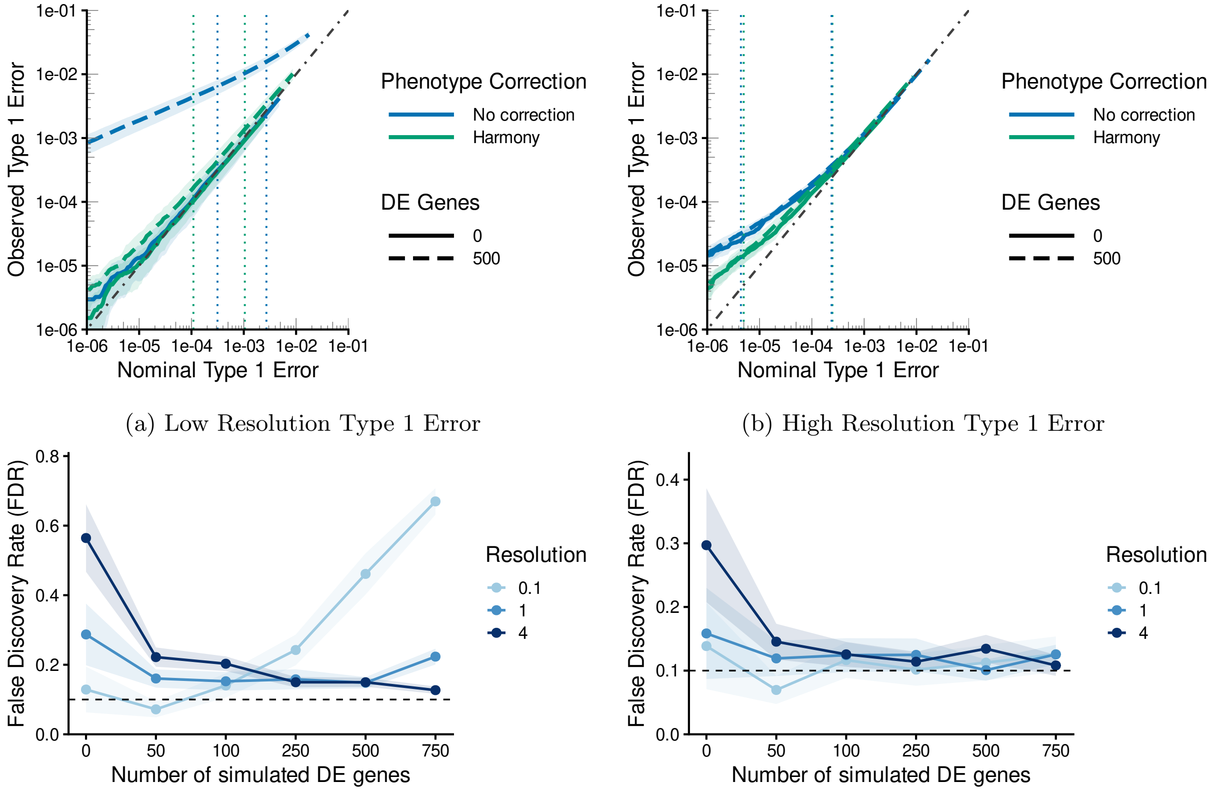


(a) Low Resolution Type 1 Error

(b) High Resolution Type 1 Error

(c) FDR for Uncorrected Phenotype

(d) FDR for Harmony Corrected Phenotype

Figure S-30: **Loss of FDR and Type 1 Error Control with varying resolutions and number of DE genes with celltype aligned DE** The top row shows compares nominal type 1 error rates with observed type 1 errors when using raw, uncorrected p-values for simulations at a given number of DE genes (shown by line type) and either with a given phenotype correction method (shown by color). (a) shows results for the low resolution case (leiden resolution parameter of 0.1) while (b) shows results for high resolution (leiden resolution parameter of 4). The bottom row shows the FDR for different resolutions (shown as differently colored lines) with increasing number of DE genes. (c) shows the results when there are no corrections to phenotype effect while (d) shows results when Harmony correction is applied for phenotype corrections. All plots were on simulations run with$N_g$ were chosen to be celltype-aligned and null genes were Harmony corrected.

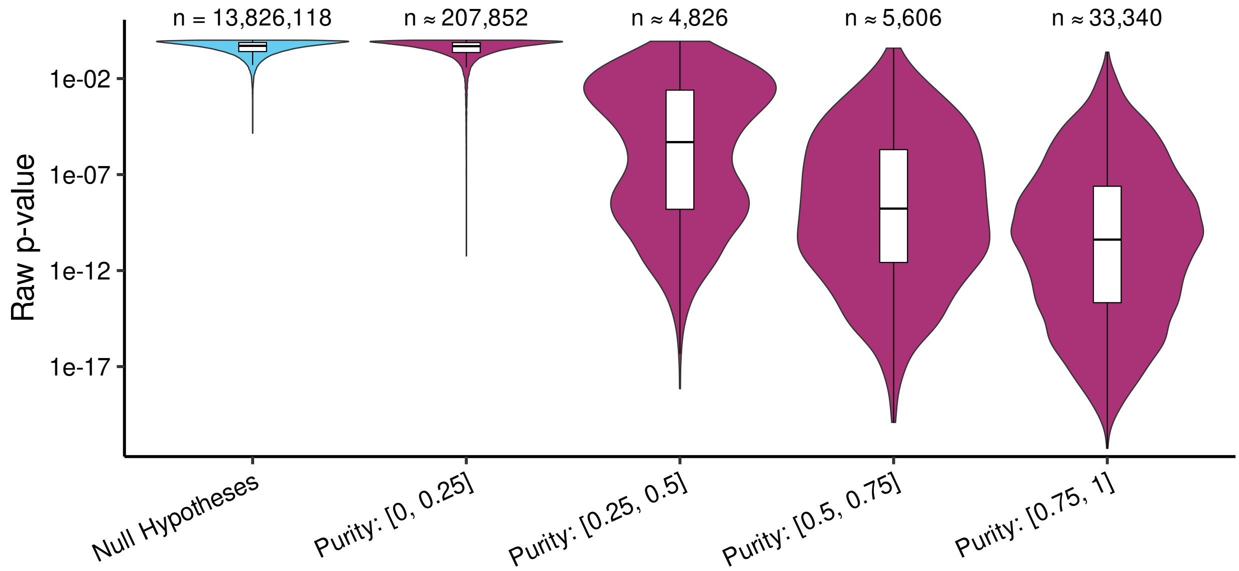


(a) Celltype Aligned DE

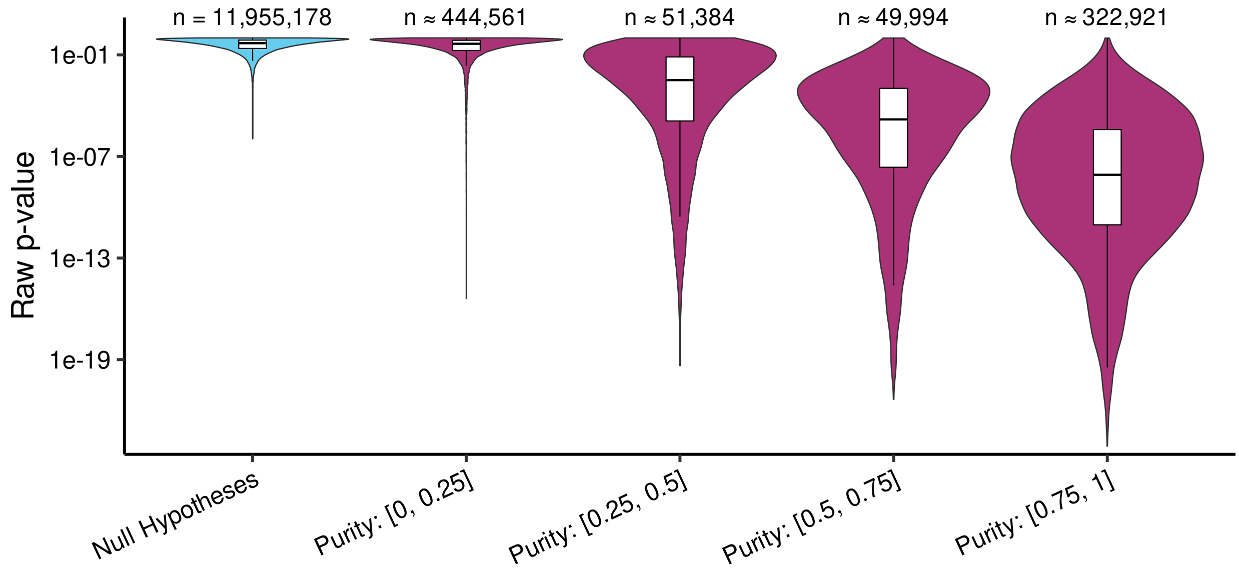


(b) Celltype Non-Aligned DE

Figure S-31: **Strength of DE signal for simulated genes** Violin plots show the distribution of raw p-values stratified by increasing DE purity, starting with the null hypotheses. Approximate number of hypotheses in each bucket across all simulations. The top row shows results for simulations with aligned DE neighborhoods while the bottom rows is for simulations with non-aligned DE neighborhoods. All simulations here were made with 750 simulated DE genes, Leiden resolution of 4, using harmony corrected embeddings for $N_g$ and phenotype correction using Harmony. Up to 50,000 values were subsampled for estimating the distribution.

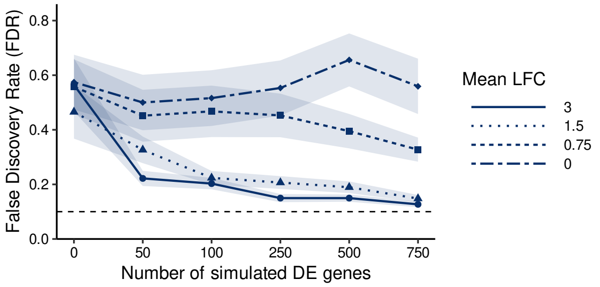


(a) Celltype Aligned DE, No Correction

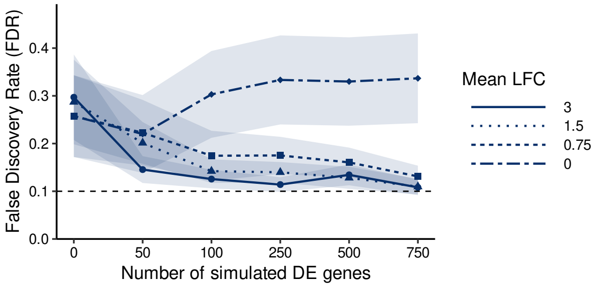


(b) Celltype Aligned DE, Harmony Correction

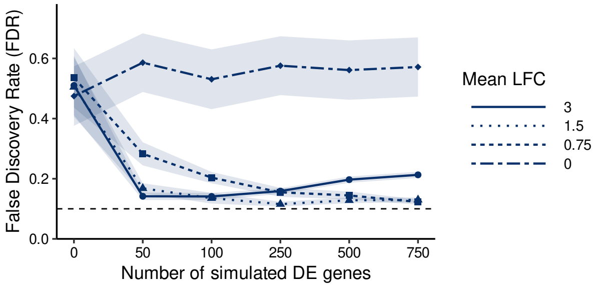


(c) Celltype Non-Aligned DE, No Correction

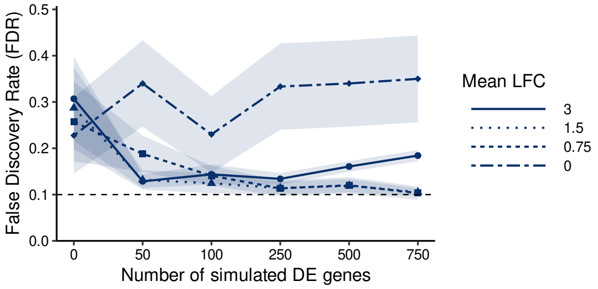


(d) Celltype Non-Aligned DE, Harmony Correction

Figure S-32: **Effect of lower log fold change on FDR at high resolutions** These figures show FDR averaged across simulations that are run with high resolution (leiden resolution of 4) to show the difference between the behavior of the FDR with increasing number of simulated DE genes when those DE genes have different log fold changes (different line-types). Each column of plots show different phenotype removal methods while the rows show results for simulations with celltype aligned DE neighborhoods (top) and celltype non-aligned DE neighborhoods (bottom)

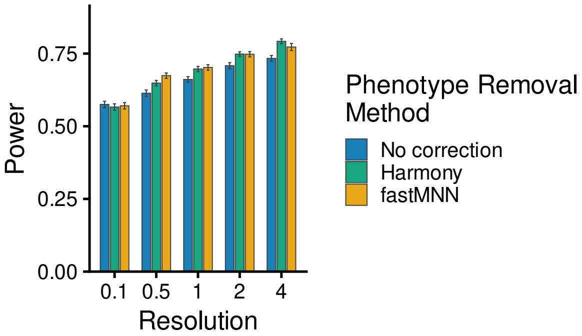


(a) Power with 100 DE genes

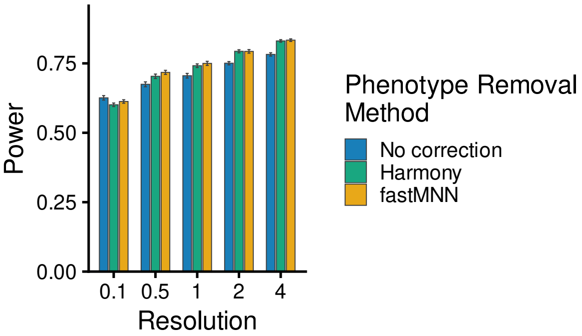


(b) Power with 500 DE genes

Figure S-33: **Power with varying resolutions for celltype non-aligned DE** Power is shown with increasingly higher resolutions, with different phenotype removal methods shown as differently colored bars. (a) is for simulations with 100 DE genes while (b) is for those with 500 DE genes. $N_g$ were chosen using celltype non-aligned DE and Harmony sample-correction for the null genes.

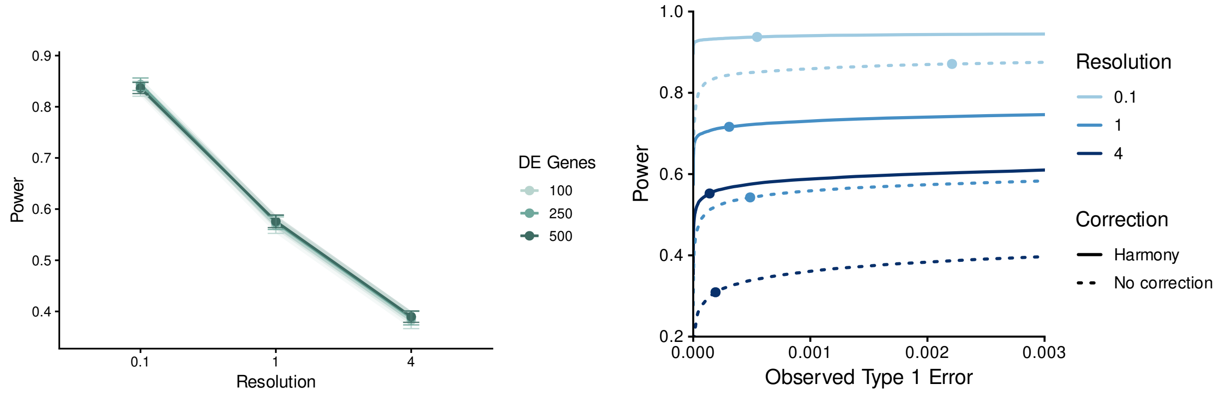


(a) Power

(b) ROC curves

Figure S-34: **Power and ROC curves when the varying resolutions lead to clusters that are mostly subsets of the true $N_g$ DE neighborhoods.** The above plots replicate the plots of Figure 7 for the simulation setting where the DE neighborhoods are chosen to align with celltypes. This means that the true $N_g$ are themselves Leiden clusters *with resolution 0.1* on the null genes. Thus when higher Leiden resolutions are applied to the simulated data, the found clusters $\hat{C}_k$ are generally close to being pure subsets of the true $N_g$. (a) shows power (x-axis) with increasingly higher resolutions (y-axis). Different number of DE genes are shown as differently colored lines. The result is averaged across all simulations regardless of phenotype removal method. (b) shows the power (y-axis) vs observed type 1 error (x-axis) for varying raw p-value cutoffs. Different resolutions are shown with different lines; all have 250 DE genes. The plot displays results with and without Harmony phenotype correction (solid vs dotted respectively). Solid points on each curve are located by determining the raw p-value that would match a BH threshold of 0.1, averaging this across simulations, and then plotting the value of Type 1 error and power that corresponds to this raw p-value cutoff.I changed this because didn't understand. Check if correct . For both plots, $N_g$ were on Harmony sample-correction embeddings on the null genes.

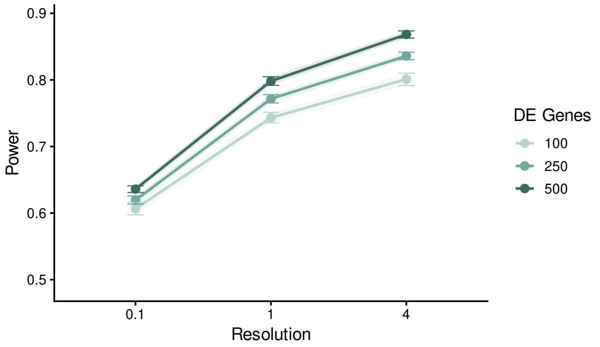


(a) Large clusters, $k = 3$

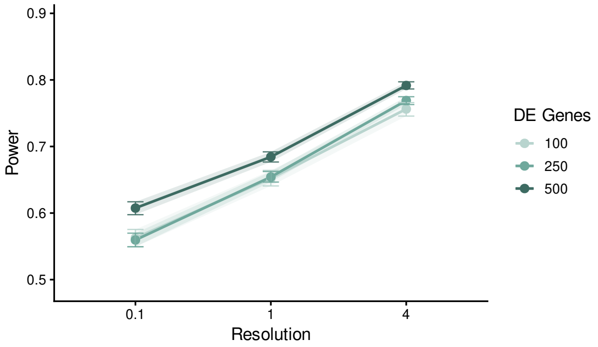


(b) Medium clusters, $k = 10$

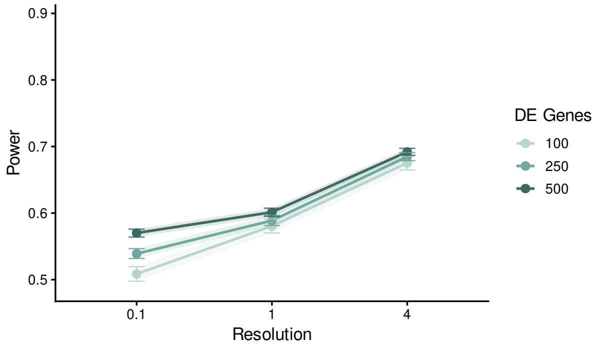


(c) Small clusters, $k = 20$

Figure S-35: **Power for differently sized non-aligned DE neighborhoods** Each plot shows power (y-axis) for increasingly higher resolutions (x-axis). Different number of DE genes are shown as differently colored lines. These results are using the simulation where the DE regions are not aligned to celltype, and thus are chosen by $K-$means clustering of cells with varying choice of the number of clusters $k$ for different genes ($k = 3$, $k = 10$, or $k = 20$), see Methods. In these plots, we stratify the non-null hypotheses according to the choice of $k$ used for the corresponding DE region, with larger $k$ giving generally smaller sized $N_g$. The result is averaged across simulations with different phenotype removal methods. For all plots, $N_g$ were chosen using Harmony sample-correction of null genes.

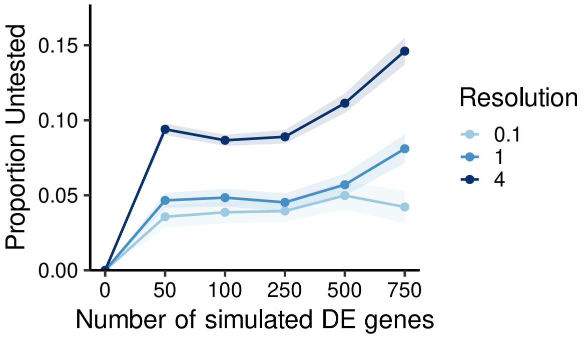


(a) Clusters dropped

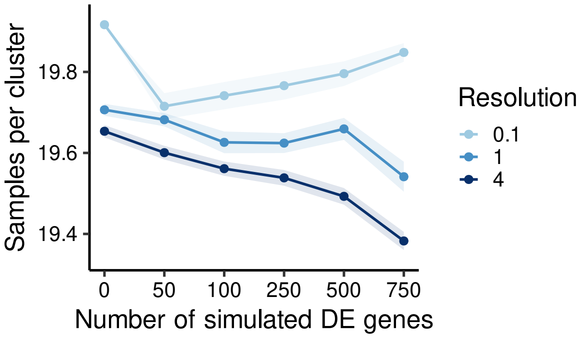


(b) Average Sample Count Per Cluster

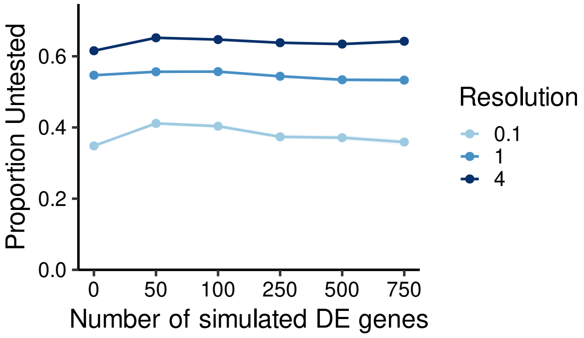


(c) Null Hypotheses dropped

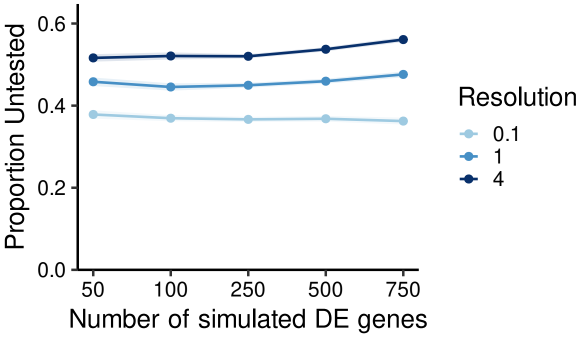


(d) Simulated Hypotheses dropped

Figure S-36: **Levels of filtering with varying resolutions and number of DE genes for DE regions non-aligned with celltype** Plots show measures of filtering averaged across simulations (y-axis) with growing number of DE genes (x-axis); different resolutions are shown by differently colored lines. The different filtering metrics calculated per simulation and then averaged across simulations are:(a) the proportion of dropped clusters, (b) the total number of samples kept in a tested cluster (out of a possible 20), averaged across tested clusters and then averaged across simulations (c) the proportion of null hypotheses dropped and (d) the proportion of simulated hypotheses dropped. We show results here for simulations that used Harmony for phenotype removal as well as for sample correction of null embeddings for $N_g$. Resolution ordering are similar for other phenotype removal methods for all filtering metrics.
