## Supplemental Text for "“Double-dipping” in testing for phenotypic differences with scRNA-Seq"

### Supplementary Text

### S1 Description of Simulations

We evaluate methods using multi-sample single-cell RNA-seq data simulated with known ground-truth differential expression (DE). The simulation design adapts the augmented data strategy of [1] to our cluster-based DE setting (Section 3.1). We start from real scRNA-seq counts in which no DE is expected. Samples are then randomly assigned to two phenotypic groups. The retained real genes form the null gene set $\mathcal{G}_{\text{null}}$; additional genes are simulated with expression that depends on phenotype only within a gene-specific DE neighborhood $N_g$ that will constitute the set of DE genes $\mathcal{G}_{\text{DE}}$ (see S1.4). The augmented count matrix–combining the data from the gene sets $\mathcal{G}_{\text{null}}$ and $\mathcal{G}_{\text{DE}}$–is the input to downstream clustering and DE analyses.

#### S1.1 Data and pre-processing

We start from preprocessed single-cell data (count matrix and cell-level metadata) derived from the Pelka *et al.* colorectal-cancer atlas [22]. The full atlas includes tissues from mismatch-repair-deficient (MMRd), mismatch-repair-proficient (MMRp), and normal samples across dozens of patients [22]. For simulation, we use a subset restricted to normal tissue: ∼59,800 cells from $n = 20$ samples (median ∼3,000 cells per sample), spanning six annotated cell-types (epithelial, B, T/NK/ILC, plasma, myeloid, and stromal). Because these cells come from a single biological context (normal) and phenotypes will be assigned at random, observed genes the Pelka dataset are treated as null for DE with respect to the assigned phenotype. We retain the top $G_{null} = 5{,}000$ genes by variance of log-normalized expression, which constitute our set of $\mathcal{G}_{\text{null}}$.

#### S1.2 Assigned phenotypes

To create a two-group comparison, we assign each sample $j$ independently to a control or treatment phenotype at random. All cells from sample $j$ inherit that assignment. This preserves within-sample correlation for null genes, treating the sample as the experimental unit (as in pseudobulk DE; Section 3.1).

We write $z_j \in \{\text{ctrl}, \text{trt}\}$ for the assigned phenotype of sample $j$ and $z_{i,j} = z_j$ for cell $c_{i,j}$.

#### S1.3 Choice of DE neighborhoods

For each simulated gene $g \in \mathcal{G}_{\text{DE}}$ we specify a DE neighborhood $N_g$ (Section 3.1): the set of cells in which the gene expression differs based on the assigned phenotype.

Neighborhoods are defined by clustering cells in a low-dimensional embedding computed from $\mathcal{G}_{\text{null}}$ only; neither the embedding nor the clustering uses assigned phenotypes.

##### S1.3.1 Low-dimensional embeddings

Cells are first embedded in $\mathbb{R}^d$ ($d = 50$) using PCA on scaled $\mathcal{G}_{\text{null}}$ expression. We consider three sample-integration strategies applied to this PCA space before clustering:

- **No correction:** PCA coordinates used directly.
- **Harmony:** Harmony correction [12] with batch variable chosen to be the sample.
- **fastMNN:** Mutual nearest-neighbor correction via `batchelor::fastMNN` [7], with batch defined by the sample.

Using Harmony and fastMNN reduce the risk sample-driven variations in embeddings, and thus of choosing DE regions $N_g$ which demonstrate sample bias.

##### S1.3.2 Candidate neighborhoods and sampling $N_g$

Let $\mathcal{N}$ denote the set of clusters obtained from a clustering of the embedded cells, from which an $N_g$ is chosen for each simulated $g \in \mathcal{G}_{\text{DE}}$. We use two approaches to construct $\mathcal{N}$:

- **$K$-means:** For each $K \in \{3, 10, 20\}$, run $K$-means on the embedding and record cluster labels. The candidate set $\mathcal{N}$ is the union of all clusters across these three partitions. For gene $g$, draw $k_g$ uniformly from $\{3, 10, 20\}$, use the corresponding partition, and set $N_g$ equal to a cluster chosen uniformly at random from that partition. For this construction, the average $\log_2$ fold change $|\beta_g^{\text{DE}}|$ (see below) is tied to $k_g$. These are set to 1.5, 3, and 4.5 for $k_g = 3$, 10, and 20. The differences in choice of $|\beta_g^{\text{DE}}|$ is try to equalize the power between the generally larger and smaller clusters expected for the different partitions from the $K$-means clusters.

- **Leiden:** Run Leiden clustering at resolution 0.1; $\mathcal{N}$ is the set the resulting clusters. For each $g$, choose $N_g$ uniformly from $\mathcal{N}$. For this construction, the average $\log_2$ fold change $|\beta_g^{\text{DE}}|$ is 3. Note this approach will closely match the clustering approach applied to the simulated data .

Genes in $G_{\text{null}}$ have $N_g = \emptyset$ by construction.

#### S1.4 Generative model for simulated counts

For each $g \in G_{\text{DE}}$ we generate counts $x_{i,j,g}$ from a negative-binomial model with log-linear mean, following [1]. Let $s(c_{i,j}) = j$ denote the sample of cell $c_{i,j}$, and let $\text{sf}_{i,j}$ be the cell size factor (library size divided by the median library size across cells). The log linear mean is given by:

$$\eta_{i,j,g} = \beta_g^{(0)} + \beta_g^{\text{DE}}\, \mathbf{1}[c_{i,j} \in N_g]\, \mathbf{1}[z_j = \text{trt}] + \beta_j^{(s)}, \tag{4}$$

where $\beta_g^{(0)}$ is a gene-specific baseline on the $\log_2$ scale, drawn uniformly on $[-7, 3]$; $\beta_j^{(s)} \overset{\text{i.i.d.}}{\sim} \mathcal{N}(0, \sigma^2_{\text{samp}})$ is a sample-level random effect ($\sigma^2_{\text{samp}} = 0.01$); and $\beta_g^{\text{DE}}$ is the treatment effect in $\log_2$ space for treatment cells in $N_g$ (control cells contribute 0 regardless of $N_g$ membership). Counts are then

$$x_{i,j,g} \sim \text{NB}(\mu_{i,j,g} = 2^{\eta_{i,j,g}} \cdot \text{sf}_{i,j},\ \phi)\,, \tag{5}$$

with a common dispersion $\phi = 0.2$ for all $g \in G_{\text{DE}}$ . In Leiden-based simulation settings we hold $|\beta_g^{\text{DE}}|$ fixed across genes (e.g. 3 or 1.5); in the $K$-means hierarchy we set $|\beta_g^{\text{DE}}|$ per gene depending on the choice of $k$ used to pick $N_g$, as described in S1.3.2.

##### S1.4.1 Structure of simulated DE

Notably from Equations 4 and 5, for a given gene $g$, a cell follows one of of two distributions. All cells assigned to treatment and in the DE neighborhood ($N_g \cap \{c_{i,j} | z_j = \text{trt}\}$) share a differentially expressed distribution. Conversely cells either assigned to control or outside of the DE neighborhood ($N_g^C \cup \{c_{i,j} | z_j = \text{ctrl}\}$) also share the same count distribution, removing sample and other cell type effects. In other words, for a given gene $g$, expected expression is the same for (i) control cells in $N_g$, (ii) control cells outside $N_g$, and (iii) treatment cells outside $N_g$. Only treatment cells in $N_g$ differ, by a gene-specific $\log_2$ fold change $\beta_g^{\text{DE}}$ whose sign is drawn uniformly at random.

### S2 Evaluation of testing of candidate neighborhoods

#### S2.1 Determining non-null hypotheses

**The null hypothesis for DE testing** The null hypothesis for found or candidate cluster $\hat{C}_k$ is generally the standard test, $\mu_{g,k,A} = \mu_{g,k,B}$, i.e. whether the mean gene expression of the two phenotypic groups $A$ and $B$ are the same for the set of cells in $\hat{C}_{g,k}$. For our simulation setup this is equivalent to

$$H_{0,g,k}^{DE} : |\hat{C}_{g,k} \cap N_{g,\ell}| = 0, \forall \ell.$$

Indeed, if $g \in \mathcal{G}_{\text{null}}$ this follows from the facts that $N_g = \emptyset$ and that samples from a same condition are assumed to be coming from a same distribution and randomly assigned to treatment or control. For $g \in \mathcal{G}_{\text{DE}}$ this is true by construction: cells in $(N_g \cap \{c_{i,j}|z_j = \text{trt}\})$ share a differentially expressed distribution and cells in $N_g^C \cup \{c_{i,j}|z_j = \text{ctrl}\}$ also share a different count distribution with a different mean. Hence any amount of overlap of $\hat{C}_k$ with $N_g$ results in $H_{0,g,k}$ being false (though small amounts of overlap will mean low power for detection).

**Requiring minimal overlap** In evaluating the performance of the DE tests, this definition can be overly harsh in practice. For example, extremely small amounts of overlap are unlikely to be detectable by any method, resulting in a low estimation of power. This could lead to poor concordance with the biological meaning, where small amounts of overlap of cells with differing mean would not be considered to be biologically meaningful as differentially expressed. This is a problem common in settings that are both detecting regions and testing for differences, such as chromosome segmentation analysis [11]. It can also be viewed as tolerance for a small amount of contamination or error in the data distribution specified by the null hypothesis.

Thus we may want to consider the amount of cluster overlap in determining which hypotheses are null or non-null, with a definition

$$H^{ov}_{0,g,k} : |\hat{C}_{g,k} \cap N_{g,\ell}| \text{ is sufficiently small } \forall \ell.$$

**Defining sufficiently small** To allow for tolerance in the amount of overlap or contamination, we need to define what is meant by "sufficient" overlap of a cluster $\hat{C}_k$ with a neighborhood $N_{g,\ell}$.

We first define the overlap between a considered neighborhood $\hat{C}_k$ and true neighborhood $N_g$ as

$$O_{g,k} = |\hat{C}_k \cap N_g|.$$

and define the percentage overlap, $\rho_{g,k,\ell}$, in two ways:

1. Union Proportion
$$\rho_{g,k} = \frac{O_{g,k}}{|\hat{C}_k \cup N_g|}$$
This would penalize large discrepancies between the cluster and neighborhood definitions.

2. Cluster Proportion
$$\rho_{g,k} = \frac{O_{g,k}}{|\hat{C}_k|}$$
This would penalize having diluted clusters (i.e. including cells from different neighborhoods in a single clusters)

The Union Proportion was used for testing the stability of the FDR control against small overlaps vs the strict no overlap definition. This permitted the definition of small to be relative to both $N_g$ and $\hat{C}_k$. The Cluster Proportion was used for the Power analysis since that is more directly compared to the size of $\hat{C}_k$ which is the basis for the pseudo-bulk counts that are passed for the testing.

In both cases, "sufficient" overlap of a cluster $\hat{C}_k$ with a neighborhood $N_g$ can be determined by the value of $\rho_{g,k}$ against a chosen threshold $b_{ov}$, yielding:

$$H^{ov}_{0,g,k} : \rho_{g,k} \leq b_{ov}$$

#### S2.2 Untested candidate regions

While the above analysis pipeline defines the same set of clusters $\hat{C}_1, \ldots, \hat{C}_K$ as candidate DE neighborhoods for every gene, in practice not all clusters are tested on every gene. Filtering steps applied during pseudobulk differential expression analysis remove some sample–cluster combinations, some entire clusters, and some

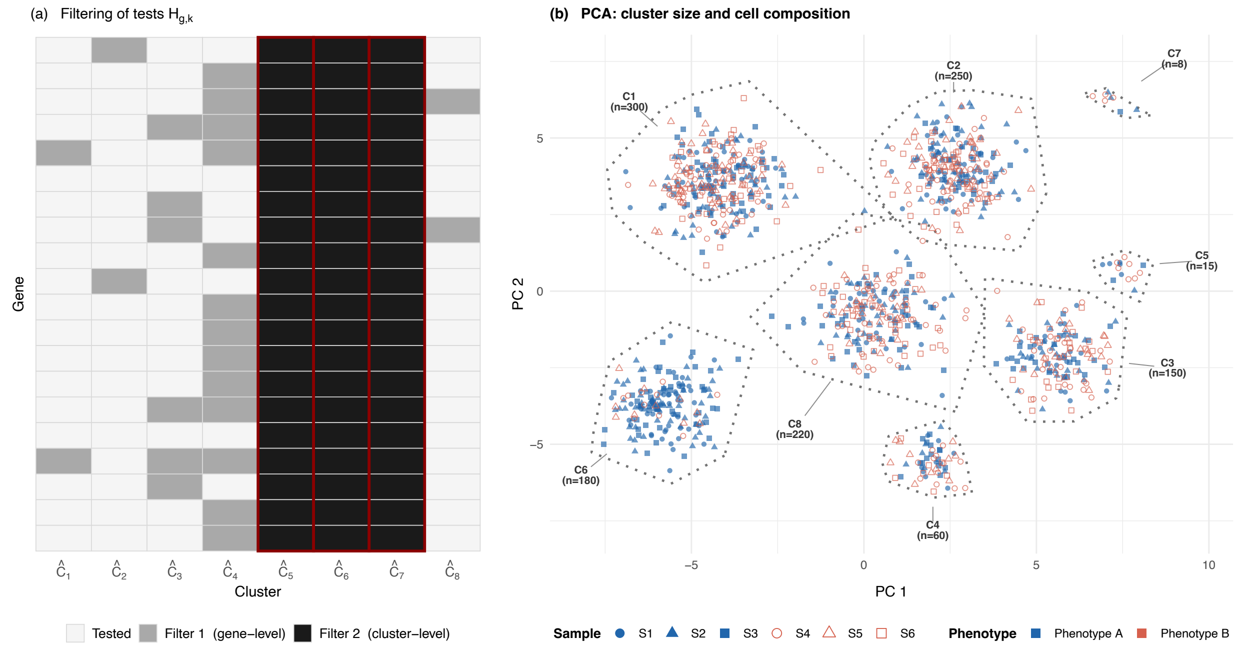


Supp. Fig. 1: **Illustration of Hypothesis Filtering** There are multiple types of filtering that can lead to a cluster/gene hypothesis to not be tested as outlined in more detail in S2.2. Notably, certain filters lead to a specific hypothesis not being tested, often due to low-expression genes, while other filters lead to all hypotheses associated with a given cluster to be dropped, often due to phenotype or sample imbalance.

gene–cluster pairs before hypothesis tests are run. As a result, the number of tests performed for gene $g$, which we denote $K_g$, satisfies $K_g \leq K$ and can vary across genes (see Supp. Fig. 1)

Untested hypotheses are never rejected, so they do not influence FDR calculations. For power calculations, we only consider tested hypotheses, meaning that untested gene-cluster pairs are not included in the denominator even if the cluster overlaps with $N_g$

##### S2.2.1 Filters applied within each cluster

In our case, differential expression was performed with `muscat::pbDS` using `method = "edgeR"` [23] with default filtering settings with the following steps are applied *in order* by muscat/edgeR.

1. **Minimum cells per sample–cluster combination.** Any sample with less than 10 cells for that sample-cluster pair is excluded from the pseudo-bulk analysis of cluster $k$.

2. **Low-library pseudobulk sample removal** This filtering step is done for each cluster. The sum of pseudobulk counts across genes is computed for each sample within the cluster. Outlier low count samples are removed from that cluster, defined as being below the median by more than three median absolute deviations on the log-count scale. All genes in the sample-cluster pair are then removed for testing.

3. **Design specific cluster exclusion** A cluster is dropped entirely if any of the following hold:

   - any phenotype group only has 0 or 1 sample;
   - every sample has a distinct covariate pattern.
   - the model matrix is rank-deficient;

   For our purpose the first is particularly relevant when clusters are dominated by one phenotype. In the extreme case, a cluster $\hat{C}_k$ may contain cells from only one phenotype and is necessarily excluded.

| $H_{0,g,k}$ | | $\tilde{p}_{g,k} > \alpha$ | $\tilde{p}_{g,k} \leq \alpha$ | Unable to test |
|---|---|---|---|---|
| False | $\rho_{g,k} > b_{ov}$ | FN | TP | $\text{FN}_{\text{untested}}$ |
| True | $\rho_{g,k} \leq b_{ov}$ | TN | FP | $\text{TN}_{\text{untested}}$ |

Supp. Table 1: Defining true positives and false positives based on overlap cutoff.

4. **Removal of zero-count genes** Genes with zero total pseudobulk count across those samples are removed.
5. **gene expression filtering (edgeR filter).** A gene is retained only if it reaches sufficient count-based expression thresholds in enough samples to support estimation in the two-group design: at least 10 counts per million in a sufficient number of samples, at least, and at least 15 total counts across samples with expression in at least 70% of the samples in the smallest phenotype group for large groups sizes.
6. **Removal of samples with no expression among retained genes.** Any remaining sample with non-zero gene counts are dropped, and the design matrix is updated accordingly.

Our pipeline pools raw $p$-values across all tested gene–cluster pairs and applies a single Benjamini–Hochberg adjustment across the full family of tests (`adjust_all_pvals`); gene–cluster pairs that were filtered out are not assigned $p$-values, are not included in the p-value adjustment, and are treated as untested.

##### S2.2.2 Relation of Filters to Phenotype effects

The filters above map onto two qualitatively different mechanisms, namely gene-level versus cluster/sample level filtering. However, it is important to note that, for strong enough phenotype effects, clusters $|\hat{C}_k|$ dominated by specific subtype will likely trigger both types of filters due to the small number of cells of one of the phenotypes. A similar argument can be made for clusters that are inherently smaller in size.

1. **Gene-level filtering (steps 4–5).** Standard bulk RNA-seq practice removes genes whose pseudobulk counts in cluster $k$ are too low or too sparsely detected for reliable inference. Because $X_{g,k,j}$ is the sum of single-cell counts in $\hat{C}_k \cap$ sample $j$, genes in small or low-RNA clusters are more likely to fail `filterByExpr`. Thus, even though every gene is *eligible* in principle for testing in each of the $K$ clusters, the number of tested pairs $K_g$ varies by gene.
2. **Sample- and cluster-level filtering (steps 1–3, and step 6).** These filters determine which pseudobulk samples contribute to cluster $k$, and whether cluster $k$ is tested at all. Step 1 removes samples with fewer than ten cells in $\hat{C}_k$. Step 2 removes samples with aberrantly low total pseudobulk expression in that cluster. Step 3 removes the entire cluster when fewer than two samples per phenotype remain or the design is not estimable—including the practically important case of phenotype-skewed clusters created by post-clustering selection. When a cluster is removed at step 3, *all* genes are untested in that cluster; when only steps 1–2 apply, different clusters $k$ are analyzed with different numbers of pseudobulk replicates.

When $|\hat{C}_k|$ is small, both mechanisms could remove many hypotheses: the cluster may fail step 3 outright, and genes in the cluster are more likely to fail the expression filters in steps 4–5. (However, we should note that there is an order in the filtering, and so hypothesis are first removed based on sample and cluster-level filtering so the filtering may never get to steps 4–5).

#### S2.3 Definition of TP/FP

With our hypotheses definitions and an understanding of the existence of tested and untested hypotheses, we can determine determine true/false positives and negatives based on our adjusted p-value FDR cutoff $\alpha$ (Supp. Table 1). For our analyses, $\alpha = 0.1$ unless otherwise noted.

#### S3 Strong DE signal drives FDR decrease

In Section 2.3, we discussed the behavior of type I error for different resolution, and noted that the FDR trends in our simulations do not always match the type I error trends, since FDR correction methods, like Benjamini-Hochberg, involve the joint distribution of both the true positives and the false positives. Thus, at high resolutions we see a large drop in FDR with the introduction of DE signal which continues to drop as we add more DE genes (Figures 6c and 6d). As we will detail more in this section, this is due to our simulation having introduced true positives with large signal with a p-value distribution which is highly separated from the distribution of that of the null hypotheses.

Upon inspection of type I error (Figure S-27), higher resolutions result in worse type 1 error control at the levels of type I error for which BH-correction would control for the multiple testing. For 0 DE genes, this lack of type I error control consistently translates into higher FDR at 0 DE genes (which in the case of 0 DE genes is equivalent to family-wise error rate). As the number of included simulated DE genes increases, type 1 error remains similarly poorly controlled, yet FDR, based on the BH corrected p-values, decreases (Figures S-27, 6d). The inclusion of DE genes changes the choice of BH cutoff to include a disproportionally larger portion of non-null hypotheses rather than null hypotheses. In the case of our simulation, the p-value distribution of null hypotheses stay relatively stable (as supported by the consistent type 1 error), but the non-null hypotheses are highly skewed towards 0, particularly for large log-fold changes. Indeed, using log fold change of 3, p-values from nun-null hypotheses have have their mass consistently between 1e-16 and 1e-9 while non-null hypotheses between 1e-5 and 1e-3. This results in a clear separation between the null and non-null hypotheses, thus leading the BH method to pick a correction level that excludes more null genes than when there were no DE genes present. In the setting of lower log-fold change, we see the drop of FDR diminish, as expected, (Figure S-32) and this separation effect becomes less pronounced - null hypotheses maintain similar ranges while non-null hypotheses now have half their mass between 1e-7 and 1e-11.

#### S4 Datasets Analyzed

In addition to the simulation study (Supplementary Section S2), we applied the cluster-based DE pipeline to four published scRNA-seq datasets spanning distinct tissues and experimental designs (Table 2). The Pelka, Liu and Melms datasets were initially pre-processed following to the Gloscope procedure [[30]]. The Liu dataset was directly taken from the source in the paper [15]. All four datasets were then analyzed with the shared workflow described in Section 3.4.

Supp. Table 2: **Summary of real-data benchmarks**. “Sample unit” is the sample identifier used for muscat aggregation and edgeR testing.

| Dataset | Reference | Phenotype contrast | Cells | Sample $n$ | Sample Identifier |
|---|---|---|---|---|---|
| Kang PBMC | [10] | IFN-$\beta$ stim. vs. control | 24,673 | 16 | donor×condition |
| Pelka CRC | [22] | MMRd tumor vs. adjacent normal | 230,334 | 118 | sequencing library |
| Liu rash | [15] | Psoriasis vs. healthy control | 68,835 | 15 | patient |
| Melms COVID lung | [20] | COVID-19 vs. control | 116,313 | 26 | patient |

##### S4.1 Kang *et al.* multiplexed PBMC dataset

Kang *et al.* performed droplet scRNA-seq on multiplexed peripheral blood mononuclear cell (PBMC) pools from eight individuals [10]. Each donor contributed cells to both an untreated control condition and a 6-hour IFN-$\beta$ stimulation condition; sample identity was resolved with demuxlet. We used the processed Seurat object `Kang2018.rds` (15,706 genes; 24,673 singlet cells after doublet exclusion).

This is a **paired** design: every donor appears in both phenotypes. Samples were defined as donor×condition, yielding 16 samples.

#### S4.2 Pelka *et al.* colorectal cancer atlas

Pelka *et al.* profiled colorectal tumors and adjacent normal tissue from 62 patients using 10x Genomics scRNA-seq [22]. The published atlas includes mismatch-repair-deficient (MMRd) tumors, mismatch-repair-proficient (MMRp) tumors, and adjacent normal specimens.

We retained Normal cells and cells with MMRd, yielding 117,470 MMRd tumor cells (66 sequencing libraries, 34 donors) and 112,864 adjacent-normal cells (52 libraries, 36 donors); MMRp tumors were excluded.

**Relation to simulations.** The semi-synthetic simulations (S2.1) used a different subset Pelka subset: normal tissue only, removing donors with abnormally low or high number of cells (retaining $\sim$59,800 cells from 20 samples), chosen so that observed genes could be treated as null before random phenotype assignment.

#### S4.3 Liu *et al.* inflammatory skin disease cohort

Liu *et al.* profiled $CD45^+$ immune cells from 31 skin biopsies (psoriasis, atopic dermatitis, clinically indeterminate rashes, lichen planus, bullous pemphigoid, and healthy controls) using 10x Genomics scRNA-seq [15](145,810 singlet cells; 22,253 genes).

For the benchmark we restricted to psoriasis vulgaris versus healthy skin control (7 controls, 8 psoriasis patients; 68,835 cells). Each patient contributes one biopsy and maps to a single phenotype so samples are indexed by patient (15 donors).

#### S4.4 Melms *et al.* lethal COVID-19 lung atlas

Melms *et al.* generated a single-nucleus RNA-seq atlas of lung tissue from 19 individuals who died of COVID-19 (rapid autopsy) and seven pre-pandemic control individuals [20] (116,313 nuclei; 29,925 genes after removing genes detected in fewer than ten nuclei).

The contrast is lethal COVID-19 versus pre-pandemic control with 19 COVID-19 and 7 control samples.
